# Lifespan Remodeling of Cerebral Biomechanical Network Organization Reveals Coordinated Maturation and Aging-Related Convergence

**DOI:** 10.64898/2026.11.17.767305

**Authors:** Jakob Ludwig, Biru Huang, Tom Meyer, Joanes Grandjean, Guillaume Flé, Anna S. Morr, Rafaela V. Silva, Melanie Estrella, Carmen Infante-Duarte, Ingolf Sack, Jing Guo

## Abstract

Brain biomechanical connectivity (BC) captures shared regional viscoelastic properties that may support functional interactions. Although functional and structural brain networks are well characterized, the systems-level organization of brain biomechanics remains unknown. Using in-vivo magnetic resonance elastography (MRE) and community detection, we established BC and its network organization across the mouse lifespan, spanning seven stages from early adolescence (4 weeks) to old age (72 weeks). BC was benchmarked against established functional and structural connectivity, revealing shared connectivity features. Biomechanical network organization showed increasing segregation during adolescence, peaking in late adolescence, followed by progressive reintegration with aging. Biomechanical network dynamics defined by the pace of age-related change showed coordinated maturation during adolescence, regionally differentiated trajectories throughout adulthood, and partial reconvergence in aging. These findings establish a lifespan framework for cerebral biomechanical connectivity in mice and reveal age-dependent systems-level network reorganization that may underlie critical windows of brain maturation, aging, and disease vulnerability.

## Introduction

The brain is an intricate network comprising regions that are anatomically interconnected and interact functionally to process sensory information, facilitate neuronal processes, and control motor and vital functions. Each region maintains its distinct structure and neural and glial populations for functional specialization, while concurrently integrating into the global brain network for coherent perception and behavior[1, 2]. Using resting-state functional magnetic resonance imaging (rs-fMRI), the functional connectivity of brain networks and the functional synchronization of brain subregions have been extensively explored by the temporal co-variation of hemodynamic responses [3, 4]. These signals have revealed that the functional brain network is composed of segregated groups of well-connected communities[5]. Changes in functional communities were associated with various brain disorders such as sleep deprivation[6], epilepsy[7], and Alzheimer’s disease[8]. Furthermore, previous data revealed that functionally connected brain regions often, though not invariably, exhibit direct structural connectivity[9, 10].

Cerebral biomechanical properties, as quantified by magnetic resonance elastography (MRE)[11], have been shown to be sensitive to alterations in both structure and hemodynamics during physiological processes such as aging[12], hyperthermia[13], and hypoxia[14], as well as in pathologies including multiple sclerosis[15], Alzheimer’s disease[16], and Parkinson’s disease[17]. Therefore, the biomechanics-based brain connectivity could serve as an effective connectivity that combines anatomical connections (anatomical networks) and dynamic interactions (functional networks). The objective of this study was therefore to establish cerebral biomechanical connectivity (BC) using in vivo brain MRE and characterize its age-dependent network organization through community detection in a cerebral maturation and aging mouse model. Based on biomechanical similarities of brain subregions, we have examined the evolution and degeneration of the cerebral biomechanical network and map the trajectories of the cerebral segregation and integration over the course of a mouse lifespan. Unlike previous MRE studies of the biomechanical properties of maturing and aging brains, where pairwise and group comparisons were performed separately for each brain region[12, 18, 19], our study focused on identifying network communities on a systemic level that follow biomechanical organizational principles and distinguishing brain regions with high and low biomechanical specialization, thereby providing new insights into how the brain reconfigures and recalibrates itself at different phases in life, which could reveal the brain’s flexibility, adaptability, and resilience across the lifespan. The proposed cerebral biomechanical connectivity and its network organization could serve as age-dependent reference for the detection and monitoring of neurological disorders and injuries.

## Results

### Changes of cerebral biomechanical properties over age

Representative brain elastographic atlases are shown in one central slice for all age groups (EAdol: early adolescence; MAdol: mid-adolescence; LAdol: late adolescence; EAdult, early adulthood; MAdult: mature adulthood; LAdult: late adulthood; Old: old age) (Figure 1). Based on the biomechanical contrast, the brain regions are clearly visible and manifest age-dependent variations. Global brain stiffness, assessed by the group-averaged shear wave speed (*c*), varied significantly with age (p < 0.001). The greatest reduction in *c* occurred between MAdol and Old age (−7.5 ± 0.1%, p = 0.01; Supplemental Figure SF1A). Phase angle *φ*, surrogate marker for viscosity, increased progressively over age (p < 0.001) with the most prominent change between EAdult and Old age (26.7 ± 0.1%, p = 0.02, Supplemental Figure SF1B). For each brain region, the group-averaged biomechanical properties are illustrated by heatmaps (Figure 2A), which depict relative changes in *c* and *φ* at all ages compared to EAdol values. As shown in Figure 2A, in comparison to EAdol, 77% of all 122 subregions demonstrated an increase in stiffness (higher *c*) at MAdol. Comparing to EAdol, 51% of all 122 subregions showed signs of softening already at EAdult, particularly in isocortex (57% subregions) and hypothalamus (83% subregions). The Cornu Ammonis (CA) subfields and Dentate Gyrus (DG) of the hippocampal formation exhibited distinct age-related trajectories (p < 0.001), with tissue stiffness remaining elevated relative to EAdol till MAdult (CA: 18.5 ± 0.6%, p = 0.02; DG: 23.4 ± 0.4%, p = 0.004, Supplemental Figure SF1C-D). In contrast, the hypothalamus demonstrated the earliest onset of age-related softening, already evident in LAdol, and showed the most pronounced decrease in stiffness with advancing age (MAdol vs. Old age −26.0 ± 0.4%, p < 0.001, Supplemental Figure SF1E). Compared to EAdol, a significant increase in viscosity was observed in 80% of brain subregions starting from LAdult. A marked decline in viscosity was evident in the thalamus as early as LAdol (EAdol vs. LAdol, −17.2 ± 0.1%, p = 0.002, Supplemental Figure SF1F). The results of the pairwise comparison over the age phases are shown in Figure 2B. In general, more significant pairs were identified for viscosity than for stiffness across age phases and regions (306 vs. 254), showing however greater variation in viscosity across time points and regions. As shown in the outermost section of the circular plot, there is no biomechanical difference between LAdult and Old age across regions. Moreover, most of the significant pairs were identified by comparing LAdult or Old age with any other age phase, as shown by the outer area of each circular section in the plot. The complete set of 122 brain regions, together with their corresponding biomechanical properties and statistical analyses, is provided in Supplemental Table ST1.

**Figure 1.**
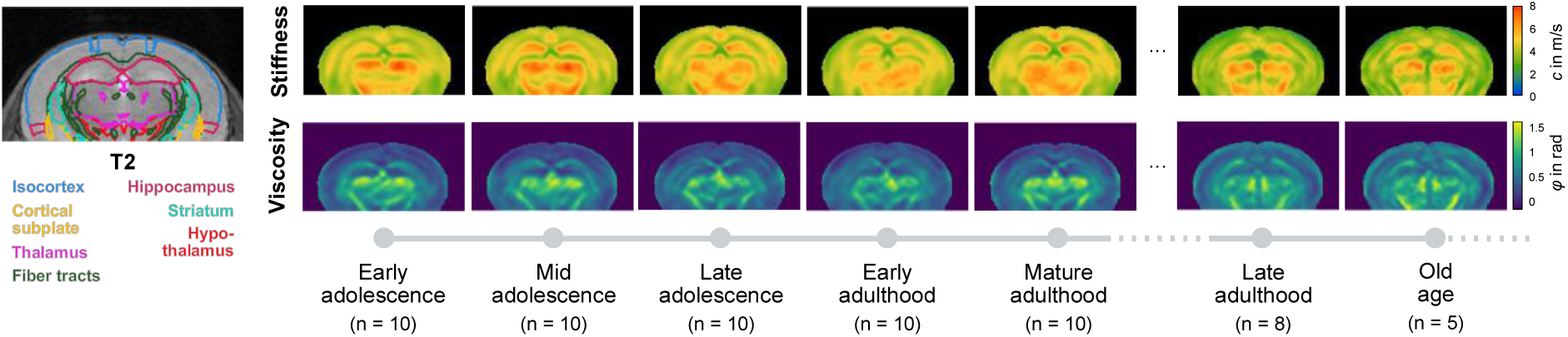
Atlas-registered, group-averaged MRE parameter maps of stiffness and viscosity are shown for a representative central brain slice across the lifespan. Regions of interest defined by the Allen Brain Atlas are overlaid in color on the corresponding T2-weighted image of the same slice. The age points are: early adolescence (4 weeks), mid-adolescence (6 weeks), late adolescence (8 weeks), early adulthood (10-12 weeks), mature adulthood (16-20 weeks), late adulthood (48 weeks), and old age (72 weeks).

**Figure 2.**
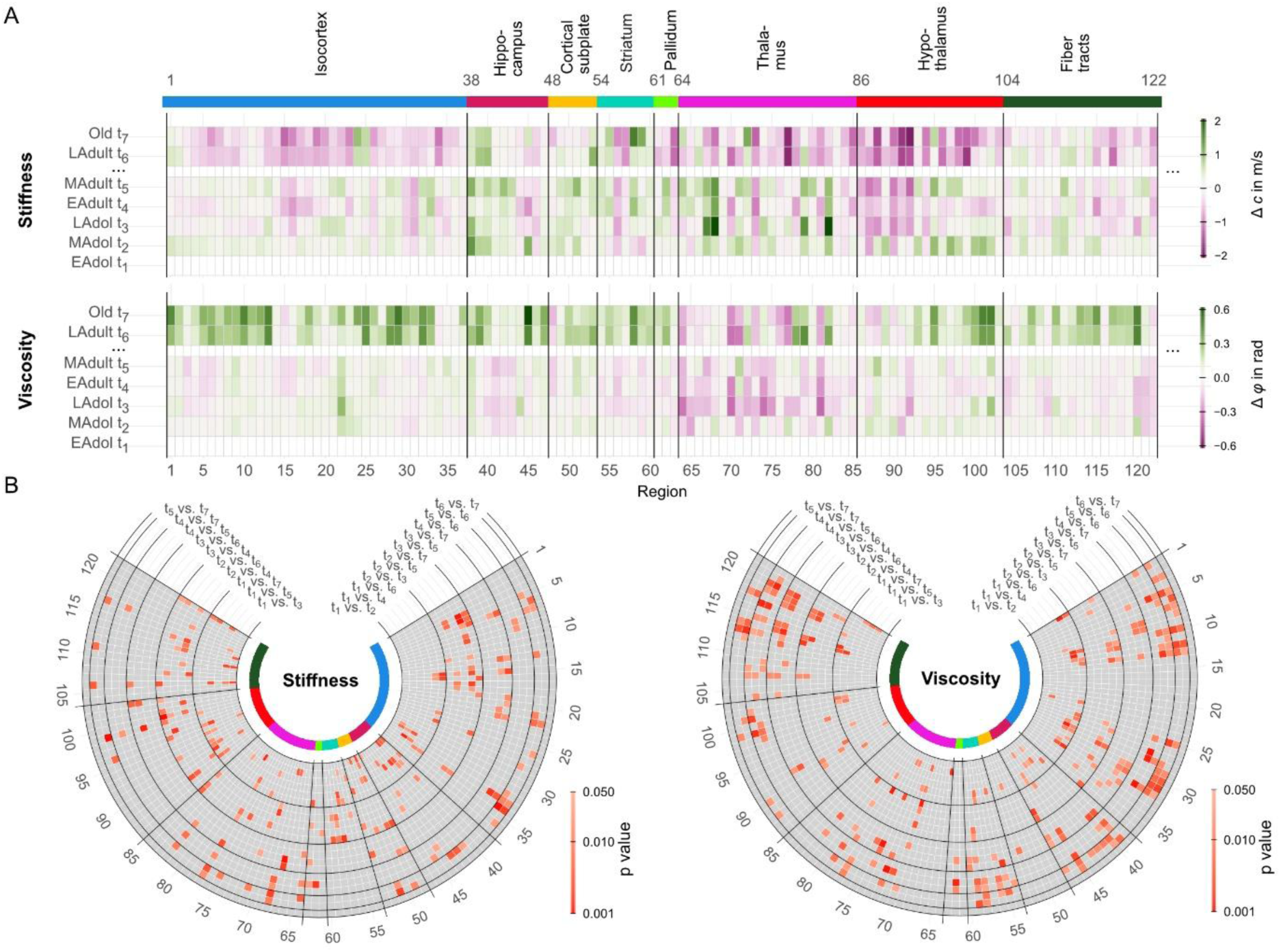
Changes in cerebral biomechanical properties across the lifespan. (A) Heatmaps illustrating changes in stiffness and viscosity across 122 subregions belonging to eight major brain regions, with each age group compared to values obtained during early adolescence. (B) Circular maps showing the results of pairwise comparisons between all age groups across the entire lifespan for both stiffness and viscosity. Red-highlighted cells indicate statistically significant differences between age-group pairs. The age groups are: early adolescence (4 weeks, EAdol, *t1*), mid-adolescence (6 weeks, MAdol, *t2*), late adolescence (8 weeks, LAdol, *t3*), early adulthood (10-12 weeks, EAdult, *t4*), mature adulthood (16-20 weeks, MAdult, *t5*), late adulthood (48 weeks, LAdult, *t6*), and old age (72 weeks, Old, *t7*).

### *Cerebral biomechanical connectivity (BC)* compared with functional and structural connectivity

Biomechanical connectivity (BC), based on age-specific biomechanical similarity (static BC, sBC) and age-dependent regional covariance (dynamic BC, dBC), was compared with resting-state fMRI-derived functional connectivity (FC) and viral tracing-derived structural connectivity (SC). The connectivity matrices are shown in Figure 3, methodological details are provided in the Methods section, and the corresponding parameter values are summarized in Supplemental Tables ST2 and ST3. Ipsilateral connections were stronger than contralateral connections in both SC (7.6% higher ipsilaterally) and FC (3.1% higher ipsilaterally). In contrast, ipsilateral and contralateral connection strengths were nearly identical across most BC matrices, with differences of less than 0.2% for both sBC and dBC. The exception was dBC based on tissue viscosity where ipsilateral connections were 1.8% stronger than contralateral connections.

**Figure 3.**
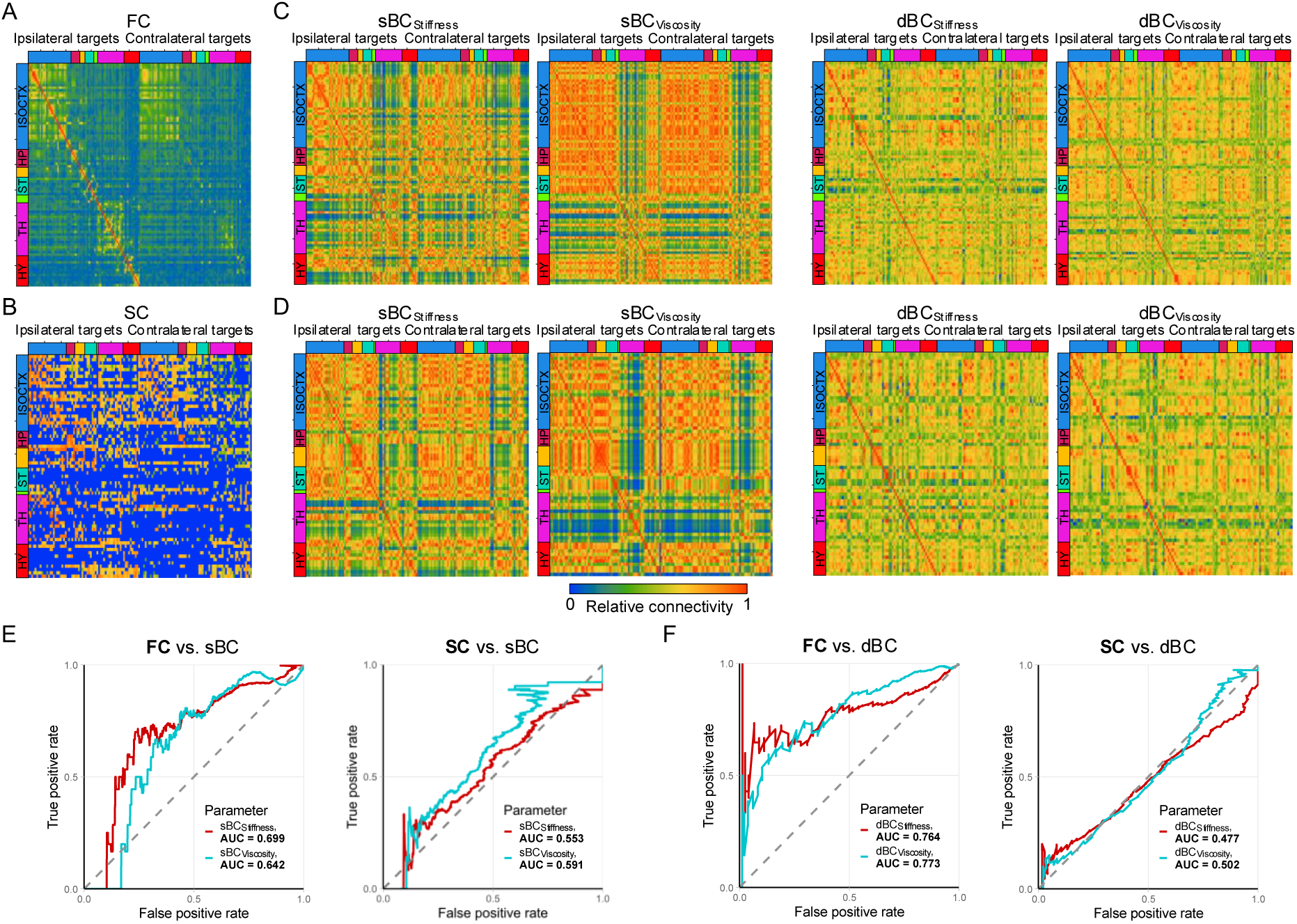
Comparison of biomechanical, functional, and structural connectivity. Matrices of cerebral (A) functional connectivity (FC), (B) structural connectivity (SC), and (C, D) biomechanical connectivity (BC) were constructed. BC was defined based on age-specific biomechanical similarity (static BC, sBC) or age-dependent interregional covariance across the lifespan (dynamic BC, dBC). For both sBC and dBC, two sets of stiffness- and viscosity-based matrices were constructed using datasets with different age and regional parcellations to match those of the available FC and SC datasets. Similarities between FC and BC (panels A vs. C) and between SC and BC (panels B vs. D) were quantified using the area under the curve (AUC; panels E-F).

Matrix comparisons revealed greater similarity between BC and FC than between BC and SC. Quantitative matrix-similarity analysis combining ipsilateral and contralateral connections yielded AUCs of 0.70 and 0.64 for sBC-FC comparisons based on stiffness and viscosity, respectively, and 0.55 and 0.59 for sBC-SC comparisons. For dBC, the corresponding AUCs were 0.76 and 0.77 for FC and 0.48 and 0.50 for SC, based on stiffness and viscosity, respectively (Figure 3). Although the overall similarity between BC and FC or SC across all brain regions was comparable between sBC and dBC, dBC exhibited more interregional similarities with both FC and SC, with a greater number of region pairs showing AUCs ≥ 0.6 (stiffness, dBC vs. sBC: 19 vs. 13; viscosity: dBC vs. sBC: 15 vs. 5. The corresponding region pairs are detailed in Supplemental Figure SF2.

### Cerebral biomechanical network organization over age

Leiden community detection, applied to the biomechanical properties of all regions across all age points, revealed a dynamic reorganization of community structure over the lifespan. Sankey plots in Figure 4 show the changes in network architecture and its reorganization with colors illustrating different anatomic regions. As seen in these Sankey plots, the number of detected communities increased from four in EAdol to six during LAdol for both stiffness (Figure 4A) and viscosity (Figure 4B), indicating increased biomechanical differentiation. With increasing aging, stiffness-based communities decreased to five only at Old age, whereas viscosity-based communities exhibited an earlier reduction, returning to four communities already by MAdult. Additionally, transitioning between communities and no visually stable communities evident over the whole age span. Temporal stability was observed in isocortex, with 46% of regions remaining within the same community from EAdol to LAdol, and in the thalamus, where 70% of regions maintained their community affiliation from EAdult to MAdult. Compared to stiffness, viscosity-based community stability exhibits a markedly different pattern: 32% of isocortex, 60% of hippocampal formation, 60% of cortical subplate, and 41% of hypothalamus remains stable across the entire age span. Notably, regions that transiently split into separate communities during MAdol subsequently reintegrate into single communities by EAdult. The observed changes in the number of communities over age that reflects biomechanical segregation and integration were further quantified using two global graph metrics: modularity and characteristic path length (CPL), which are plotted over time (Figure 5A). Modularity increased from EAdol to LAdol in both stiffness- (52.5 ± 21.7%) and viscosity-based (59.3 ± 10.5%) networks, indicating greater community segregation. Stiffness-based modularity then plateaued across the remaining age groups. In contrast, viscosity-based modularity declined after LAdol, reached a minimum in MAdult (−29.3 ± 4.7% compared to EAdol), and increased again to EAdol levels in LAdult and Old age. The trajectory of stiffness-based CPL paralleled those of modularity, showing a pronounced increase during adolescence, with a 31.1 ± 1.3% rise between EAdult and EAdol. In contrast, viscosity-based CPL exhibited a marked decline with aging, with an −17.9 ± 0.7% reduction from EAdol to Old age, mirroring the age-related decreases observed in modularity, based on viscosity. However, unlike modularity, viscosity-based CPL did not capture the initial increase observed during adolescence.

**Figure 4.**
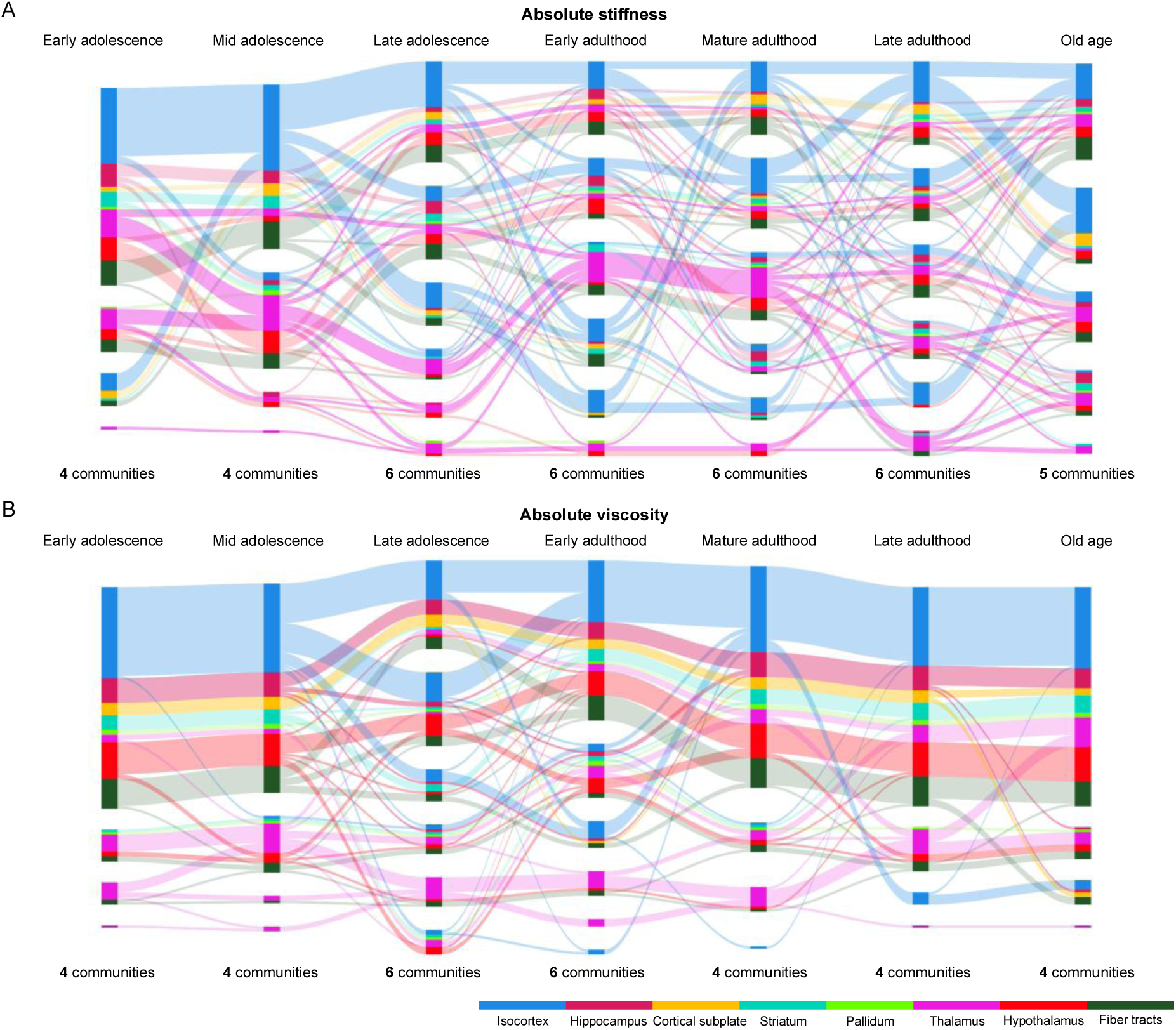
Sankey plots illustrating the results of the community analysis based on the absolute values of stiffness and viscosity across the lifespan, showing the number of detected communities and the trajectories of brain subregions between communities over age.

**Figure 5.**
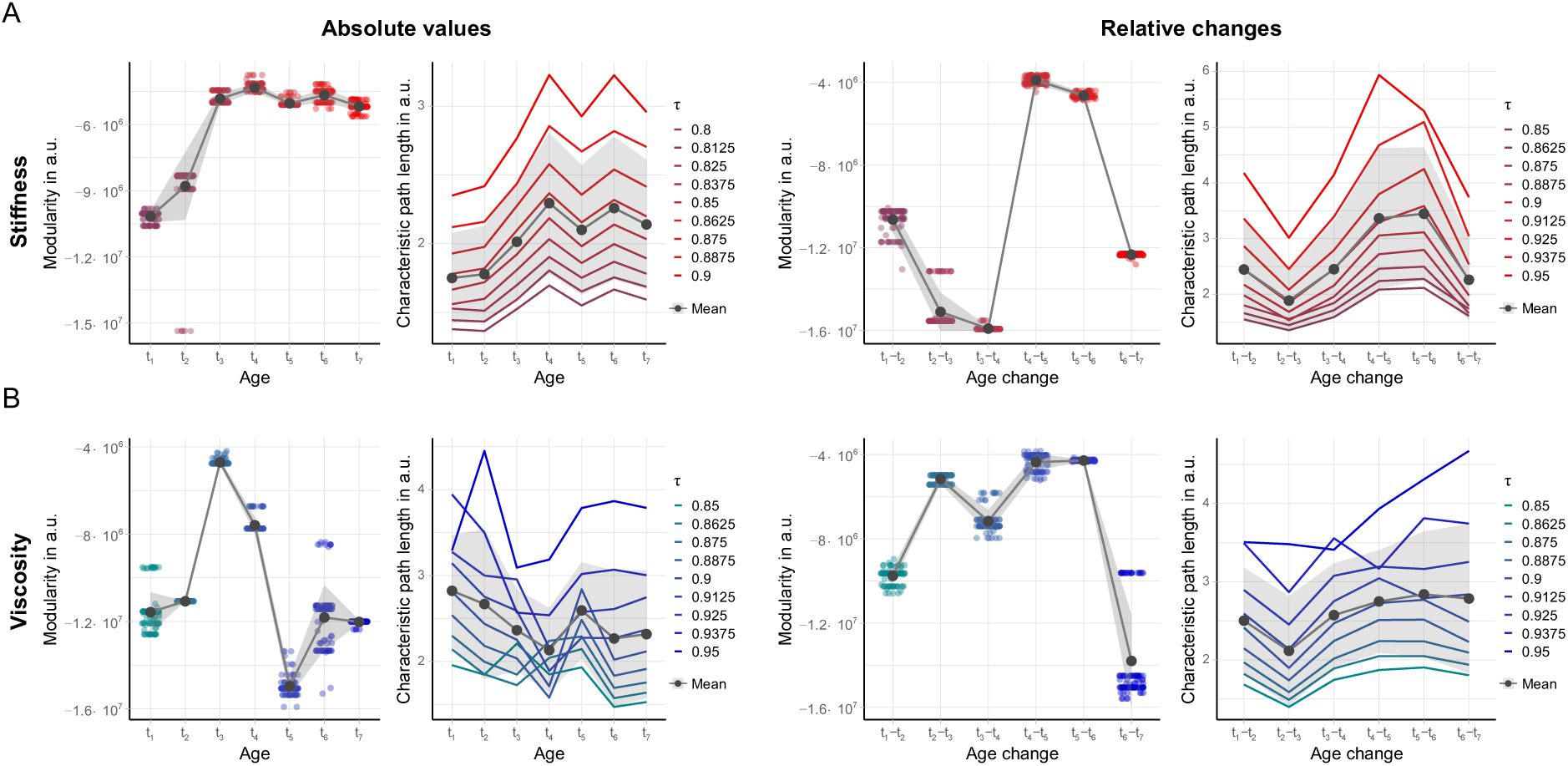
Quantitative graph metrics characterizing biomechanical network organization based on (A) the absolute values of stiffness and viscosity, as well as (B) the relative changes in these parameters across the lifespan. Gray shaded areas indicate standard deviations, calculated from 100 bootstrap iterations for modularity and across nine iterative *τ* values for characteristic path length. The age groups are: early adolescence (*t1*), mid-adolescence (*t2*), late adolescence (*t3*), early adulthood (*t4*), mature adulthood (*t5*), late adulthood (*t6*), and old age (*t7*).

Leiden community detection was applied to relative biomechanical properties across adjacent age groups to evaluate biomechanical connectivity associated with the pace of age-related change (Figure 6). Stiffness-based pace exhibited pronounced segregation from EAdult to MAdult, reflected by an increased number of communities (n = 6 vs. n = 4), higher modularity (−3.9×10^6^ ± 1.9×10^5^ vs. −1.6×10^7^ ± 9.0×10^4^, Figure 5A), relative to the LAdol-to-EAdult transition. This pattern gradually shifted toward greater integration from LAdult-to-Old with four communities and lower modularity of - 1.2×10^7^ ± 6.5×10^4^. In contrast, viscosity-based pace trajectories exhibited an earlier segregation, emerging during MAdol-to-LAdol (6 communities, modularity = −5.2×10^6^ ± 2.2×10^5^) and persisting through MAdult-to-LAdult (6 communities, modularity = - 4.3×10^6^ ± 3.3×10^4^) relative to the MAdol-to-EAdol transition (4 communities, modularity = −9.8×10^6^ ± 3.7×10^5^). Only during the transition from LAdult to Old age did viscosity changes become more integrated across the brain with four communities, modularity of −1.4×10^7^ ± 2.2×10^6^. This late-life integration in viscosity paralleled the stiffness-related pace pattern observed during the same age transition. The trajectory of stiffness-based CPL paralleled that of modularity, exhibiting a pronounced increase during adulthood, with a 37.5 ± 0.8% increase between EAdult-to-MAdult and EAdol-to-MAdol, as well as a significant decrease over aging (−7.5 ± 0.2% reduction between LAdult-to-Old and MAdult-to-LAdult). In contrast, viscosity-based CPL showed a gradual increase across age, except for an −15.4 ± 0.6% decrease between MAdol-to-LAdol and EAdol-to-MAdol. Community assignments based on relative changes in stiffness remained largely stable in the isocortex and fiber tracts, with 81% and 73% of subregions in the same community from EAdol to EAdult, respectively. In contrast, substantial temporal reconfiguration was observed throughout adulthood across all regions. Community assignments based on relative viscosity changes were highly unstable, with widespread transitions and no persistent community assignment across the age span.

**Figure 6.**
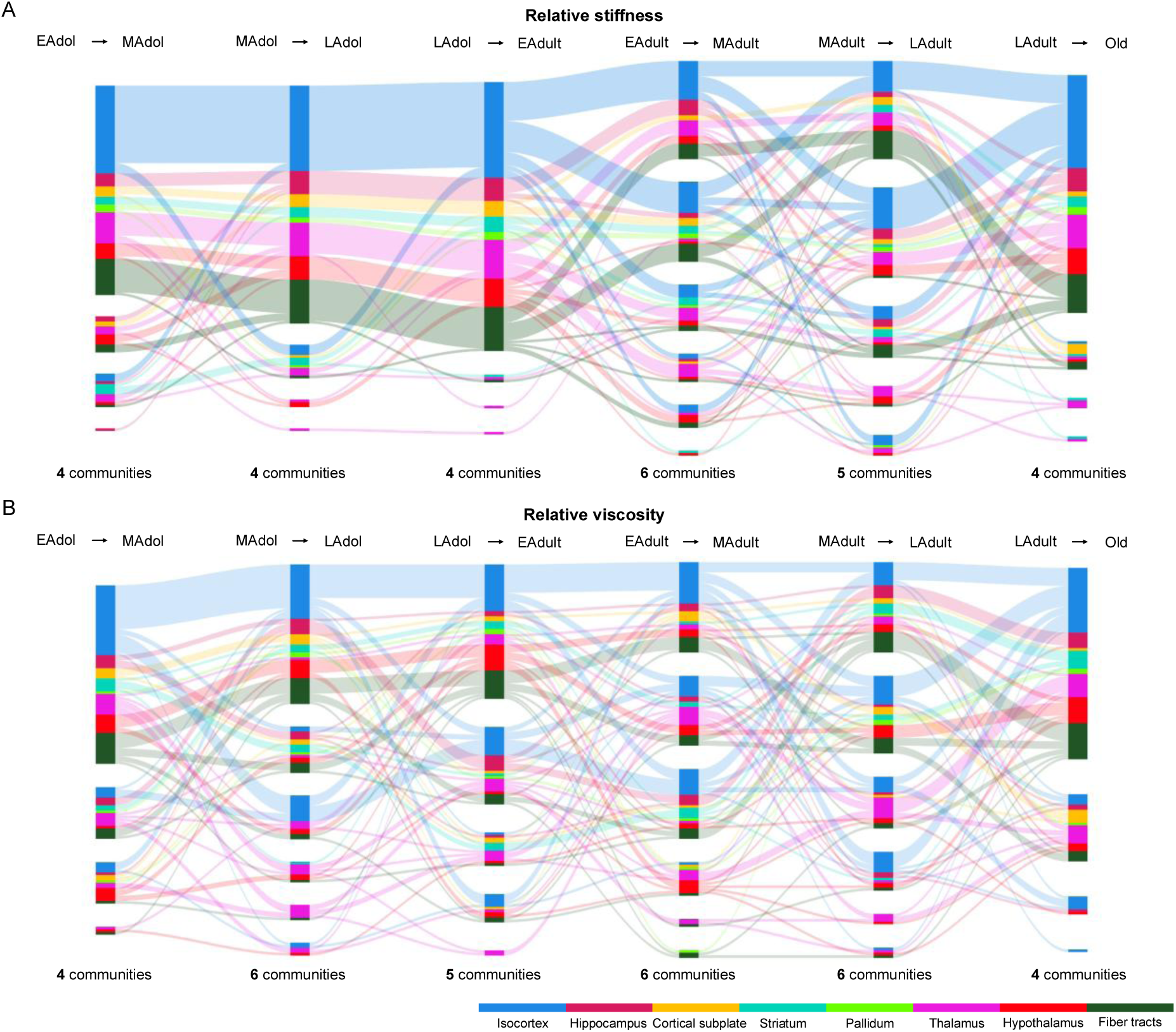
Sankey plots illustrating the results of the community analysis based on the relative changes in stiffness and viscosity (pacing) between consecutive age points across the lifespan, showing the number of detected communities and the trajectories of brain subregions between communities over time.

Color-coded regions and their corresponding community assignment over age are summarized in Supplemental Table ST4. Figure 5 and Table 1 summarize global graph metrics at individual age points and across age intervals, based on absolute biomechanical properties and their relative changes over time.

**Table 1.**
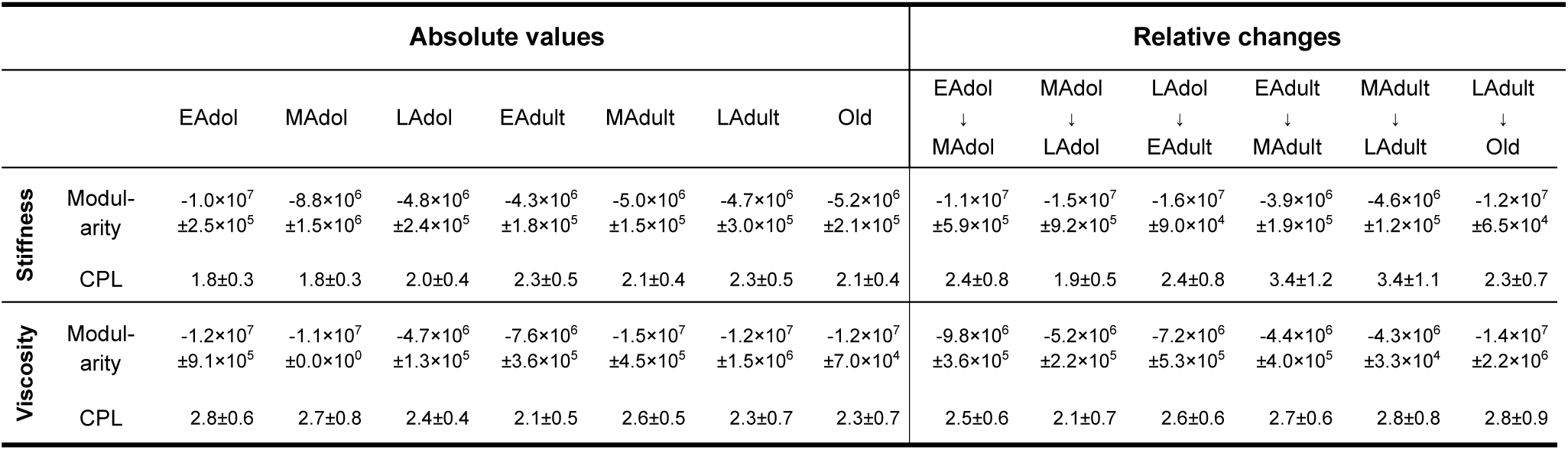
Group-averaged modularity and characteristic path length calculated from the absolute values of stiffness and viscosity, as well as from the relative changes in these parameters across the lifespan. Standard deviations were obtained from 100 bootstrap iterations for modularity and across nine iterative *τ* values for characteristic path length.

## Discussion

In this study, we investigated lifespan changes in mouse cerebral biomechanical properties, their connectivity, and network organization using in vivo MRE and community detection analysis. Our data revealed that, at the global level, the brain exhibits progressive softening and increased viscosity with aging. Previous MRE studies have demonstrated that brain mechanical properties change across the lifespan. In humans, aging is generally associated with reductions in brain stiffness and viscoelasticity, reflecting microstructural alterations such as neuronal loss, extracellular matrix remodeling, and altered tissue water content[20, 21]. Similar biomechanical alterations have been observed in murine models using MRE, indicating that brain biomechanics are sensitive to structural changes occurring during aging and neurodegeneration[22, 23]. As shown by our data, in the mouse, age-related softening was earliest and most pronounced in the hypothalamus. This finding aligns with evidence that the hypothalamus is among the first brain regions exhibiting molecular aging signatures, and may act as an early regulatory node of brain aging, characterized by neuronal dysfunction, reactive gliosis, chronic low-grade inflammation, as well as remodeling of the extracellular matrix and vascular microenvironment[24–26]. In contrast, CA and DG, subfields of the hippocampal formation, did not display significant age-associated softening in our study. This is consistent with reports that DG is one of the few brain areas where neurogenesis persists into adulthood[27] and the aged murine hippocampus largely preserves its population of principal neurons[28]. Our data also revealed that most brain regions exhibit increased viscosity from late adulthood onwards. However, the thalamus showed consistently lower viscosity from late adolescence through adulthood to old age, likely reflecting its stable nuclear architecture, dense myelinated tracts, and largely preserved extracellular matrix[29, 30]. This compact structural organization may support high regional integrity throughout maturation, contributing to its relatively low viscosity.

Biomechanical connectivity (BC) was introduced as a measure of age-specific regional similarity (static BC, sBC) or age-dependent covariation (dynamic BC, dBC) in cerebral biomechanical properties, including stiffness and viscosity. Because in vivo cerebral biomechanics reflect tissue structure, composition [31], and hemodynamic factors such as blood flow and perfusion [32, 33], we hypothesized that regions with similar biomechanical properties or coordinated trajectories across the lifespan may share structural and/or functional characteristics. Accordingly, we considered such regions “connected” within our biomechanical connectivity framework. To test this hypothesis, we compared the two BCs (sBC and dBC), with established measures of structural and functional connectivity. The results demonstrated a stronger resemblance between BC and FC than between BC and SC. This may partly reflect the limited brain coverage of BC, which excluded the olfactory areas and 74.7% of the isocortex, regions in which SC was particularly prominent. Moreover, SC derived from viral tracing primarily captures axonal projection density, representing only one component of the biological determinants of in vivo biomechanical properties. Other factors, including cellular density, extracellular matrix composition, and vascular architecture and perfusion, may also contribute to the biomechanical features measured in vivo. Thus, the observed correspondence between BC and FC, as well as the comparatively weaker correspondence with SC, may reflect the multidimensional biological determinants underlying regional cerebral biomechanics. Despite only partial resemblance to established structural and functional connectivity, the comparable overall similarity of sBC and dBC to FC and SC, together with stronger correspondence in selected regional pairs, supports biomechanical “connectivity” as a distinct measure of shared biomechanical characteristics, potentially reflecting underlying biological associations.

Based on the results of our community detection analysis, biomechanical network organization became increasingly structured during late adolescence (8 weeks), leading to a more specialized and segregated network architecture in adulthood. This pattern may reflect maturation processes, such as synaptic pruning and myelination[34, 35], which refine neural circuits during early and middle adolescence, when network organization patterns are more diffuse and variable. Decrease in the biomechanical network organization, and thus de-differentiation, was observed in old age, and is a characteristic feature of the aging mammalian brain[36]. This process reflects a progressive loss of structural organization and functional specialization. Similarly, early rodent studies reported that resting-state functional network connectivity increases from adolescence to adulthood and declines with aging, with ∼8 months marking a turning point in this trajectory [37, 38]. The peak of biomechanical segregation (∼8 weeks) observed in the current study precedes that of functional network connectivity, likely reflecting earlier structural network formation that scaffolds subsequent functional connections.

In addition to age-dependent global biomechanical segregation and integration, community assignments within brain subregions showed distinct age-associated fluctuations, particularly in viscosity-based measures. Most viscosity-defined communities remained stable across the lifespan, except during mid-adolescence to early adulthood. This period coincides with the last major organizational refinement in which the mouse brain transition to a stable, adult-like organization with consolidated circuitry and increased functional specialization[39, 40]. In contrast, stiffness-based community assignments did not reveal a comparable age-specific transition but exhibited substantial variability across the lifespan. This divergence suggests that viscosity may be more sensitive to global structural integrity, connectivity, and large-scale network organization, whereas stiffness reflects more local and subtle variations in cerebral microstructure, such as cellular composition and fiber density, which display greater variability over age.

Cerebral biomechanical network organization derived from the pace of age-related change (i.e. relative changes between adjacent age points) indicated a transition from globally coordinated dynamics in early adolescence to asynchronous, region-specific trajectories in midlife, followed by a more uniform decline in old age. This lifespan biomechanical shift in rate, coordination, and regional heterogeneity is consistent with heightened neuroplasticity during early adolescence, when widespread, concurrent remodeling occurs[41]. Thereafter, trajectories become increasingly specialized and heterogeneous, reflecting a transition from coordinated growth to maintenance-dominated processes[42]. In late adulthood, system-level constraints – including chronic inflammation, metabolic dysregulation, and loss of gene regulatory control, re-emerge and promote partial re-synchronization of regional dynamics[43, 44]. Furthermore, in contrast to community assignments derived from absolute stiffness values, those based on the pace of stiffness change remained remarkably stable across regions during adolescence. This stability suggests that, despite intrinsic regional differences in structure and composition, the mechanisms governing the buildup of tissue rigidity are largely similar.

Finally, we extended our network analysis to an experimental autoimmune encephalomyelitis (EAE) mouse model of multiple sclerosis (MS) to showcase the potential of biomechanical connectivity for detecting early MS-associated changes. The two datasets were acquired at the onset of disease progression. Based on our previous publication [15], none of the conventional imaging parameters, including MRE parameters, showed significant differences between EAE mice and healthy controls at this early stage. In contrast, in both cases, our proposed network analysis already revealed in both cases a reduction in the number of viscosity-based communities from 6 in the age- and sex-matched healthy mice (10 weeks) presented in the current study to 4 in the EAE mice (Supplemental Figure SF3). This very preliminary finding serves as a promising outlook for future applications without extending beyond the scope of the current study.

Despite its methodological strengths and broad age-span coverage, this study has limitations. First, the MR imaging volume was limited by coil coverage, precluding analysis of regions such as the olfactory bulb and cerebellum. Second, only female mice were examined; given known sex-dependence in brain maturation and aging[45], future studies will include age-matched male cohorts. Third, the cross-sectional design limits the ability to longitudinally assess the influence of social and environmental factors on cerebral aging. Finally, behavioral testing was not incorporated, precluding direct assessment of cognitive and motor performance in the context of aging.

In summary, we established cerebral biomechanical connectivity (BC) in mice using in vivo MRE and benchmarked it against established structural and functional connectivity, demonstrating its biological relevance. By integrating age-dependent BC with network-based analyses, we demonstrated that the healthy lifespan of the mouse brain is characterized by increased network biomechanical segregation during adolescence, followed by progressive re-integration in aging. Notably, the peak of biomechanical segregation precedes that reported for functional networks, suggesting a temporally distinct sequence in the emergence and organization of structural and functional networks. Network organization analyses based on the pace of age-related biomechanical change further reveal a coherent lifespan trajectory, spanning early global coordination, midlife functional and structural differentiation, and late-life re-synchronization. Together, these findings provide a comprehensive framework of biomechanical connectivity and its network organization across maturation and aging in the mouse brain, establishing a reference blueprint for future studies of neurological and neurodegenerative conditions that disrupt biomechanical network organization. Furthermore, age-related systems-level reorganization of biomechanical networks may mark key stages of brain maturation and periods of increased neurological vulnerability.

## Methods

### Animal use and experiment design

All experimental protocols were approved by the Berlin State Office for Health and Social Affairs (LaGeSo, 2019-106-G, G0284/13) and performed at Charité – Universitätsmedizin Berlin. Animal testing and research conformed to the guidelines set forth in the EU Directive 2010/63 and complied with the Federal German animal protection laws. A total of 63 female wild-type C57BL/6J mice (Charles Rivers Laboratories) were used in the study and maintained at the institutional animal facility with normal water and food supply *ad libitum*. To assess the biomechanical changes of the mouse brain from adolescence through adulthood to old age, seven phases in the mouse life span were investigated using in vivo MRI and MRE. Firstly, based on the timing of behavioral and neurological changes characteristic of adolescence[46], three adolescent periods were included: early adolescence (EAdol, n = 10 at 4 weeks old), mid-adolescence (MAdol, n = 10 at 6 weeks old), late adolescence (LAdol, n = 10 at 8 weeks old). These periods were followed by early adulthood (EAdult, n = 10 at 10-12 weeks old), mature adulthood (MAdult, n = 10 at 16-20 weeks old), late adulthood (LAdult, n = 8 at 48 weeks old), and old age (Old, n = 5 at 72 weeks old). The study design and the timeline is shown in Figure 7A.

**Figure 7.**
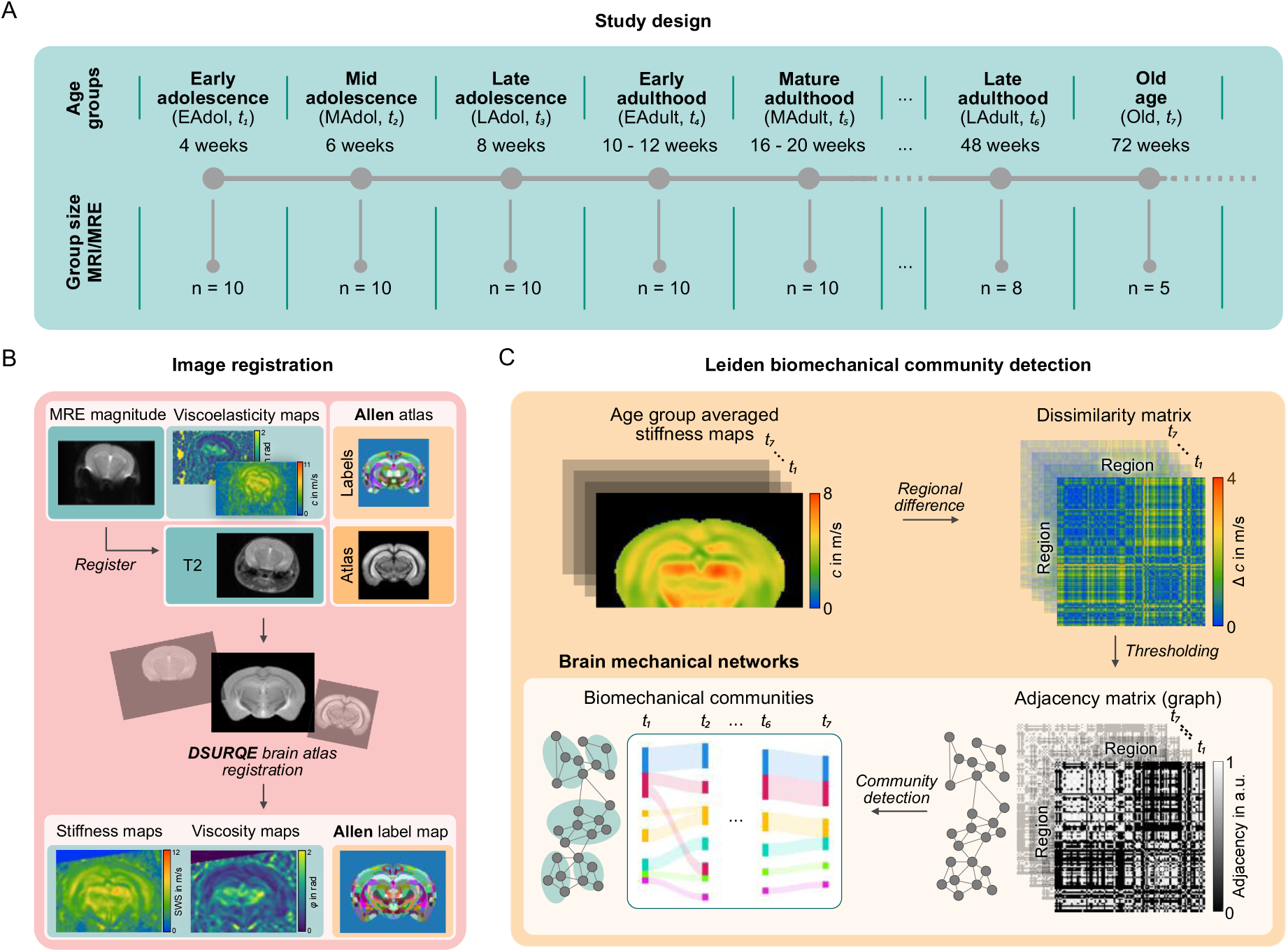
Overview of the study design and analytical workflow. (A) Definition of the age phases and corresponding sample sizes for the imaging examinations. (B) Schematic illustration of the image-registration pipeline. (C) Workflow of the Leiden community detection analysis, illustrated using the stiffness map as an example.

### MRI and MRE image acquisition and postprocessing

In vivo cerebral MRI and MRE data was acquired on a 7T small animal scanner (Bruker, Biospec, Ettlingen, Germany, Paravision 6.0.1) using a 20-mm volume coil (RAPID Biomedical, Rimpar, Germany). During the measurement, animals were anesthetized with 1.5% isoflurane with a mixture of 70% O_2_ and 30% N_2_O of gas while measuring *in vivo*. The application of a circulating water warming pad kept the mouse body temperature at 36.0 ± 1.0 °C.

### T2-weighted (T2w) imaging

T2w anatomical mages were obtained using a rapid acquisition with refocused echoes (RARE) sequence. 9 continuous axial slices with an in-plan resolution of 0.078 × 0.078 mm² and a slice thickness of 1 mm. Imaging parameters were: echo times (TE) of 54 ms, repetition time (TR) of 1800 ms, RARE factor of 8. The total scan duration was 2.8 minutes.

### Multifrequency MRE

For MRE measurements, harmonic vibrations of 1000, 1100, 1200, 1300 and 1400 Hz were generated by a non-magnetic piezo-ceramic actuator (CEDRAT Technologies, Meylan Cedex, France) and transferred to the brain via a head holder as depicted in [13]. 3D-wave fields were acquired in 9 contiguous slices of 0.2×0.2×1 mm^3^ resolution at eight time steps and 3 motion encoding gradients (MEG) using a single-shot spin-echo echo-planar imaging (EPI) sequence, the total acquisition time was 1.5 minutes. Further imaging parameters were: echo time: 52 ms; repetition time: 2000 ms.

MRE wave data were reconstructed using k-MDEV, a wave-number based multifrequency inversion method as described in [47, 48]. In brief, raw phase was unwrapped before Fourier transformation, followed by further removal of compression wave components using a third-order high-pass Butterworth filter (threshold of 200 m⁻¹). Finally, the wave field was decomposed into 4 orthogonal directions using a cos² directional filter as input for the gradient inversion. The inversion provided shear wave speed (*c*, m/s) and penetration rate (*a*, m/s). Phase angle *φ* was reconstructed from *c* and *a* with

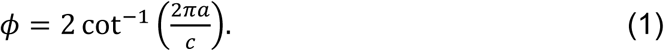

*c* and *φ* are surrogates for tissue stiffness and viscosity, respectively. To avoid areas with blood and corticospinal fluid, a shear wave speed threshold of 2.5 m/s was used for MRE parameter averaging[49].

### Image registration and elastographic atlases

A hierarchical image registration was performed using ANTx2 method described in [50]. Briefly, individual magnitude images obtained from the MRE scans were co-registered to the corresponding T2-weighted RARE images using elastix[51]. The co-registered MRE magnitude images were further transformed first to the MRI-based DSURQE atlas[52, 53] then to the Allen mouse brain atlas (Allen Institute for Brain Science, USA)[54]. The resulting transformation matrices were then applied directly to the MRE parameter maps of shear wave speed *c* and phase angle *φ* to transfer both maps into the standard Allen atlas space. Finally, elastographic atlases for *c* and *φ* at each age point were created by averaging the registered maps of all mice in the group. The steps involved in image registration and elastographic atlas generation are illustrated in Figure 7B. For subregional analysis, biomechanic parameters were averaged for 122 regions from the eight major areas prescribed in the Allen Brain Atlas: isocortex, hippocampal formation, cortical subplate, striatum, pallidum, hypothalamus, thalamus, and fiber tracts.

### Analysis of biomechanical, functional, and structural connectivity

BC, constructed based on age-specific biomechanical similarity and age-dependent regional covariance, was first compared with FC derived from resting-state fMRI (rs-fMRI) and SC derived from viral tracing data (Allen Institute, [55]). The parcellation scheme (i.e., region of interest (ROI) labels) was defined according to the spatial coverage of the MRE data and subsequently applied to both the rs-fMRI and SC datasets. Specifically, for BC analyses only, the Allen ROI labels were further divided by hemisphere to enable separate analyses of ipsilateral and contralateral connections, facilitating direct comparisons with FC and SC.

BC matrices were constructed either at specific age points (16-20 weeks for rs-fMRI and 8 weeks for SC), termed static BC (sBC), or across the lifespan by quantifying age-dependent interregional covariance, termed dynamic BC (dBC). Age-specific connectivity was represented by a metric of similarity as described in [56–58]. Let 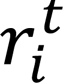 be a biomechanical parameters value corresponding to a region *i* for an age *t*. sBC matrices were then constructed by calculating the biomechanical dissimilarity 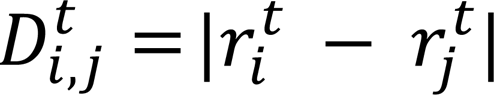 between all region pairs using either stiffness- or viscosity-based elastographic atlases. The resulting similarity matrices were exponentially transformed and rescaled to a range of 0-1. dBC matrices were generated by calculating Spearman correlation coefficients between the biomechanical parameters (stiffness and viscosity) of each pair of brain regions across all age points, followed by Fisher z transformation and rescaling to a range of 0-1.

The rs-fMRI data was obtained from a previous study (C57BL/6 mice, female, n = 10, imaged over 2 sessions, with 3 runs per sessions) and are publicly available at 10.18112/openneuro.ds001653.v1.0.2. Mice were anesthetized with isoflurane, positioned onto a MRI cradle, and anesthesia maintenance was provided with 1.2% isoflurane delivered via face mask. Standard gradient-echo echo planar images (TR/TE: 1200/15 ms, FA 60°, 180 volumes per run) were acquired at 11.7T with a 2×2 phased-array cryogenic probe. Data-processing pipeline and confound correction was carried out with RABIES 0.6.0 [53]. The confound correction model included a 0.01-0.2 Hz bandpass filter, aCompCor regression, and a 0.5 mm^2^ spatial filter [59]. FC between ROIs was calculated using Pearson correlation coefficients and subsequently Fisher z transformed. SC, quantified as normalized projection strength and obtained from Supplemental Table 3 in [55], was likewise log-transformed and rescaled to the range of 0-1.

To assess the overall agreement between BCs and FC, as well as between BCs and SC, receiver operating characteristic (ROC) analysis was performed following the approach described by [60]. Briefly, each of the connectivity matrices (BCs, FC, and SC) was thresholded across 1,000 evenly spaced thresholds corresponding to connection densities ranging from 0% to 100%. For each threshold, BC was benchmarked against either FC or SC, and the corresponding sensitivity and specificity were calculated. The resulting area under the curve (AUC) was used as a measure of the overall similarity between the biomechanical and functional or structural connectivity patterns.

### Leiden community detection

Leiden community detection[61] was performed on the aforementioned elastographic atlases. To assess the biomechanical organization of brain networks, graphs *G^t^* = (*N^t^*, *E^t^*) were constructed for all MRE parameters for all timepoints. In the graph *G^t^*, each vertex in *N^t^* represented a regional biomechanical parameter value 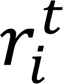. Adjacency matrices 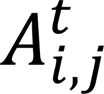 for each biomechanical dissimilarity *D_t_^i,j^* were constructed as follows:

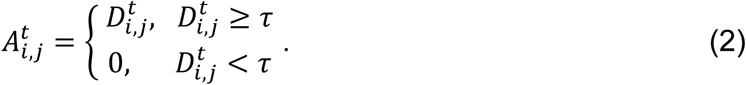

For each candidate value the resulting edge density was computed, and *τ* was set to the value yielding an approximately balanced graph, i.e. equal proportions of retained and discarded region pairs (edge density ≈ 50%). To confirm that this choice did not determine the subsequent results, the graph construction was repeated across the full range of candidate thresholds and the topology of the resulting networks was quantified before community detection (Figure 5). CPL varied only marginally over this range, indicating that graph structure is largely insensitive to the specific value of *τ* and that the parameter selection is robust. The metrices 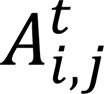 represented the weighted edges *E^t^* between vertices of graph *G^t^* according to Eq. (2).

Cerebral biomechanical communities were constructed with the Leiden algorithm. In brief, modularity based on the Constant Potts Model (CPM)[61, 62] quality metric was minimized to partition the nodes in *G^t^* into distinct communities *c*:

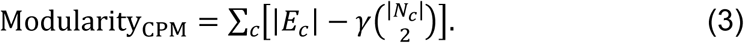

In Eq. (3), the resolution parameter *γ* acts as the inner and outer edge density threshold. It is tuned algorithmically via grid search to identify the smallest partition in which all communities exceed a minimum voxel count and lie in a community size range of 4 to 6. Small outlier communities falling below a filter threshold are removed beforehand. Measurements for differentiation include a varying bootstrapping-based partitioning result for 100 runs. The modularity parameter describes the strength of connections among nodes within the same community. Higher modularity is generally interpreted as greater network segregation, whereas reduced modularity mainly indicates increased integration. The community detection workflow is illustrated in Figure 7C.

To assess global efficiency of biomechanical network organization between brain regions independently of community structures, the CPL, which is the average of the shortest path lengths between all nodes (brain regions), was computed as follows:

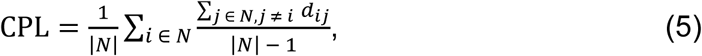

where *d_ij_* is the shortest path between node *i* and *j*. CPL and modularity represent complementary properties of brain network organization. Networks with higher modularity often exhibit increased CPL due to reduced within community connectivity, whereas lower modularity may facilitate shorter communication paths through enhanced network integration. As both modularity and CPL were derived from age-group-averaged biomechanical parameter maps to ensure numerical stability and robustness, they represent group-level parameters.

In addition to characterizing biomechanical communities at each age point, brain regions were also partitioned by the pace of their biomechanical changes over time, using community detection applied to difference maps between adjacent time points.

### Implementation and statistical analysis

Algorithmic calculation of Leiden communities, together with corresponding connectivity and adjacency matrices and statistical analysis was implemented and performed in Matlab (R2024a, MathWorks, USA). For statistical analysis, changes of biomechanical parameters over age were analyzed with the Kruskal-Wallis and the subsequent *post hoc* pairwise comparison with Bonferroni corrected Dunn’s test. Variance over group-level parameters (i.e., modularity) were estimated from 100 bootstrapped iterations. Statistics and the corresponding figures were performed and generated with R (4.4.1, R-Foundation, Austria).

## Acknowledgements

This work was supported by the German Research Foundation [grant numbers 460333672 CRC1540 EBM, RTG2260 BIOQIC, CRC1340 Matrix-in-vision, FOR5628, 540759292 Sa901/33-1 M5, Heisenberg Grant GU 1726/5-1 No. 504873209].

## Author contributions

Conceptualization: J.L., B.H., I.S. and J.Gu. Methodology: J.L., B.H., T.M., J.Gr., G.F., R.V.S., C.I., I.S. and J.Gu. Investigation: J.L., B.H., I.S. and J.Gu. Formal analysis: J.L., B.H., T.M. and J.Gu. Data curation: J.L., B.H., T.M., A.S.M. and J.Gu. Validation: J.L., B.H., T.M., C.I., I.S. and J.Gu. Visualization: J.L., B.H., T.M. and J.Gu. Writing–original draft: J.L., B.H., T.M. and J.Gu. Writing–review and editing: J.L., B.H., T.M., J.Gr., A.S.M., R.V.S., C.I., I.S. and J.Gu. Supervision: C.I., I.S. and J.Gu. Resources: B.H., J.Gr., R.V.S., C.I., I.S. and J.Gu. Project administration: M.E., I.S. and J.Gu. Funding acquisition: C.I., I.S. and J.Gu.

## Data availability

The data supporting the findings of this study is publicly available in the Zenodo repository at https://doi.org/10.5281/zenodo.20841001

## Supplementary material

**Supplemental Figure SF1.**
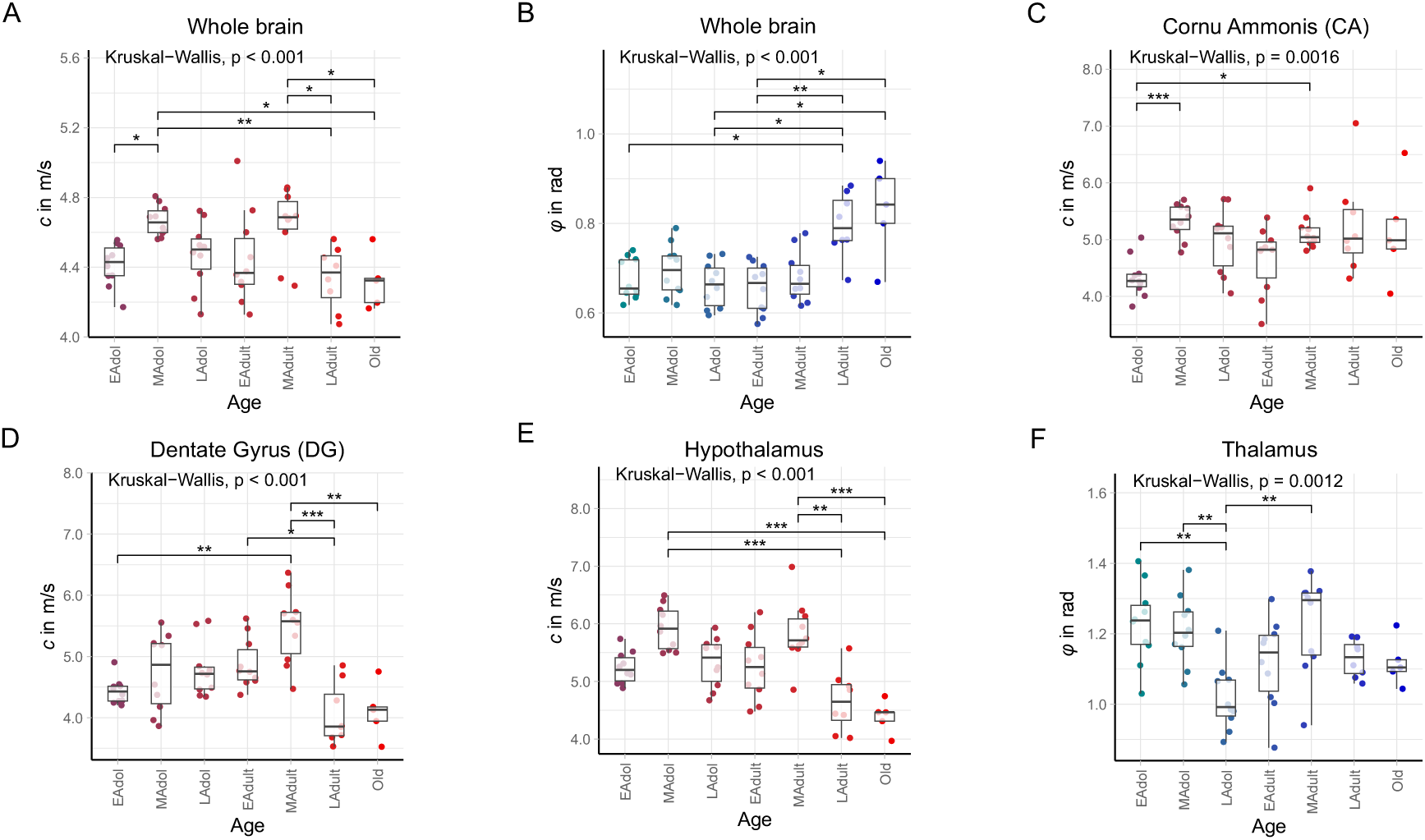
Box-and-scatter plots illustrating age-dependent changes in the biomechanical properties of selected brain regions. Statistically significant differences between age-groups, determined by *post hoc* pairwise comparisons, are indicated by asterisks. * P < 0.05, ** P < 0.01, *** P < 0.001.

**Supplemental Figure SF2.**
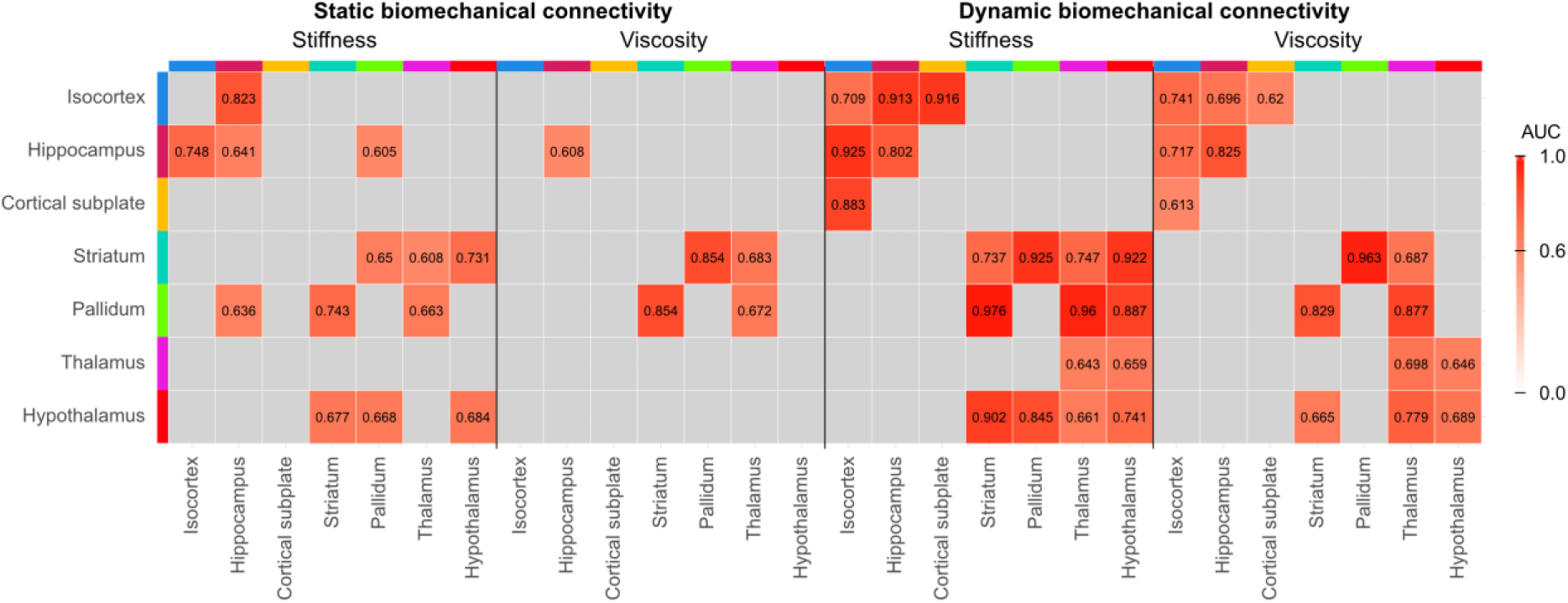
Interregional similarities between biomechanical connectivity (BC), including static BC (sBC) and dynamic BC (dBC), and functional connectivity (FC) or structural connectivity (SC), based on the matrices shown in Figure 3. Only region pairs with an AUC ≥ 0.6 are highlighted.

**Supplemental Figure SF3.**
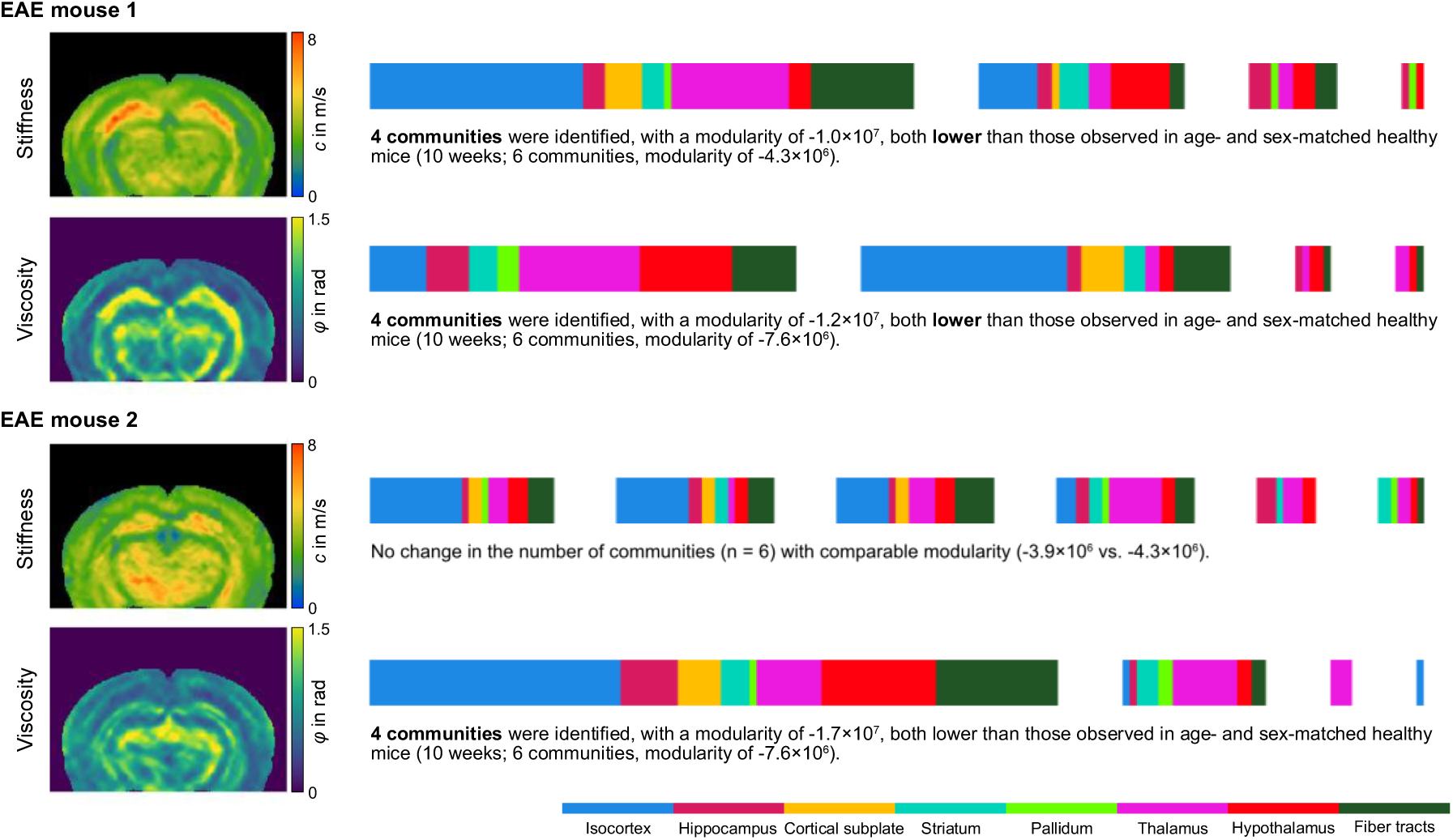
Community analysis of in vivo stiffness and viscosity in two experimental autoimmune encephalomyelitis (EAE) mice using the same parameters as in the present study. Parameter maps, detected communities, and color-coded brain regions with their corresponding community assignments are shown.

Supplemental Tables ST1 - ST4 are provided as supplementary Excel files.

**Supplemental Table ST1.**
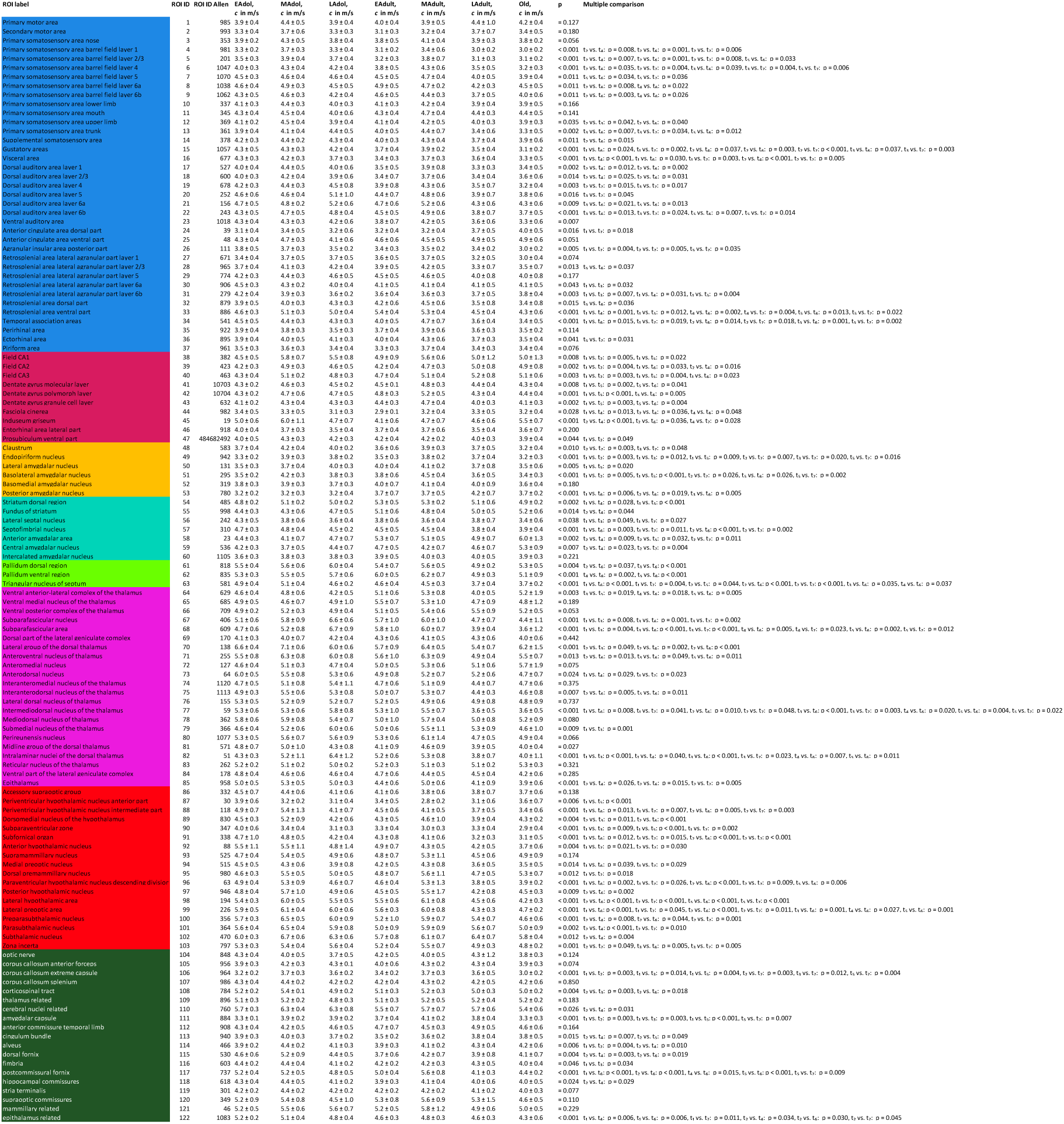
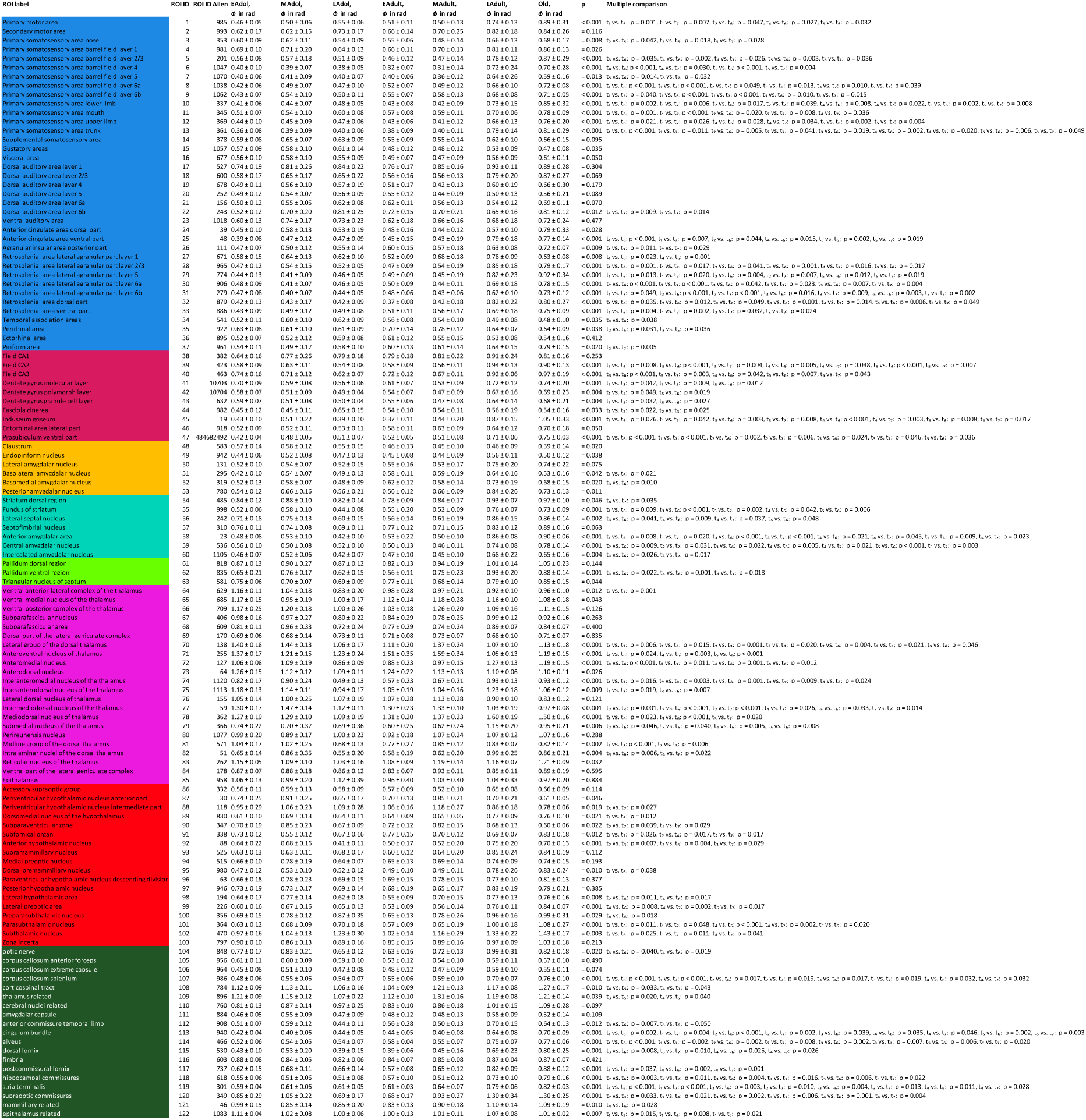
Group means ± standard deviations of the biomechanical properties for all 122 brain regions across age groups, along with the results of the corresponding Kruskal-Wallis analyses and post hoc pairwise comparisons.

| ROI label | ROI ID | ROI ID Allen | EAdol,<br>*c* in m/s | MAdol,<br>*c* in m/s | LAdol,<br>*c* in m/s | EAdult,<br>*c* in m/s | MAdult,<br>*c* in m/s | LAdult,<br>*c* in m/s | Old,<br>*c* in m/s | p | Multiple comparison |
|---|---|---|---|---|---|---|---|---|---|---|---|
| Primary motor area | 1 | 985 | 3.9 ± 0.4 | 4.4 ± 0.5 | 3.9 ± 0.4 | 4.0 ± 0.4 | 3.9 ± 0.5 | 4.4 ± 1.0 | 4.2 ± 0.4 | = 0.127 |  |
| Secondary motor area | 2 | 993 | 3.3 ± 0.3 | 3.7 ± 0.6 | 3.3 ± 0.3 | 3.1 ± 0.3 | 3.2 ± 0.4 | 3.7 ± 0.7 | 3.4 ± 0.5 | = 0.180 |  |
| Primary somatosensory area nose | 3 | 353 | 3.9 ± 0.2 | 4.3 ± 0.5 | 3.8 ± 0.4 | 3.8 ± 0.5 | 4.1 ± 0.4 | 3.9 ± 0.3 | 3.8 ± 0.2 | = 0.056 |  |
| Primary somatosensory area barrel field layer 1 | 4 | 981 | 3.3 ± 0.2 | 3.7 ± 0.3 | 3.3 ± 0.4 | 3.1 ± 0.2 | 3.4 ± 0.6 | 3.0 ± 0.2 | 3.0 ± 0.2 | < 0.001 | t<sub>2</sub> vs. t<sub>4</sub>: p = 0.008, t<sub>2</sub> vs. t<sub>6</sub>: p = 0.001, t<sub>2</sub> vs. t<sub>7</sub>: p = 0.006 |
| Primary somatosensory area barrel field layer 2/3 | 5 | 201 | 3.5 ± 0.3 | 3.9 ± 0.4 | 3.7 ± 0.4 | 3.2 ± 0.3 | 3.8 ± 0.7 | 3.1 ± 0.3 | 3.1 ± 0.2 | < 0.001 | t<sub>2</sub> vs. t<sub>4</sub>: p = 0.007, t<sub>2</sub> vs. t<sub>6</sub>: p = 0.001, t<sub>2</sub> vs. t<sub>7</sub>: p = 0.008, t<sub>3</sub> vs. t<sub>6</sub>: p = 0.033 |
| Primary somatosensory area barrel field layer 4 | 6 | 1047 | 4.0 ± 0.3 | 4.3 ± 0.4 | 4.2 ± 0.4 | 3.8 ± 0.5 | 4.3 ± 0.6 | 3.5 ± 0.5 | 3.2 ± 0.3 | < 0.001 | t<sub>1</sub> vs. t<sub>7</sub>: p = 0.035, t<sub>2</sub> vs. t<sub>7</sub>: p = 0.004, t<sub>3</sub> vs. t<sub>6</sub>: p = 0.039, t<sub>3</sub> vs. t<sub>7</sub>: p = 0.004, t<sub>5</sub> vs. t<sub>7</sub>: p = 0.006 |
| Primary somatosensory area barrel field layer 5 | 7 | 1070 | 4.5 ± 0.3 | 4.6 ± 0.4 | 4.6 ± 0.4 | 4.5 ± 0.5 | 4.7 ± 0.4 | 4.0 ± 0.5 | 3.9 ± 0.4 | = 0.011 | t<sub>5</sub> vs. t<sub>6</sub>: p = 0.034, t<sub>5</sub> vs. t<sub>7</sub>: p = 0.036 |
| Primary somatosensory area barrel field layer 6a | 8 | 1038 | 4.6 ± 0.4 | 4.9 ± 0.3 | 4.5 ± 0.5 | 4.9 ± 0.5 | 4.7 ± 0.2 | 4.2 ± 0.3 | 4.5 ± 0.5 | = 0.011 | t<sub>2</sub> vs. t<sub>6</sub>: p = 0.008, t<sub>4</sub> vs. t<sub>6</sub>: p = 0.022 |
| Primary somatosensory area barrel field layer 6b | 9 | 1062 | 4.3 ± 0.5 | 4.6 ± 0.3 | 4.2 ± 0.4 | 4.6 ± 0.5 | 4.4 ± 0.3 | 3.7 ± 0.5 | 4.0 ± 0.6 | = 0.011 | t<sub>2</sub> vs. t<sub>6</sub>: p = 0.003, t<sub>4</sub> vs. t<sub>6</sub>: p = 0.026 |
| Primary somatosensory area lower limb | 10 | 337 | 4.1 ± 0.3 | 4.4 ± 0.3 | 4.0 ± 0.3 | 4.1 ± 0.3 | 4.2 ± 0.4 | 3.9 ± 0.4 | 3.9 ± 0.5 | = 0.166 |  |
| Primary somatosensory area mouth | 11 | 345 | 4.3 ± 0.4 | 4.5 ± 0.4 | 4.0 ± 0.6 | 4.3 ± 0.4 | 4.7 ± 0.4 | 4.4 ± 0.3 | 4.4 ± 0.5 | = 0.141 |  |
| Primary somatosensory area upper limb | 12 | 369 | 4.1 ± 0.2 | 4.5 ± 0.4 | 3.9 ± 0.4 | 4.1 ± 0.4 | 4.2 ± 0.5 | 4.0 ± 0.3 | 3.9 ± 0.3 | = 0.035 | t<sub>2</sub> vs. t<sub>3</sub>: p = 0.042, t<sub>2</sub> vs. t<sub>7</sub>: p = 0.040 |
| Primary somatosensory area trunk | 13 | 361 | 3.9 ± 0.4 | 4.1 ± 0.4 | 4.4 ± 0.5 | 4.0 ± 0.5 | 4.4 ± 0.7 | 3.4 ± 0.6 | 3.3 ± 0.5 | = 0.002 | t<sub>3</sub> vs. t<sub>6</sub>: p = 0.007, t<sub>3</sub> vs. t<sub>7</sub>: p = 0.034, t<sub>5</sub> vs. t<sub>6</sub>: p = 0.012 |
| Supplemental somatosensory area | 14 | 378 | 4.2 ± 0.3 | 4.4 ± 0.2 | 4.3 ± 0.4 | 4.0 ± 0.3 | 4.3 ± 0.4 | 3.7 ± 0.4 | 3.9 ± 0.6 | = 0.011 | t<sub>2</sub> vs. t<sub>6</sub>: p = 0.015 |
| Gustatory areas | 15 | 1057 | 4.3 ± 0.5 | 4.3 ± 0.3 | 4.2 ± 0.4 | 3.7 ± 0.4 | 3.9 ± 0.2 | 3.5 ± 0.4 | 3.1 ± 0.2 | < 0.001 | t<sub>1</sub> vs. t<sub>6</sub>: p = 0.024, t<sub>1</sub> vs. t<sub>7</sub>: p = 0.002, t<sub>2</sub> vs. t<sub>6</sub>: p = 0.037, t<sub>2</sub> vs. t<sub>7</sub>: p = 0.003, t<sub>3</sub> vs. t<sub>7</sub>: p < 0.001, t<sub>4</sub> vs. t<sub>7</sub>: p = 0.037, t<sub>5</sub> vs. t<sub>7</sub>: p = 0.003 |
| Visceral area | 16 | 677 | 4.3 ± 0.3 | 4.2 ± 0.3 | 3.7 ± 0.3 | 3.4 ± 0.3 | 3.7 ± 0.3 | 3.6 ± 0.4 | 3.3 ± 0.5 | < 0.001 | t<sub>1</sub> vs. t<sub>4</sub>: p < 0.001, t<sub>1</sub> vs. t<sub>6</sub>: p = 0.030, t<sub>1</sub> vs. t<sub>7</sub>: p = 0.003, t<sub>2</sub> vs. t<sub>4</sub>: p < 0.001, t<sub>2</sub> vs. t<sub>7</sub>: p = 0.005 |
| Dorsal auditory area layer 1 | 17 | 527 | 4.0 ± 0.4 | 4.4 ± 0.5 | 4.0 ± 0.6 | 3.5 ± 0.5 | 3.9 ± 0.8 | 3.3 ± 0.3 | 3.4 ± 0.5 | = 0.002 | t<sub>2</sub> vs. t<sub>4</sub>: p = 0.012, t<sub>2</sub> vs. t<sub>6</sub>: p = 0.002 |
| Dorsal auditory area layer 2/3 | 18 | 600 | 4.0 ± 0.3 | 4.2 ± 0.4 | 3.9 ± 0.6 | 3.4 ± 0.7 | 3.7 ± 0.6 | 3.4 ± 0.4 | 3.6 ± 0.6 | = 0.014 | t<sub>2</sub> vs. t<sub>4</sub>: p = 0.025, t<sub>2</sub> vs. t<sub>6</sub>: p = 0.031 |
| Dorsal auditory area layer 4 | 19 | 678 | 4.2 ± 0.3 | 4.4 ± 0.3 | 4.5 ± 0.8 | 3.9 ± 0.8 | 4.3 ± 0.6 | 3.5 ± 0.7 | 3.2 ± 0.4 | = 0.003 | t<sub>2</sub> vs. t<sub>7</sub>: p = 0.015, t<sub>3</sub> vs. t<sub>7</sub>: p = 0.017 |
| Dorsal auditory area layer 5 | 20 | 252 | 4.6 ± 0.6 | 4.6 ± 0.4 | 5.1 ± 1.0 | 4.4 ± 0.7 | 4.8 ± 0.6 | 3.9 ± 0.7 | 3.8 ± 0.6 | = 0.016 | t<sub>3</sub> vs. t<sub>7</sub>: p = 0.045 |
| Dorsal auditory area layer 6a | 21 | 156 | 4.7 ± 0.5 | 4.8 ± 0.2 | 5.2 ± 0.6 | 4.7 ± 0.6 | 5.2 ± 0.6 | 4.3 ± 0.6 | 4.3 ± 0.6 | = 0.009 | t<sub>3</sub> vs. t<sub>6</sub>: p = 0.021, t<sub>5</sub> vs. t<sub>6</sub>: p = 0.013 |
| Dorsal auditory area layer 6b | 22 | 243 | 4.3 ± 0.5 | 4.7 ± 0.5 | 4.8 ± 0.4 | 4.5 ± 0.5 | 4.9 ± 0.6 | 3.8 ± 0.7 | 3.7 ± 0.5 | < 0.001 | t<sub>3</sub> vs. t<sub>6</sub>: p = 0.013, t<sub>3</sub> vs. t<sub>7</sub>: p = 0.024, t<sub>5</sub> vs. t<sub>6</sub>: p = 0.007, t<sub>5</sub> vs. t<sub>7</sub>: p = 0.014 |
| Ventral auditory area | 23 | 1018 | 4.3 ± 0.4 | 4.3 ± 0.3 | 4.2 ± 0.6 | 3.8 ± 0.7 | 4.2 ± 0.5 | 3.6 ± 0.6 | 3.3 ± 0.6 | = 0.007 |  |
| Anterior cingulate area dorsal part | 24 | 39 | 3.1 ± 0.3 | 3.4 ± 0.5 | 3.2 ± 0.6 | 3.2 ± 0.4 | 3.2 ± 0.4 | 3.7 ± 0.5 | 4.0 ± 0.5 | = 0.016 | t<sub>1</sub> vs. t<sub>7</sub>: p = 0.018 |
| Anterior cingulate area ventral part | 25 | 48 | 4.3 ± 0.4 | 4.7 ± 0.3 | 4.1 ± 0.6 | 4.6 ± 0.6 | 4.5 ± 0.5 | 4.9 ± 0.5 | 4.9 ± 0.6 | = 0.051 |  |
| Agranular insular area posterior part | 26 | 111 | 3.8 ± 0.5 | 3.7 ± 0.3 | 3.5 ± 0.2 | 3.4 ± 0.3 | 3.5 ± 0.2 | 3.4 ± 0.2 | 3.0 ± 0.2 | = 0.005 | t<sub>1</sub> vs. t<sub>7</sub>: p = 0.004, t<sub>2</sub> vs. t<sub>7</sub>: p = 0.005, t<sub>5</sub> vs. t<sub>7</sub>: p = 0.035 |
| Retrosplenial area lateral agranular part layer 1 | 27 | 671 | 3.4 ± 0.4 | 3.7 ± 0.5 | 3.7 ± 0.5 | 3.6 ± 0.5 | 3.7 ± 0.5 | 3.2 ± 0.5 | 3.0 ± 0.4 | = 0.074 |  |
| Retrosplenial area lateral agranular part layer 2/3 | 28 | 965 | 3.7 ± 0.4 | 4.1 ± 0.3 | 4.2 ± 0.4 | 3.9 ± 0.5 | 4.2 ± 0.5 | 3.3 ± 0.7 | 3.5 ± 0.7 | = 0.013 | t<sub>3</sub> vs. t<sub>6</sub>: p = 0.037 |
| Retrosplenial area lateral agranular part layer 5 | 29 | 774 | 4.2 ± 0.3 | 4.4 ± 0.3 | 4.6 ± 0.5 | 4.5 ± 0.5 | 4.6 ± 0.5 | 4.0 ± 0.8 | 4.0 ± 0.8 | = 0.177 |  |
| Retrosplenial area lateral agranular part layer 6a | 30 | 906 | 4.5 ± 0.3 | 4.3 ± 0.2 | 4.0 ± 0.4 | 4.1 ± 0.5 | 4.1 ± 0.4 | 4.1 ± 0.5 | 4.1 ± 0.5 | = 0.043 | t<sub>1</sub> vs. t<sub>3</sub>: p = 0.032 |
| Retrosplenial area lateral agranular part layer 6b | 31 | 279 | 4.2 ± 0.4 | 3.9 ± 0.3 | 3.6 ± 0.2 | 3.6 ± 0.4 | 3.6 ± 0.3 | 3.7 ± 0.5 | 3.8 ± 0.4 | = 0.003 | t<sub>1</sub> vs. t<sub>3</sub>: p = 0.007, t<sub>1</sub> vs. t<sub>4</sub>: p = 0.031, t<sub>1</sub> vs. t<sub>5</sub>: p = 0.004 |
| Retrosplenial area dorsal part | 32 | 879 | 3.9 ± 0.5 | 4.0 ± 0.3 | 4.3 ± 0.3 | 4.2 ± 0.6 | 4.5 ± 0.6 | 3.5 ± 0.8 | 3.4 ± 0.8 | = 0.015 | t<sub>5</sub> vs. t<sub>6</sub>: p = 0.036 |
| Retrosplenial area ventral part | 33 | 886 | 4.6 ± 0.3 | 5.1 ± 0.3 | 5.0 ± 0.4 | 5.4 ± 0.4 | 5.3 ± 0.4 | 4.5 ± 0.4 | 4.3 ± 0.6 | < 0.001 | t<sub>1</sub> vs. t<sub>4</sub>: p = 0.001, t<sub>1</sub> vs. t<sub>5</sub>: p = 0.012, t<sub>4</sub> vs. t<sub>6</sub>: p = 0.002, t<sub>4</sub> vs. t<sub>7</sub>: p = 0.004, t<sub>5</sub> vs. t<sub>6</sub>: p = 0.013, t<sub>5</sub> vs. t<sub>7</sub>: p = 0.022 |
| Temporal association areas | 34 | 541 | 4.5 ± 0.5 | 4.4 ± 0.3 | 4.3 ± 0.3 | 4.0 ± 0.5 | 4.7 ± 0.7 | 3.6 ± 0.4 | 3.4 ± 0.5 | < 0.001 | t<sub>1</sub> vs. t<sub>6</sub>: p = 0.015, t<sub>1</sub> vs. t<sub>7</sub>: p = 0.019, t<sub>2</sub> vs. t<sub>6</sub>: p = 0.014, t<sub>2</sub> vs. t<sub>7</sub>: p = 0.018, t<sub>5</sub> vs. t<sub>6</sub>: p = 0.001, t<sub>5</sub> vs. t<sub>7</sub>: p = 0.002 |
| Perirhinal area | 35 | 922 | 3.9 ± 0.4 | 3.8 ± 0.3 | 3.5 ± 0.3 | 3.7 ± 0.4 | 3.9 ± 0.6 | 3.6 ± 0.3 | 3.5 ± 0.2 | = 0.114 |  |
| Ectorhinal area | 36 | 895 | 3.9 ± 0.4 | 4.0 ± 0.5 | 4.1 ± 0.3 | 4.0 ± 0.4 | 4.3 ± 0.6 | 3.7 ± 0.3 | 3.5 ± 0.4 | = 0.041 | t<sub>5</sub> vs. t<sub>7</sub>: p = 0.031 |
| Piriform area | 37 | 961 | 3.5 ± 0.3 | 3.6 ± 0.3 | 3.4 ± 0.4 | 3.3 ± 0.3 | 3.7 ± 0.4 | 3.4 ± 0.3 | 3.4 ± 0.4 | = 0.076 |  |
| Field CA1 | 38 | 382 | 4.5 ± 0.5 | 5.8 ± 0.7 | 5.5 ± 0.8 | 4.9 ± 0.9 | 5.6 ± 0.6 | 5.0 ± 1.2 | 5.0 ± 1.3 | = 0.008 | t<sub>1</sub> vs. t<sub>2</sub>: p = 0.005, t<sub>1</sub> vs. t<sub>5</sub>: p = 0.022 |
| Field CA2 | 39 | 423 | 4.2 ± 0.3 | 4.9 ± 0.3 | 4.6 ± 0.5 | 4.2 ± 0.4 | 4.7 ± 0.3 | 5.0 ± 0.8 | 4.9 ± 0.8 | = 0.002 | t<sub>1</sub> vs. t<sub>2</sub>: p = 0.004, t<sub>1</sub> vs. t<sub>6</sub>: p = 0.033, t<sub>2</sub> vs. t<sub>4</sub>: p = 0.016 |
| Field CA3 | 40 | 463 | 4.3 ± 0.4 | 5.1 ± 0.2 | 4.8 ± 0.3 | 4.7 ± 0.4 | 5.1 ± 0.4 | 5.2 ± 0.8 | 5.1 ± 0.6 | = 0.003 | t<sub>1</sub> vs. t<sub>2</sub>: p = 0.003, t<sub>1</sub> vs. t<sub>5</sub>: p = 0.004, t<sub>1</sub> vs. t<sub>6</sub>: p = 0.023 |
| Dentate gyrus molecular layer | 41 | 10703 | 4.3 ± 0.2 | 4.6 ± 0.3 | 4.5 ± 0.2 | 4.5 ± 0.1 | 4.8 ± 0.3 | 4.4 ± 0.4 | 4.3 ± 0.4 | = 0.008 | t<sub>1</sub> vs. t<sub>5</sub>: p = 0.002, t<sub>5</sub> vs. t<sub>6</sub>: p = 0.041 |
| Dentate gyrus polymorph layer | 42 | 10704 | 4.3 ± 0.2 | 4.7 ± 0.6 | 4.7 ± 0.5 | 4.8 ± 0.3 | 5.2 ± 0.5 | 4.3 ± 0.4 | 4.4 ± 0.4 | = 0.001 | t<sub>1</sub> vs. t<sub>5</sub>: p < 0.001, t<sub>5</sub> vs. t<sub>6</sub>: p = 0.005 |
| Dentate gyrus granule cell layer | 43 | 632 | 4.1 ± 0.2 | 4.3 ± 0.4 | 4.3 ± 0.3 | 4.4 ± 0.3 | 4.7 ± 0.4 | 4.0 ± 0.3 | 4.1 ± 0.3 | = 0.002 | t<sub>1</sub> vs. t<sub>5</sub>: p = 0.003, t<sub>5</sub> vs. t<sub>6</sub>: p = 0.004 |
| Fasciola cinerea | 44 | 982 | 3.4 ± 0.5 | 3.3 ± 0.5 | 3.1 ± 0.3 | 2.9 ± 0.1 | 3.2 ± 0.4 | 3.3 ± 0.5 | 3.2 ± 0.4 | = 0.028 | t<sub>1</sub> vs. t<sub>4</sub>: p = 0.013, t<sub>2</sub> vs. t<sub>4</sub>: p = 0.036, t<sub>4</sub> vs. t<sub>6</sub>: p = 0.048 |
| Induseum griseum | 45 | 19 | 5.0 ± 0.6 | 6.0 ± 1.1 | 4.7 ± 0.7 | 4.1 ± 0.6 | 4.7 ± 0.7 | 4.6 ± 0.6 | 5.5 ± 0.7 | < 0.001 | t<sub>2</sub> vs. t<sub>4</sub>: p < 0.001, t<sub>2</sub> vs. t<sub>6</sub>: p = 0.036, t<sub>4</sub> vs. t<sub>7</sub>: p = 0.028 |
| Entorhinal area lateral part | 46 | 918 | 4.0 ± 0.4 | 3.7 ± 0.3 | 3.5 ± 0.4 | 3.7 ± 0.4 | 3.7 ± 0.6 | 3.5 ± 0.4 | 3.6 ± 0.7 | = 0.200 |  |
| Prosubiculum ventral part | 47 | 484682492 | 4.0 ± 0.5 | 4.3 ± 0.3 | 4.2 ± 0.3 | 4.2 ± 0.4 | 4.2 ± 0.2 | 4.0 ± 0.3 | 3.9 ± 0.4 | = 0.044 | t<sub>1</sub> vs. t<sub>3</sub>: p = 0.049 |
| Claustrum | 48 | 583 | 3.7 ± 0.4 | 4.2 ± 0.4 | 4.0 ± 0.2 | 3.6 ± 0.6 | 3.9 ± 0.3 | 3.7 ± 0.5 | 3.2 ± 0.4 | = 0.010 | t<sub>2</sub> vs. t<sub>7</sub>: p = 0.003, t<sub>3</sub> vs. t<sub>7</sub>: p = 0.048 |
| Endopiriform nucleus | 49 | 942 | 3.3 ± 0.2 | 3.9 ± 0.3 | 3.8 ± 0.2 | 3.5 ± 0.3 | 3.8 ± 0.2 | 3.7 ± 0.4 | 3.2 ± 0.3 | < 0.001 | t<sub>1</sub> vs. t<sub>2</sub>: p = 0.003, t<sub>1</sub> vs. t<sub>3</sub>: p = 0.012, t<sub>1</sub> vs. t<sub>5</sub>: p = 0.009, t<sub>2</sub> vs. t<sub>7</sub>: p = 0.007, t<sub>3</sub> vs. t<sub>7</sub>: p = 0.020, t<sub>5</sub> vs. t<sub>7</sub>: p = 0.016 |
| Lateral amygdalar nucleus | 50 | 131 | 3.5 ± 0.3 | 3.7 ± 0.4 | 4.0 ± 0.3 | 4.0 ± 0.4 | 4.1 ± 0.2 | 3.7 ± 0.8 | 3.5 ± 0.6 | = 0.005 | t<sub>1</sub> vs. t<sub>5</sub>: p = 0.020 |
| Basolateral amygdalar nucleus | 51 | 295 | 3.5 ± 0.2 | 4.2 ± 0.3 | 3.8 ± 0.3 | 3.8 ± 0.6 | 4.5 ± 0.4 | 3.6 ± 0.5 | 3.4 ± 0.3 | < 0.001 | t<sub>1</sub> vs. t<sub>2</sub>: p = 0.005, t<sub>1</sub> vs. t<sub>5</sub>: p < 0.001, t<sub>2</sub> vs. t<sub>7</sub>: p = 0.026, t<sub>5</sub> vs. t<sub>6</sub>: p = 0.026, t<sub>5</sub> vs. t<sub>7</sub>: p = 0.002 |
| Basomedial amygdalar nucleus | 52 | 319 | 3.8 ± 0.3 | 3.9 ± 0.3 | 3.7 ± 0.3 | 4.0 ± 0.7 | 4.1 ± 0.4 | 4.0 ± 0.9 | 3.6 ± 0.4 | = 0.180 |  |
| Posterior amygdalar nucleus | 53 | 780 | 3.2 ± 0.2 | 3.2 ± 0.3 | 3.2 ± 0.4 | 3.7 ± 0.7 | 3.7 ± 0.5 | 4.2 ± 0.7 | 3.7 ± 0.2 | < 0.001 | t<sub>1</sub> vs. t<sub>6</sub>: p = 0.006, t<sub>2</sub> vs. t<sub>6</sub>: p = 0.019, t<sub>3</sub> vs. t<sub>6</sub>: p = 0.005 |
| Striatum dorsal region | 54 | 485 | 4.8 ± 0.2 | 5.1 ± 0.2 | 5.0 ± 0.2 | 5.3 ± 0.5 | 5.3 ± 0.2 | 5.1 ± 0.6 | 4.9 ± 0.2 | = 0.002 | t<sub>1</sub> vs. t<sub>4</sub>: p = 0.028, t<sub>1</sub> vs. t<sub>5</sub>: p < 0.001 |
| Fundus of striatum | 55 | 998 | 4.4 ± 0.3 | 4.3 ± 0.6 | 4.7 ± 0.5 | 5.1 ± 0.6 | 4.8 ± 0.4 | 5.0 ± 0.5 | 5.2 ± 0.6 | = 0.014 | t<sub>2</sub> vs. t<sub>4</sub>: p = 0.044 |
| Lateral septal nucleus | 56 | 242 | 4.3 ± 0.5 | 3.8 ± 0.6 | 3.6 ± 0.4 | 3.8 ± 0.6 | 3.6 ± 0.4 | 3.8 ± 0.7 | 3.4 ± 0.6 | = 0.038 | t<sub>1</sub> vs. t<sub>3</sub>: p = 0.049, t<sub>1</sub> vs. t<sub>7</sub>: p = 0.027 |
| Septofimbrial nucleus | 57 | 310 | 4.7 ± 0.3 | 4.8 ± 0.4 | 4.5 ± 0.2 | 4.5 ± 0.5 | 4.5 ± 0.4 | 3.8 ± 0.4 | 3.9 ± 0.4 | < 0.001 | t<sub>1</sub> vs. t<sub>6</sub>: p = 0.003, t<sub>1</sub> vs. t<sub>7</sub>: p = 0.011, t<sub>2</sub> vs. t<sub>6</sub>: p < 0.001, t<sub>2</sub> vs. t<sub>7</sub>: p = 0.002 |
| Anterior amygdalar area | 58 | 23 | 4.4 ± 0.3 | 4.1 ± 0.7 | 4.7 ± 0.7 | 5.3 ± 0.7 | 5.1 ± 0.5 | 4.9 ± 0.7 | 6.0 ± 1.3 | = 0.002 | t<sub>2</sub> vs. t<sub>4</sub>: p = 0.009, t<sub>2</sub> vs. t<sub>5</sub>: p = 0.032, t<sub>2</sub> vs. t<sub>7</sub>: p = 0.011 |
| Central amygdalar nucleus | 59 | 536 | 4.2 ± 0.3 | 3.7 ± 0.5 | 4.4 ± 0.7 | 4.7 ± 0.5 | 4.2 ± 0.7 | 4.6 ± 0.7 | 5.3 ± 0.9 | = 0.007 | t<sub>2</sub> vs. t<sub>4</sub>: p = 0.023, t<sub>2</sub> vs. t<sub>7</sub>: p = 0.004 |
| Intercalated amygdalar nucleus | 60 | 1105 | 3.6 ± 0.3 | 3.8 ± 0.3 | 3.8 ± 0.3 | 3.9 ± 0.5 | 4.0 ± 0.3 | 4.0 ± 0.5 | 3.9 ± 0.3 | = 0.221 |  |
| Pallidum dorsal region | 61 | 818 | 5.5 ± 0.4 | 5.6 ± 0.6 | 6.0 ± 0.4 | 5.4 ± 0.7 | 5.6 ± 0.5 | 4.9 ± 0.2 | 5.3 ± 0.5 | = 0.004 | t<sub>2</sub> vs. t<sub>6</sub>: p = 0.037, t<sub>3</sub> vs. t<sub>6</sub>: p < 0.001 |
| Pallidum ventral region | 62 | 835 | 5.3 ± 0.3 | 5.5 ± 0.5 | 5.7 ± 0.6 | 6.0 ± 0.5 | 6.2 ± 0.7 | 4.9 ± 0.3 | 5.1 ± 0.9 | < 0.001 | t<sub>4</sub> vs. t<sub>6</sub>: p = 0.002, t<sub>5</sub> vs. t<sub>6</sub>: p < 0.001 |
| Triangular nucleus of septum | 63 | 581 | 4.9 ± 0.4 | 5.1 ± 0.4 | 4.6 ± 0.2 | 4.6 ± 0.4 | 4.5 ± 0.3 | 3.7 ± 0.4 | 3.7 ± 0.2 | < 0.001 | t<sub>1</sub> vs. t<sub>6</sub>: p < 0.001, t<sub>1</sub> vs. t<sub>7</sub>: p = 0.004, t<sub>2</sub> vs. t<sub>5</sub>: p = 0.044, t<sub>2</sub> vs. t<sub>6</sub>: p < 0.001, t<sub>2</sub> vs. t<sub>7</sub>: p < 0.001, t<sub>3</sub> vs. t<sub>6</sub>: p = 0.035, t<sub>4</sub> vs. t<sub>6</sub>: p = 0.037 |
| Ventral anterior-lateral complex of the thalamus | 64 | 629 | 4.6 ± 0.4 | 4.8 ± 0.6 | 4.2 ± 0.5 | 5.1 ± 0.6 | 5.3 ± 0.8 | 4.0 ± 0.5 | 5.2 ± 1.9 | = 0.003 | t<sub>3</sub> vs. t<sub>5</sub>: p = 0.019, t<sub>4</sub> vs. t<sub>6</sub>: p = 0.018, t<sub>5</sub> vs. t<sub>6</sub>: p = 0.005 |
| Ventral medial nucleus of the thalamus | 65 | 685 | 4.9 ± 0.5 | 4.6 ± 0.7 | 4.9 ± 1.0 | 5.5 ± 0.7 | 5.3 ± 1.0 | 4.7 ± 0.9 | 4.8 ± 1.2 | = 0.189 |  |
| Ventral posterior complex of the thalamus | 66 | 709 | 4.9 ± 0.2 | 5.2 ± 0.3 | 4.9 ± 0.4 | 5.1 ± 0.5 | 5.4 ± 0.6 | 5.5 ± 0.9 | 5.2 ± 0.5 | = 0.053 |  |
| Subparafascicular nucleus | 67 | 406 | 5.1 ± 0.6 | 5.8 ± 0.9 | 6.6 ± 0.6 | 5.7 ± 1.0 | 6.0 ± 1.0 | 4.7 ± 0.7 | 4.4 ± 1.1 | < 0.001 | t<sub>1</sub> vs. t<sub>3</sub>: p = 0.008, t<sub>3</sub> vs. t<sub>6</sub>: p = 0.001, t<sub>3</sub> vs. t<sub>7</sub>: p = 0.002 |
| Subparafascicular area | 68 | 609 | 4.7 ± 0.6 | 5.2 ± 0.8 | 6.7 ± 0.9 | 5.8 ± 1.0 | 6.0 ± 0.7 | 3.9 ± 0.4 | 3.6 ± 1.2 | < 0.001 | t<sub>1</sub> vs. t<sub>3</sub>: p = 0.004, t<sub>3</sub> vs. t<sub>6</sub>: p < 0.001, t<sub>3</sub> vs. t<sub>7</sub>: p < 0.001, t<sub>4</sub> vs. t<sub>6</sub>: p = 0.005, t<sub>4</sub> vs. t<sub>7</sub>: p = 0.023, t<sub>5</sub> vs. t<sub>6</sub>: p = 0.002, t<sub>5</sub> vs. t<sub>7</sub>: p = 0.012 |
| Dorsal part of the lateral geniculate complex | 69 | 170 | 4.1 ± 0.3 | 4.0 ± 0.7 | 4.2 ± 0.4 | 4.3 ± 0.6 | 4.1 ± 0.5 | 4.3 ± 0.6 | 4.0 ± 0.6 | = 0.442 |  |
| Lateral group of the dorsal thalamus | 70 | 138 | 6.6 ± 0.4 | 7.1 ± 0.6 | 6.0 ± 0.6 | 5.7 ± 0.9 | 6.4 ± 0.5 | 5.4 ± 0.7 | 6.2 ± 1.5 | < 0.001 | t<sub>2</sub> vs. t<sub>3</sub>: p = 0.049, t<sub>2</sub> vs. t<sub>4</sub>: p = 0.002, t<sub>2</sub> vs. t<sub>6</sub>: p < 0.001 |
| Anteroventral nucleus of thalamus | 71 | 255 | 5.5 ± 0.8 | 6.3 ± 0.8 | 6.0 ± 0.8 | 5.6 ± 1.0 | 6.3 ± 0.9 | 4.9 ± 0.4 | 5.5 ± 0.7 | = 0.013 | t<sub>2</sub> vs. t<sub>6</sub>: p = 0.013, t<sub>3</sub> vs. t<sub>6</sub>: p = 0.049, t<sub>5</sub> vs. t<sub>6</sub>: p = 0.011 |
| Anteromedial nucleus | 72 | 127 | 4.6 ± 0.4 | 5.1 ± 0.3 | 4.7 ± 0.4 | 5.0 ± 0.5 | 5.3 ± 0.6 | 5.1 ± 0.6 | 5.7 ± 1.9 | = 0.075 |  |
| Anterodorsal nucleus | 73 | 64 | 6.0 ± 0.5 | 5.5 ± 0.8 | 5.3 ± 0.6 | 4.9 ± 0.8 | 5.2 ± 0.7 | 5.2 ± 0.6 | 4.7 ± 0.7 | = 0.024 | t<sub>1</sub> vs. t<sub>4</sub>: p = 0.029, t<sub>1</sub> vs. t<sub>7</sub>: p = 0.023 |
| Interanteromedial nucleus of the thalamus | 74 | 1120 | 4.7 ± 0.5 | 5.1 ± 0.8 | 5.4 ± 1.1 | 4.7 ± 0.6 | 5.1 ± 0.9 | 4.4 ± 0.7 | 4.7 ± 0.6 | = 0.375 |  |
| Interanterodorsal nucleus of the thalamus | 75 | 1113 | 4.9 ± 0.3 | 5.5 ± 0.6 | 5.3 ± 0.8 | 5.0 ± 0.7 | 5.3 ± 0.7 | 4.4 ± 0.3 | 4.6 ± 0.8 | = 0.007 | t<sub>2</sub> vs. t<sub>6</sub>: p = 0.005, t<sub>5</sub> vs. t<sub>6</sub>: p = 0.011 |
| Lateral dorsal nucleus of thalamus | 76 | 155 | 5.3 ± 0.5 | 5.2 ± 0.9 | 5.2 ± 0.7 | 5.2 ± 0.5 | 4.9 ± 0.6 | 4.9 ± 0.8 | 4.8 ± 0.9 | = 0.737 |  |
| Intermediodorsal nucleus of the thalamus | 77 | 59 | 5.3 ± 0.6 | 5.3 ± 0.6 | 5.8 ± 0.8 | 5.3 ± 1.0 | 5.5 ± 0.7 | 3.6 ± 0.5 | 3.6 ± 0.5 | < 0.001 | t<sub>1</sub> vs. t<sub>6</sub>: p = 0.008, t<sub>1</sub> vs. t<sub>7</sub>: p = 0.041, t<sub>2</sub> vs. t<sub>6</sub>: p = 0.010, t<sub>2</sub> vs. t<sub>7</sub>: p = 0.048, t<sub>3</sub> vs. t<sub>6</sub>: p < 0.001, t<sub>3</sub> vs. t<sub>7</sub>: p = 0.003, t<sub>4</sub> vs. t<sub>6</sub>: p = 0.020, t<sub>5</sub> vs. t<sub>6</sub>: p = 0.004, t<sub>5</sub> vs. t<sub>7</sub>: p = 0.022 |
| Mediodorsal nucleus of thalamus | 78 | 362 | 5.8 ± 0.6 | 5.9 ± 0.8 | 5.4 ± 0.7 | 5.0 ± 1.0 | 5.7 ± 0.4 | 5.0 ± 0.8 | 5.2 ± 0.9 | = 0.080 |  |
| Submedial nucleus of the thalamus | 79 | 366 | 4.6 ± 0.4 | 5.2 ± 0.6 | 6.0 ± 0.6 | 5.0 ± 0.6 | 5.5 ± 1.1 | 5.3 ± 0.9 | 4.6 ± 1.0 | = 0.009 | t<sub>1</sub> vs. t<sub>3</sub>: p = 0.001 |
| Perireunensis nucleus | 80 | 1077 | 5.3 ± 0.5 | 5.6 ± 0.7 | 5.6 ± 0.9 | 5.3 ± 0.6 | 6.1 ± 1.4 | 4.7 ± 0.5 | 4.9 ± 0.4 | = 0.066 |  |
| Midline group of the dorsal thalamus | 81 | 571 | 4.8 ± 0.7 | 5.0 ± 1.0 | 4.3 ± 0.8 | 4.1 ± 0.9 | 4.6 ± 0.9 | 3.9 ± 0.5 | 4.0 ± 0.4 | = 0.027 |  |
| Intralaminar nuclei of the dorsal thalamus | 82 | 51 | 4.3 ± 0.3 | 5.2 ± 1.1 | 6.4 ± 1.2 | 5.2 ± 0.6 | 5.3 ± 0.8 | 3.8 ± 0.7 | 4.0 ± 1.1 | < 0.001 | t<sub>1</sub> vs. t<sub>3</sub>: p < 0.001, t<sub>2</sub> vs. t<sub>6</sub>: p = 0.040, t<sub>3</sub> vs. t<sub>6</sub>: p < 0.001, t<sub>3</sub> vs. t<sub>7</sub>: p = 0.023, t<sub>4</sub> vs. t<sub>6</sub>: p = 0.007, t<sub>5</sub> vs. t<sub>6</sub>: p = 0.011 |
| Reticular nucleus of the thalamus | 83 | 262 | 5.2 ± 0.2 | 5.1 ± 0.2 | 5.0 ± 0.2 | 5.2 ± 0.3 | 5.1 ± 0.3 | 5.1 ± 0.2 | 5.3 ± 0.3 | = 0.321 |  |
| Ventral part of the lateral geniculate complex | 84 | 178 | 4.8 ± 0.4 | 4.6 ± 0.6 | 4.6 ± 0.4 | 4.7 ± 0.6 | 4.4 ± 0.5 | 4.5 ± 0.4 | 4.2 ± 0.6 | = 0.285 |  |
| Epithalamus | 85 | 958 | 5.0 ± 0.5 | 5.3 ± 0.5 | 5.0 ± 0.3 | 4.4 ± 0.6 | 5.0 ± 0.6 | 4.1 ± 0.9 | 3.9 ± 0.6 | < 0.001 | t<sub>2</sub> vs. t<sub>4</sub>: p = 0.026, t<sub>2</sub> vs. t<sub>6</sub>: p = 0.015, t<sub>2</sub> vs. t<sub>7</sub>: p = 0.005 |
| Accessory supraoptic group | 86 | 332 | 4.5 ± 0.7 | 4.4 ± 0.6 | 4.1 ± 0.6 | 4.1 ± 0.6 | 3.8 ± 0.6 | 3.8 ± 0.7 | 3.7 ± 0.6 | = 0.138 |  |
| Periventricular hypothalamic nucleus anterior part | 87 | 30 | 3.9 ± 0.6 | 3.2 ± 0.2 | 3.1 ± 0.4 | 3.4 ± 0.5 | 2.8 ± 0.2 | 3.1 ± 0.6 | 3.6 ± 0.7 | = 0.006 | t<sub>1</sub> vs. t<sub>5</sub>: p < 0.001 |
| Periventricular hypothalamic nucleus intermediate part | 88 | 118 | 4.9 ± 0.7 | 5.4 ± 1.3 | 4.1 ± 0.7 | 4.5 ± 0.6 | 4.1 ± 0.5 | 3.7 ± 0.5 | 3.4 ± 0.6 | < 0.001 | t<sub>1</sub> vs. t<sub>6</sub>: p = 0.013, t<sub>1</sub> vs. t<sub>7</sub>: p = 0.007, t<sub>2</sub> vs. t<sub>6</sub>: p = 0.005, t<sub>2</sub> vs. t<sub>7</sub>: p = 0.003 |
| Dorsomedial nucleus of the hypothalamus | 89 | 830 | 4.5 ± 0.3 | 5.2 ± 0.9 | 4.2 ± 0.6 | 4.3 ± 0.5 | 4.6 ± 1.0 | 3.9 ± 0.4 | 4.3 ± 0.2 | = 0.004 | t<sub>2</sub> vs. t<sub>3</sub>: p = 0.011, t<sub>2</sub> vs. t<sub>6</sub>: p < 0.001 |
| Subparaventricular zone | 90 | 347 | 4.0 ± 0.6 | 3.4 ± 0.4 | 3.1 ± 0.3 | 3.3 ± 0.4 | 3.0 ± 0.3 | 3.3 ± 0.4 | 2.9 ± 0.4 | < 0.001 | t<sub>1</sub> vs. t<sub>3</sub>: p = 0.009, t<sub>1</sub> vs. t<sub>5</sub>: p < 0.001, t<sub>1</sub> vs. t<sub>7</sub>: p = 0.002 |
| Subfornical organ | 91 | 338 | 4.7 ± 1.0 | 4.8 ± 0.5 | 4.2 ± 0.4 | 4.3 ± 0.8 | 4.1 ± 0.6 | 3.2 ± 0.3 | 3.1 ± 0.5 | < 0.001 | t<sub>1</sub> vs. t<sub>6</sub>: p = 0.012, t<sub>1</sub> vs. t<sub>7</sub>: p = 0.015, t<sub>2</sub> vs. t<sub>6</sub>: p < 0.001, t<sub>2</sub> vs. t<sub>7</sub>: p < 0.001 |
| Anterior hypothalamic nucleus | 92 | 88 | 5.5 ± 1.1 | 5.5 ± 1.1 | 4.8 ± 1.4 | 4.9 ± 0.7 | 4.3 ± 0.5 | 4.2 ± 0.5 | 3.7 ± 0.6 | = 0.004 | t<sub>1</sub> vs. t<sub>7</sub>: p = 0.021, t<sub>2</sub> vs. t<sub>7</sub>: p = 0.030 |
| Supramammillary nucleus | 93 | 525 | 4.7 ± 0.4 | 5.4 ± 0.5 | 4.9 ± 0.6 | 4.8 ± 0.7 | 5.3 ± 1.1 | 4.5 ± 0.6 | 4.9 ± 0.9 | = 0.174 |  |
| Medial preoptic nucleus | 94 | 515 | 4.5 ± 0.5 | 4.3 ± 0.6 | 3.9 ± 0.8 | 4.2 ± 0.5 | 4.3 ± 0.8 | 3.6 ± 0.5 | 3.5 ± 0.5 | = 0.014 | t<sub>1</sub> vs. t<sub>6</sub>: p = 0.039, t<sub>1</sub> vs. t<sub>7</sub>: p = 0.029 |
| Dorsal premammillary nucleus | 95 | 980 | 4.6 ± 0.3 | 5.5 ± 0.5 | 5.0 ± 0.5 | 4.8 ± 0.7 | 5.6 ± 1.1 | 4.7 ± 0.5 | 5.3 ± 0.7 | = 0.012 | t<sub>1</sub> vs. t<sub>5</sub>: p = 0.018 |
| Paraventricular hypothalamic nucleus descending division | 96 | 63 | 4.9 ± 0.4 | 5.3 ± 0.9 | 4.6 ± 0.7 | 4.6 ± 0.4 | 5.3 ± 1.3 | 3.8 ± 0.5 | 3.9 ± 0.2 | < 0.001 | t<sub>1</sub> vs. t<sub>6</sub>: p = 0.002, t<sub>1</sub> vs. t<sub>7</sub>: p = 0.026, t<sub>2</sub> vs. t<sub>6</sub>: p < 0.001, t<sub>2</sub> vs. t<sub>7</sub>: p = 0.009, t<sub>5</sub> vs. t<sub>6</sub>: p = 0.006 |
| Posterior hypothalamic nucleus | 97 | 946 | 4.8 ± 0.4 | 5.7 ± 1.0 | 4.9 ± 0.6 | 4.5 ± 0.5 | 5.5 ± 1.7 | 4.2 ± 0.8 | 4.5 ± 0.3 | = 0.009 | t<sub>2</sub> vs. t<sub>6</sub>: p = 0.002 |
| Lateral hypothalamic area | 98 | 194 | 5.4 ± 0.3 | 6.0 ± 0.5 | 5.5 ± 0.5 | 5.5 ± 0.6 | 6.1 ± 0.8 | 4.5 ± 0.6 | 4.2 ± 0.3 | < 0.001 | t<sub>2</sub> vs. t<sub>6</sub>: p < 0.001, t<sub>2</sub> vs. t<sub>7</sub>: p < 0.001, t<sub>5</sub> vs. t<sub>6</sub>: p < 0.001, t<sub>5</sub> vs. t<sub>7</sub>: p < 0.001 |
| Lateral preoptic area | 99 | 226 | 5.9 ± 0.5 | 6.1 ± 0.4 | 6.0 ± 0.6 | 5.6 ± 0.3 | 6.0 ± 0.8 | 4.3 ± 0.3 | 4.7 ± 0.2 | < 0.001 | t<sub>1</sub> vs. t<sub>6</sub>: p < 0.001, t<sub>1</sub> vs. t<sub>7</sub>: p = 0.045, t<sub>2</sub> vs. t<sub>6</sub>: p < 0.001, t<sub>2</sub> vs. t<sub>7</sub>: p = 0.011, t<sub>3</sub> vs. t<sub>6</sub>: p = 0.001, t<sub>4</sub> vs. t<sub>6</sub>: p = 0.027, t<sub>5</sub> vs. t<sub>6</sub>: p = 0.001 |
| Preparasubthalamic nucleus | 100 | 356 | 5.7 ± 0.3 | 6.5 ± 0.5 | 6.0 ± 0.9 | 5.2 ± 1.0 | 5.9 ± 0.7 | 5.4 ± 0.7 | 4.6 ± 0.6 | < 0.001 | t<sub>2</sub> vs. t<sub>4</sub>: p = 0.008, t<sub>2</sub> vs. t<sub>6</sub>: p = 0.044, t<sub>2</sub> vs. t<sub>7</sub>: p = 0.001 |
| Parasubthalamic nucleus | 101 | 364 | 5.6 ± 0.4 | 6.5 ± 0.4 | 5.9 ± 0.8 | 5.0 ± 0.9 | 5.9 ± 0.9 | 5.6 ± 0.7 | 5.0 ± 0.9 | = 0.002 | t<sub>2</sub> vs. t<sub>4</sub>: p < 0.001, t<sub>2</sub> vs. t<sub>7</sub>: p = 0.010 |
| Subthalamic nucleus | 102 | 470 | 6.0 ± 0.3 | 6.7 ± 0.6 | 6.3 ± 0.6 | 5.7 ± 0.8 | 6.1 ± 0.7 | 6.4 ± 0.7 | 5.8 ± 0.4 | = 0.012 | t<sub>2</sub> vs. t<sub>4</sub>: p = 0.004 |
| Zona incerta | 103 | 797 | 5.3 ± 0.3 | 5.4 ± 0.4 | 5.6 ± 0.4 | 5.2 ± 0.4 | 5.5 ± 0.7 | 4.9 ± 0.3 | 4.8 ± 0.2 | = 0.001 | t<sub>2</sub> vs. t<sub>7</sub>: p = 0.049, t<sub>3</sub> vs. t<sub>6</sub>: p = 0.005, t<sub>3</sub> vs. t<sub>7</sub>: p = 0.005 |
| optic nerve | 104 | 848 | 4.3 ± 0.4 | 4.0 ± 0.5 | 3.7 ± 0.5 | 4.2 ± 0.5 | 4.0 ± 0.5 | 4.3 ± 1.2 | 3.8 ± 0.3 | = 0.124 |  |
| corpus callosum anterior forceps | 105 | 956 | 3.9 ± 0.3 | 4.2 ± 0.3 | 4.3 ± 0.1 | 4.0 ± 0.6 | 4.3 ± 0.2 | 4.3 ± 0.4 | 3.9 ± 0.3 | = 0.074 |  |
| corpus callosum extreme capsule | 106 | 964 | 3.2 ± 0.2 | 3.7 ± 0.3 | 3.6 ± 0.2 | 3.4 ± 0.2 | 3.7 ± 0.3 | 3.6 ± 0.5 | 3.0 ± 0.2 | < 0.001 | t<sub>1</sub> vs. t<sub>2</sub>: p = 0.003, t<sub>1</sub> vs. t<sub>3</sub>: p = 0.014, t<sub>1</sub> vs. t<sub>5</sub>: p = 0.004, t<sub>2</sub> vs. t<sub>7</sub>: p = 0.003, t<sub>3</sub> vs. t<sub>7</sub>: p = 0.012, t<sub>5</sub> vs. t<sub>7</sub>: p = 0.004 |
| corpus callosum splenium | 107 | 986 | 4.3 ± 0.4 | 4.4 ± 0.2 | 4.2 ± 0.2 | 4.2 ± 0.4 | 4.3 ± 0.2 | 4.2 ± 0.5 | 4.2 ± 0.6 | = 0.850 |  |
| corticospinal tract | 108 | 784 | 5.2 ± 0.2 | 5.4 ± 0.1 | 4.9 ± 0.2 | 5.1 ± 0.3 | 5.2 ± 0.3 | 5.0 ± 0.3 | 5.0 ± 0.2 | = 0.004 | t<sub>2</sub> vs. t<sub>3</sub>: p = 0.003, t<sub>2</sub> vs. t<sub>6</sub>: p = 0.018 |
| thalamus related | 109 | 896 | 5.1 ± 0.3 | 5.2 ± 0.2 | 4.8 ± 0.3 | 5.1 ± 0.3 | 5.2 ± 0.5 | 5.2 ± 0.4 | 5.2 ± 0.2 | = 0.183 |  |
| cerebral nuclei related | 110 | 760 | 5.7 ± 0.3 | 6.3 ± 0.4 | 6.3 ± 0.8 | 5.5 ± 0.7 | 5.7 ± 0.7 | 5.7 ± 0.6 | 5.4 ± 0.6 | = 0.026 | t<sub>2</sub> vs. t<sub>4</sub>: p = 0.031 |
| amygdalar capsule | 111 | 884 | 3.3 ± 0.1 | 3.9 ± 0.2 | 3.9 ± 0.2 | 3.7 ± 0.4 | 4.1 ± 0.2 | 3.8 ± 0.4 | 3.3 ± 0.3 | < 0.001 | t<sub>1</sub> vs. t<sub>2</sub>: p = 0.003, t<sub>1</sub> vs. t<sub>5</sub>: p < 0.001, t<sub>5</sub> vs. t<sub>7</sub>: p = 0.007 |
| anterior commissure temporal limb | 112 | 908 | 4.3 ± 0.4 | 4.2 ± 0.5 | 4.6 ± 0.5 | 4.7 ± 0.7 | 4.5 ± 0.3 | 4.9 ± 0.5 | 4.6 ± 0.6 | = 0.164 |  |
| cingulum bundle | 113 | 940 | 3.9 ± 0.3 | 4.0 ± 0.3 | 3.7 ± 0.2 | 3.5 ± 0.2 | 3.6 ± 0.2 | 3.8 ± 0.4 | 3.8 ± 0.5 | = 0.015 | t<sub>2</sub> vs. t<sub>4</sub>: p = 0.007, t<sub>2</sub> vs. t<sub>5</sub>: p = 0.049 |
| alveus | 114 | 466 | 3.9 ± 0.2 | 4.4 ± 0.2 | 4.1 ± 0.3 | 3.9 ± 0.3 | 4.1 ± 0.2 | 4.3 ± 0.4 | 4.2 ± 0.6 | = 0.006 | t<sub>1</sub> vs. t<sub>2</sub>: p = 0.004, t<sub>2</sub> vs. t<sub>4</sub>: p = 0.010 |
| dorsal fornix | 115 | 530 | 4.6 ± 0.6 | 5.2 ± 0.9 | 4.4 ± 0.5 | 3.7 ± 0.6 | 4.2 ± 0.7 | 3.9 ± 0.8 | 4.1 ± 0.7 | = 0.004 | t<sub>2</sub> vs. t<sub>4</sub>: p = 0.003, t<sub>2</sub> vs. t<sub>6</sub>: p = 0.019 |
| fimbria | 116 | 603 | 4.4 ± 0.2 | 4.4 ± 0.4 | 4.1 ± 0.2 | 4.2 ± 0.2 | 4.2 ± 0.3 | 4.3 ± 0.5 | 4.0 ± 0.4 | = 0.046 | t<sub>1</sub> vs. t<sub>3</sub>: p = 0.034 |
| postcommissural fornix | 117 | 737 | 5.2 ± 0.4 | 5.2 ± 0.5 | 4.8 ± 0.5 | 5.0 ± 0.4 | 5.6 ± 0.8 | 4.1 ± 0.3 | 4.4 ± 0.2 | < 0.001 | t<sub>1</sub> vs. t<sub>6</sub>: p < 0.001, t<sub>2</sub> vs. t<sub>6</sub>: p < 0.001, t<sub>4</sub> vs. t<sub>6</sub>: p = 0.015, t<sub>5</sub> vs. t<sub>6</sub>: p < 0.001, t<sub>5</sub> vs. t<sub>7</sub>: p = 0.009 |
| hippocampal commissures | 118 | 618 | 4.3 ± 0.4 | 4.4 ± 0.5 | 4.1 ± 0.2 | 3.9 ± 0.3 | 4.1 ± 0.4 | 4.0 ± 0.6 | 4.0 ± 0.5 | = 0.024 | t<sub>2</sub> vs. t<sub>4</sub>: p = 0.029 |
| stria terminalis | 119 | 301 | 4.2 ± 0.2 | 4.4 ± 0.2 | 4.2 ± 0.2 | 4.2 ± 0.2 | 4.2 ± 0.2 | 4.1 ± 0.2 | 4.0 ± 0.3 | = 0.077 |  |
| supraoptic commissures | 120 | 349 | 5.2 ± 0.9 | 5.4 ± 0.8 | 4.5 ± 1.0 | 5.3 ± 0.8 | 5.6 ± 0.9 | 5.3 ± 1.5 | 4.6 ± 0.5 | = 0.110 |  |
| mammillary related | 121 | 46 | 5.2 ± 0.5 | 5.5 ± 0.6 | 5.6 ± 0.7 | 5.2 ± 0.5 | 5.8 ± 1.2 | 4.9 ± 0.6 | 5.0 ± 0.5 | = 0.229 |  |
| epithalamus related | 122 | 1083 | 5.2 ± 0.2 | 5.1 ± 0.4 | 4.8 ± 0.4 | 4.6 ± 0.3 | 4.8 ± 0.3 | 4.6 ± 0.3 | 4.3 ± 0.6 | < 0.001 | t<sub>1</sub> vs. t<sub>4</sub>: p = 0.006, t<sub>1</sub> vs. t<sub>6</sub>: p = 0.006, t<sub>1</sub> vs. t<sub>7</sub>: p = 0.011, t<sub>2</sub> vs. t<sub>4</sub>: p = 0.034, t<sub>2</sub> vs. t<sub>6</sub>: p = 0.030, t<sub>2</sub> vs. t<sub>7</sub>: p = 0.045 |
| ROI label | ROI ID | ROI ID Allen | EAdol,<br>*ϕ* in rad | MAdol,<br>*ϕ* in rad | LAdol,<br>*ϕ* in rad | EAdult,<br>*ϕ* in rad | MAdult,<br>*ϕ* in rad | LAdult,<br>*ϕ* in rad | Old,<br>*ϕ* in rad | p | Multiple comparison |
| Primary motor area | 1 | 985 | 0.46 ± 0.05 | 0.50 ± 0.06 | 0.55 ± 0.06 | 0.51 ± 0.11 | 0.50 ± 0.13 | 0.74 ± 0.13 | 0.89 ± 0.31 | < 0.001 | t<sub>1</sub> vs. t<sub>6</sub>: p = 0.001, t<sub>1</sub> vs. t<sub>7</sub>: p = 0.007, t<sub>2</sub> vs. t<sub>7</sub>: p = 0.047, t<sub>4</sub> vs. t<sub>7</sub>: p = 0.027, t<sub>5</sub> vs. t<sub>6</sub>: p = 0.032 |
| Secondary motor area | 2 | 993 | 0.62 ± 0.17 | 0.62 ± 0.15 | 0.73 ± 0.17 | 0.66 ± 0.14 | 0.70 ± 0.25 | 0.82 ± 0.18 | 0.84 ± 0.26 | = 0.116 |  |
| Primary somatosensory area nose | 3 | 353 | 0.60 ± 0.09 | 0.62 ± 0.11 | 0.54 ± 0.09 | 0.55 ± 0.06 | 0.48 ± 0.14 | 0.66 ± 0.13 | 0.68 ± 0.17 | = 0.008 | t<sub>2</sub> vs. t<sub>5</sub>: p = 0.042, t<sub>5</sub> vs. t<sub>6</sub>: p = 0.018, t<sub>5</sub> vs. t<sub>7</sub>: p = 0.028 |
| Primary somatosensory area barrel field layer 1 | 4 | 981 | 0.69 ± 0.10 | 0.71 ± 0.20 | 0.64 ± 0.13 | 0.66 ± 0.11 | 0.70 ± 0.13 | 0.81 ± 0.11 | 0.86 ± 0.13 | = 0.026 |  |
| Primary somatosensory area barrel field layer 2/3 | 5 | 201 | 0.56 ± 0.08 | 0.57 ± 0.18 | 0.51 ± 0.09 | 0.46 ± 0.12 | 0.47 ± 0.14 | 0.78 ± 0.12 | 0.87 ± 0.29 | < 0.001 | t<sub>1</sub> vs. t<sub>6</sub>: p = 0.035, t<sub>3</sub> vs. t<sub>6</sub>: p = 0.002, t<sub>4</sub> vs. t<sub>7</sub>: p = 0.026, t<sub>5</sub> vs. t<sub>6</sub>: p = 0.003, t<sub>5</sub> vs. t<sub>7</sub>: p = 0.036 |
| Primary somatosensory area barrel field layer 4 | 6 | 1047 | 0.40 ± 0.10 | 0.39 ± 0.07 | 0.38 ± 0.05 | 0.32 ± 0.07 | 0.31 ± 0.14 | 0.72 ± 0.24 | 0.70 ± 0.28 | < 0.001 | t<sub>4</sub> vs. t<sub>6</sub>: p < 0.001, t<sub>4</sub> vs. t<sub>7</sub>: p = 0.030, t<sub>5</sub> vs. t<sub>6</sub>: p < 0.001, t<sub>5</sub> vs. t<sub>7</sub>: p = 0.004 |
| Primary somatosensory area barrel field layer 5 | 7 | 1070 | 0.40 ± 0.06 | 0.41 ± 0.09 | 0.40 ± 0.07 | 0.40 ± 0.06 | 0.36 ± 0.12 | 0.64 ± 0.26 | 0.59 ± 0.16 | = 0.013 | t<sub>5</sub> vs. t<sub>6</sub>: p = 0.014, t<sub>5</sub> vs. t<sub>7</sub>: p = 0.032 |
| Primary somatosensory area barrel field layer 6a | 8 | 1038 | 0.42 ± 0.06 | 0.49 ± 0.07 | 0.47 ± 0.10 | 0.52 ± 0.07 | 0.49 ± 0.12 | 0.66 ± 0.10 | 0.72 ± 0.08 | < 0.001 | t<sub>1</sub> vs. t<sub>6</sub>: p < 0.001, t<sub>1</sub> vs. t<sub>7</sub>: p < 0.001, t<sub>2</sub> vs. t<sub>7</sub>: p = 0.049, t<sub>3</sub> vs. t<sub>6</sub>: p = 0.013, t<sub>3</sub> vs. t<sub>7</sub>: p = 0.010, t<sub>5</sub> vs. t<sub>7</sub>: p = 0.039 |
| Primary somatosensory area barrel field layer 6b | 9 | 1062 | 0.43 ± 0.07 | 0.54 ± 0.10 | 0.50 ± 0.11 | 0.55 ± 0.07 | 0.58 ± 0.13 | 0.68 ± 0.08 | 0.71 ± 0.05 | < 0.001 | t<sub>1</sub> vs. t<sub>5</sub>: p = 0.040, t<sub>1</sub> vs. t<sub>6</sub>: p < 0.001, t<sub>1</sub> vs. t<sub>7</sub>: p < 0.001, t<sub>3</sub> vs. t<sub>6</sub>: p = 0.010, t<sub>3</sub> vs. t<sub>7</sub>: p = 0.015 |
| Primary somatosensory area lower limb | 10 | 337 | 0.41 ± 0.06 | 0.44 ± 0.07 | 0.48 ± 0.05 | 0.43 ± 0.08 | 0.42 ± 0.09 | 0.73 ± 0.15 | 0.85 ± 0.32 | < 0.001 | t<sub>1</sub> vs. t<sub>6</sub>: p = 0.002, t<sub>1</sub> vs. t<sub>7</sub>: p = 0.006, t<sub>2</sub> vs. t<sub>6</sub>: p = 0.017, t<sub>2</sub> vs. t<sub>7</sub>: p = 0.039, t<sub>4</sub> vs. t<sub>6</sub>: p = 0.008, t<sub>4</sub> vs. t<sub>7</sub>: p = 0.022, t<sub>5</sub> vs. t<sub>6</sub>: p = 0.002, t<sub>5</sub> vs. t<sub>7</sub>: p = 0.008 |
| Primary somatosensory area mouth | 11 | 345 | 0.51 ± 0.07 | 0.54 ± 0.10 | 0.60 ± 0.08 | 0.57 ± 0.08 | 0.59 ± 0.11 | 0.70 ± 0.06 | 0.78 ± 0.09 | < 0.001 | t<sub>1</sub> vs. t<sub>6</sub>: p = 0.001, t<sub>1</sub> vs. t<sub>7</sub>: p < 0.001, t<sub>2</sub> vs. t<sub>6</sub>: p = 0.020, t<sub>2</sub> vs. t<sub>7</sub>: p = 0.008, t<sub>4</sub> vs. t<sub>7</sub>: p = 0.036 |
| Primary somatosensory area upper limb | 12 | 369 | 0.44 ± 0.10 | 0.45 ± 0.09 | 0.47 ± 0.06 | 0.43 ± 0.06 | 0.41 ± 0.12 | 0.66 ± 0.13 | 0.76 ± 0.20 | < 0.001 | t<sub>1</sub> vs. t<sub>6</sub>: p = 0.021, t<sub>1</sub> vs. t<sub>7</sub>: p = 0.026, t<sub>4</sub> vs. t<sub>6</sub>: p = 0.028, t<sub>4</sub> vs. t<sub>7</sub>: p = 0.034, t<sub>5</sub> vs. t<sub>6</sub>: p = 0.002, t<sub>5</sub> vs. t<sub>7</sub>: p = 0.004 |
| Primary somatosensory area trunk | 13 | 361 | 0.36 ± 0.08 | 0.39 ± 0.09 | 0.40 ± 0.06 | 0.38 ± 0.09 | 0.40 ± 0.11 | 0.79 ± 0.17 | 0.81 ± 0.29 | < 0.001 | t<sub>1</sub> vs. t<sub>6</sub>: p < 0.001, t<sub>1</sub> vs. t<sub>7</sub>: p = 0.011, t<sub>2</sub> vs. t<sub>6</sub>: p = 0.005, t<sub>2</sub> vs. t<sub>7</sub>: p = 0.041, t<sub>3</sub> vs. t<sub>6</sub>: p = 0.019, t<sub>4</sub> vs. t<sub>6</sub>: p = 0.002, t<sub>4</sub> vs. t<sub>7</sub>: p = 0.020, t<sub>5</sub> vs. t<sub>6</sub>: p = 0.006, t<sub>5</sub> vs. t<sub>7</sub>: p = 0.049 |
| Supplemental somatosensory area | 14 | 378 | 0.59 ± 0.08 | 0.65 ± 0.07 | 0.63 ± 0.09 | 0.55 ± 0.09 | 0.55 ± 0.14 | 0.62 ± 0.10 | 0.66 ± 0.15 | = 0.095 |  |
| Gustatory areas | 15 | 1057 | 0.57 ± 0.09 | 0.58 ± 0.10 | 0.61 ± 0.14 | 0.48 ± 0.12 | 0.48 ± 0.12 | 0.53 ± 0.09 | 0.47 ± 0.08 | = 0.035 |  |
| Visceral area | 16 | 677 | 0.56 ± 0.10 | 0.58 ± 0.10 | 0.55 ± 0.09 | 0.49 ± 0.07 | 0.47 ± 0.09 | 0.56 ± 0.09 | 0.61 ± 0.11 | = 0.050 |  |
| Dorsal auditory area layer 1 | 17 | 527 | 0.74 ± 0.19 | 0.81 ± 0.26 | 0.84 ± 0.22 | 0.76 ± 0.17 | 0.85 ± 0.16 | 0.92 ± 0.11 | 0.89 ± 0.28 | = 0.304 |  |
| Dorsal auditory area layer 2/3 | 18 | 600 | 0.58 ± 0.17 | 0.65 ± 0.17 | 0.65 ± 0.22 | 0.56 ± 0.16 | 0.56 ± 0.13 | 0.79 ± 0.20 | 0.87 ± 0.27 | = 0.069 |  |
| Dorsal auditory area layer 4 | 19 | 678 | 0.49 ± 0.11 | 0.56 ± 0.10 | 0.57 ± 0.19 | 0.51 ± 0.17 | 0.42 ± 0.13 | 0.60 ± 0.19 | 0.66 ± 0.30 | = 0.179 |  |
| Dorsal auditory area layer 5 | 20 | 252 | 0.49 ± 0.12 | 0.54 ± 0.07 | 0.56 ± 0.09 | 0.55 ± 0.12 | 0.44 ± 0.09 | 0.50 ± 0.13 | 0.56 ± 0.21 | = 0.089 |  |
| Dorsal auditory area layer 6a | 21 | 156 | 0.50 ± 0.12 | 0.55 ± 0.05 | 0.62 ± 0.08 | 0.62 ± 0.11 | 0.56 ± 0.13 | 0.54 ± 0.12 | 0.69 ± 0.11 | = 0.070 |  |
| Dorsal auditory area layer 6b | 22 | 243 | 0.52 ± 0.12 | 0.70 ± 0.20 | 0.81 ± 0.25 | 0.72 ± 0.15 | 0.70 ± 0.21 | 0.65 ± 0.16 | 0.81 ± 0.12 | = 0.012 | t<sub>1</sub> vs. t<sub>3</sub>: p = 0.009, t<sub>1</sub> vs. t<sub>7</sub>: p = 0.014 |
| Ventral auditory area | 23 | 1018 | 0.60 ± 0.13 | 0.74 ± 0.17 | 0.73 ± 0.23 | 0.62 ± 0.18 | 0.66 ± 0.16 | 0.68 ± 0.18 | 0.72 ± 0.24 | = 0.477 |  |
| Anterior cingulate area dorsal part | 24 | 39 | 0.45 ± 0.10 | 0.58 ± 0.13 | 0.53 ± 0.19 | 0.48 ± 0.16 | 0.44 ± 0.12 | 0.57 ± 0.10 | 0.79 ± 0.33 | = 0.028 |  |
| Anterior cingulate area ventral part | 25 | 48 | 0.39 ± 0.08 | 0.47 ± 0.12 | 0.47 ± 0.09 | 0.45 ± 0.15 | 0.43 ± 0.19 | 0.79 ± 0.18 | 0.77 ± 0.14 | < 0.001 | t<sub>1</sub> vs. t<sub>6</sub>: p < 0.001, t<sub>1</sub> vs. t<sub>7</sub>: p = 0.007, t<sub>2</sub> vs. t<sub>6</sub>: p = 0.044, t<sub>4</sub> vs. t<sub>6</sub>: p = 0.015, t<sub>5</sub> vs. t<sub>6</sub>: p = 0.002, t<sub>5</sub> vs. t<sub>7</sub>: p = 0.019 |
| Agranular insular area posterior part | 26 | 111 | 0.47 ± 0.07 | 0.50 ± 0.12 | 0.55 ± 0.14 | 0.60 ± 0.15 | 0.57 ± 0.18 | 0.63 ± 0.08 | 0.72 ± 0.07 | = 0.009 | t<sub>1</sub> vs. t<sub>7</sub>: p = 0.011, t<sub>2</sub> vs. t<sub>7</sub>: p = 0.029 |
| Retrosplenial area lateral agranular part layer 1 | 27 | 671 | 0.58 ± 0.15 | 0.64 ± 0.13 | 0.62 ± 0.10 | 0.52 ± 0.09 | 0.68 ± 0.18 | 0.78 ± 0.09 | 0.63 ± 0.08 | = 0.008 | t<sub>1</sub> vs. t<sub>6</sub>: p = 0.023, t<sub>4</sub> vs. t<sub>6</sub>: p = 0.001 |
| Retrosplenial area lateral agranular part layer 2/3 | 28 | 965 | 0.47 ± 0.12 | 0.54 ± 0.15 | 0.52 ± 0.05 | 0.47 ± 0.09 | 0.54 ± 0.19 | 0.85 ± 0.18 | 0.79 ± 0.17 | < 0.001 | t<sub>1</sub> vs. t<sub>6</sub>: p = 0.001, t<sub>1</sub> vs. t<sub>7</sub>: p = 0.017, t<sub>3</sub> vs. t<sub>6</sub>: p = 0.041, t<sub>4</sub> vs. t<sub>6</sub>: p = 0.001, t<sub>4</sub> vs. t<sub>7</sub>: p = 0.016, t<sub>5</sub> vs. t<sub>6</sub>: p = 0.017 |
| Retrosplenial area lateral agranular part layer 5 | 29 | 774 | 0.44 ± 0.13 | 0.41 ± 0.09 | 0.46 ± 0.05 | 0.49 ± 0.09 | 0.45 ± 0.19 | 0.82 ± 0.23 | 0.92 ± 0.34 | < 0.001 | t<sub>1</sub> vs. t<sub>6</sub>: p = 0.013, t<sub>1</sub> vs. t<sub>7</sub>: p = 0.020, t<sub>2</sub> vs. t<sub>6</sub>: p = 0.004, t<sub>2</sub> vs. t<sub>7</sub>: p = 0.007, t<sub>5</sub> vs. t<sub>6</sub>: p = 0.012, t<sub>5</sub> vs. t<sub>7</sub>: p = 0.019 |
| Retrosplenial area lateral agranular part layer 6a | 30 | 906 | 0.48 ± 0.09 | 0.41 ± 0.07 | 0.46 ± 0.05 | 0.50 ± 0.09 | 0.44 ± 0.11 | 0.69 ± 0.18 | 0.78 ± 0.15 | < 0.001 | t<sub>1</sub> vs. t<sub>7</sub>: p < 0.001, t<sub>2</sub> vs. t<sub>7</sub>: p < 0.001, t<sub>3</sub> vs. t<sub>6</sub>: p = 0.042, t<sub>3</sub> vs. t<sub>7</sub>: p = 0.023, t<sub>5</sub> vs. t<sub>6</sub>: p = 0.007, t<sub>5</sub> vs. t<sub>7</sub>: p = 0.004 |
| Retrosplenial area lateral agranular part layer 6b | 31 | 279 | 0.47 ± 0.08 | 0.40 ± 0.07 | 0.44 ± 0.05 | 0.48 ± 0.06 | 0.43 ± 0.06 | 0.62 ± 0.10 | 0.73 ± 0.12 | < 0.001 | t<sub>1</sub> vs. t<sub>7</sub>: p = 0.049, t<sub>2</sub> vs. t<sub>6</sub>: p < 0.001, t<sub>2</sub> vs. t<sub>7</sub>: p < 0.001, t<sub>3</sub> vs. t<sub>6</sub>: p = 0.016, t<sub>3</sub> vs. t<sub>7</sub>: p = 0.009, t<sub>5</sub> vs. t<sub>6</sub>: p = 0.003, t<sub>5</sub> vs. t<sub>7</sub>: p = 0.002 |
| Retrosplenial area dorsal part | 32 | 879 | 0.42 ± 0.13 | 0.43 ± 0.17 | 0.42 ± 0.09 | 0.37 ± 0.08 | 0.42 ± 0.18 | 0.82 ± 0.22 | 0.80 ± 0.27 | < 0.001 | t<sub>1</sub> vs. t<sub>6</sub>: p = 0.035, t<sub>2</sub> vs. t<sub>6</sub>: p = 0.012, t<sub>3</sub> vs. t<sub>6</sub>: p = 0.049, t<sub>4</sub> vs. t<sub>6</sub>: p = 0.001, t<sub>4</sub> vs. t<sub>7</sub>: p = 0.014, t<sub>5</sub> vs. t<sub>6</sub>: p = 0.006, t<sub>5</sub> vs. t<sub>7</sub>: p = 0.049 |
| Retrosplenial area ventral part | 33 | 886 | 0.43 ± 0.09 | 0.49 ± 0.12 | 0.49 ± 0.08 | 0.51 ± 0.11 | 0.56 ± 0.17 | 0.69 ± 0.18 | 0.75 ± 0.09 | < 0.001 | t<sub>1</sub> vs. t<sub>6</sub>: p = 0.004, t<sub>1</sub> vs. t<sub>7</sub>: p = 0.002, t<sub>2</sub> vs. t<sub>7</sub>: p = 0.032, t<sub>3</sub> vs. t<sub>7</sub>: p = 0.024 |
| Temporal association areas | 34 | 541 | 0.52 ± 0.11 | 0.60 ± 0.10 | 0.62 ± 0.09 | 0.56 ± 0.08 | 0.54 ± 0.10 | 0.49 ± 0.08 | 0.48 ± 0.10 | = 0.035 | t<sub>3</sub> vs. t<sub>6</sub>: p = 0.038 |
| Perirhinal area | 35 | 922 | 0.63 ± 0.08 | 0.61 ± 0.10 | 0.61 ± 0.09 | 0.70 ± 0.14 | 0.78 ± 0.12 | 0.64 ± 0.07 | 0.64 ± 0.09 | = 0.038 | t<sub>2</sub> vs. t<sub>5</sub>: p = 0.031, t<sub>3</sub> vs. t<sub>5</sub>: p = 0.036 |
| Ectorhinal area | 36 | 895 | 0.52 ± 0.07 | 0.52 ± 0.12 | 0.59 ± 0.08 | 0.61 ± 0.12 | 0.55 ± 0.15 | 0.53 ± 0.08 | 0.54 ± 0.16 | = 0.412 |  |
| Piriform area | 37 | 961 | 0.54 ± 0.11 | 0.49 ± 0.17 | 0.58 ± 0.10 | 0.60 ± 0.13 | 0.61 ± 0.14 | 0.64 ± 0.15 | 0.79 ± 0.15 | = 0.020 | t<sub>2</sub> vs. t<sub>7</sub>: p = 0.005 |
| Field CA1 | 38 | 382 | 0.64 ± 0.16 | 0.77 ± 0.26 | 0.79 ± 0.18 | 0.79 ± 0.18 | 0.81 ± 0.22 | 0.91 ± 0.24 | 0.81 ± 0.16 | = 0.253 |  |
| Field CA2 | 39 | 423 | 0.58 ± 0.09 | 0.63 ± 0.11 | 0.54 ± 0.08 | 0.58 ± 0.09 | 0.56 ± 0.11 | 0.94 ± 0.13 | 0.90 ± 0.13 | < 0.001 | t<sub>1</sub> vs. t<sub>6</sub>: p = 0.008, t<sub>3</sub> vs. t<sub>6</sub>: p < 0.001, t<sub>3</sub> vs. t<sub>7</sub>: p = 0.004, t<sub>4</sub> vs. t<sub>6</sub>: p = 0.005, t<sub>4</sub> vs. t<sub>7</sub>: p = 0.038, t<sub>5</sub> vs. t<sub>6</sub>: p < 0.001, t<sub>5</sub> vs. t<sub>7</sub>: p = 0.007 |
| Field CA3 | 40 | 463 | 0.74 ± 0.16 | 0.71 ± 0.12 | 0.62 ± 0.07 | 0.72 ± 0.12 | 0.67 ± 0.11 | 0.92 ± 0.06 | 0.97 ± 0.19 | < 0.001 | t<sub>3</sub> vs. t<sub>6</sub>: p < 0.001, t<sub>3</sub> vs. t<sub>7</sub>: p = 0.003, t<sub>4</sub> vs. t<sub>6</sub>: p = 0.042, t<sub>5</sub> vs. t<sub>6</sub>: p = 0.007, t<sub>5</sub> vs. t<sub>7</sub>: p = 0.043 |
| Dentate gyrus molecular layer | 41 | 10703 | 0.70 ± 0.09 | 0.59 ± 0.08 | 0.56 ± 0.06 | 0.61 ± 0.07 | 0.53 ± 0.09 | 0.72 ± 0.12 | 0.74 ± 0.20 | = 0.001 | t<sub>1</sub> vs. t<sub>5</sub>: p = 0.042, t<sub>3</sub> vs. t<sub>6</sub>: p = 0.009, t<sub>5</sub> vs. t<sub>6</sub>: p = 0.012 |
| Dentate gyrus polymorph layer | 42 | 10704 | 0.58 ± 0.07 | 0.51 ± 0.09 | 0.49 ± 0.04 | 0.54 ± 0.07 | 0.47 ± 0.09 | 0.67 ± 0.16 | 0.69 ± 0.23 | = 0.004 | t<sub>3</sub> vs. t<sub>6</sub>: p = 0.049, t<sub>5</sub> vs. t<sub>6</sub>: p = 0.019 |
| Dentate gyrus granule cell layer | 43 | 632 | 0.59 ± 0.07 | 0.51 ± 0.08 | 0.50 ± 0.04 | 0.55 ± 0.06 | 0.47 ± 0.08 | 0.64 ± 0.14 | 0.68 ± 0.21 | = 0.004 | t<sub>1</sub> vs. t<sub>5</sub>: p = 0.032, t<sub>5</sub> vs. t<sub>6</sub>: p = 0.027 |
| Fasciola cinerea | 44 | 982 | 0.45 ± 0.12 | 0.45 ± 0.11 | 0.65 ± 0.15 | 0.54 ± 0.10 | 0.54 ± 0.11 | 0.56 ± 0.19 | 0.54 ± 0.16 | = 0.033 | t<sub>1</sub> vs. t<sub>3</sub>: p = 0.022, t<sub>2</sub> vs. t<sub>3</sub>: p = 0.025 |
| Induseum griseum | 45 | 19 | 0.43 ± 0.10 | 0.51 ± 0.22 | 0.39 ± 0.10 | 0.37 ± 0.11 | 0.44 ± 0.20 | 0.87 ± 0.15 | 1.05 ± 0.33 | < 0.001 | t<sub>1</sub> vs. t<sub>6</sub>: p = 0.026, t<sub>1</sub> vs. t<sub>7</sub>: p = 0.042, t<sub>3</sub> vs. t<sub>6</sub>: p = 0.003, t<sub>3</sub> vs. t<sub>7</sub>: p = 0.008, t<sub>4</sub> vs. t<sub>6</sub>: p < 0.001, t<sub>4</sub> vs. t<sub>7</sub>: p = 0.003, t<sub>5</sub> vs. t<sub>6</sub>: p = 0.008, t<sub>5</sub> vs. t<sub>7</sub>: p = 0.017 |
| Entorhinal area lateral part | 46 | 918 | 0.52 ± 0.09 | 0.52 ± 0.11 | 0.53 ± 0.11 | 0.58 ± 0.11 | 0.63 ± 0.09 | 0.64 ± 0.12 | 0.70 ± 0.18 | = 0.050 |  |
| Prosubiculum ventral part | 47 | 484682492 | 0.42 ± 0.04 | 0.48 ± 0.05 | 0.51 ± 0.07 | 0.52 ± 0.05 | 0.51 ± 0.08 | 0.71 ± 0.06 | 0.75 ± 0.03 | < 0.001 | t<sub>1</sub> vs. t<sub>6</sub>: p < 0.001, t<sub>1</sub> vs. t<sub>7</sub>: p < 0.001, t<sub>2</sub> vs. t<sub>6</sub>: p = 0.002, t<sub>2</sub> vs. t<sub>7</sub>: p = 0.006, t<sub>3</sub> vs. t<sub>6</sub>: p = 0.024, t<sub>4</sub> vs. t<sub>6</sub>: p = 0.046, t<sub>5</sub> vs. t<sub>6</sub>: p = 0.036 |
| Claustrum | 48 | 583 | 0.57 ± 0.14 | 0.58 ± 0.12 | 0.55 ± 0.15 | 0.46 ± 0.13 | 0.45 ± 0.10 | 0.46 ± 0.09 | 0.39 ± 0.14 | = 0.020 |  |
| Endopiriform nucleus | 49 | 942 | 0.44 ± 0.06 | 0.52 ± 0.08 | 0.47 ± 0.13 | 0.45 ± 0.08 | 0.44 ± 0.09 | 0.56 ± 0.11 | 0.50 ± 0.12 | = 0.038 |  |
| Lateral amygdalar nucleus | 50 | 131 | 0.52 ± 0.10 | 0.54 ± 0.07 | 0.52 ± 0.15 | 0.55 ± 0.16 | 0.53 ± 0.17 | 0.75 ± 0.20 | 0.74 ± 0.22 | = 0.075 |  |
| Basolateral amygdalar nucleus | 51 | 295 | 0.42 ± 0.10 | 0.54 ± 0.07 | 0.49 ± 0.13 | 0.58 ± 0.11 | 0.59 ± 0.19 | 0.64 ± 0.16 | 0.53 ± 0.16 | = 0.042 | t<sub>1</sub> vs. t<sub>6</sub>: p = 0.021 |
| Basomedial amygdalar nucleus | 52 | 319 | 0.52 ± 0.13 | 0.58 ± 0.07 | 0.48 ± 0.09 | 0.62 ± 0.12 | 0.58 ± 0.14 | 0.73 ± 0.19 | 0.68 ± 0.15 | = 0.020 | t<sub>3</sub> vs. t<sub>6</sub>: p = 0.010 |
| Posterior amygdalar nucleus | 53 | 780 | 0.54 ± 0.12 | 0.66 ± 0.16 | 0.56 ± 0.21 | 0.56 ± 0.12 | 0.66 ± 0.09 | 0.84 ± 0.26 | 0.73 ± 0.13 | = 0.011 |  |
| Striatum dorsal region | 54 | 485 | 0.84 ± 0.12 | 0.88 ± 0.10 | 0.82 ± 0.14 | 0.78 ± 0.09 | 0.84 ± 0.17 | 0.93 ± 0.07 | 0.97 ± 0.10 | = 0.046 | t<sub>4</sub> vs. t<sub>7</sub>: p = 0.035 |
| Fundus of striatum | 55 | 998 | 0.52 ± 0.06 | 0.58 ± 0.10 | 0.44 ± 0.08 | 0.55 ± 0.20 | 0.52 ± 0.09 | 0.76 ± 0.07 | 0.73 ± 0.09 | < 0.001 | t<sub>1</sub> vs. t<sub>6</sub>: p = 0.009, t<sub>3</sub> vs. t<sub>6</sub>: p < 0.001, t<sub>3</sub> vs. t<sub>7</sub>: p = 0.002, t<sub>4</sub> vs. t<sub>6</sub>: p = 0.042, t<sub>5</sub> vs. t<sub>6</sub>: p = 0.006 |
| Lateral septal nucleus | 56 | 242 | 0.71 ± 0.18 | 0.75 ± 0.13 | 0.60 ± 0.15 | 0.56 ± 0.14 | 0.61 ± 0.19 | 0.86 ± 0.15 | 0.86 ± 0.14 | = 0.002 | t<sub>3</sub> vs. t<sub>6</sub>: p = 0.041, t<sub>4</sub> vs. t<sub>6</sub>: p = 0.009, t<sub>4</sub> vs. t<sub>7</sub>: p = 0.037, t<sub>5</sub> vs. t<sub>6</sub>: p = 0.048 |
| Septofimbrial nucleus | 57 | 310 | 0.76 ± 0.11 | 0.74 ± 0.08 | 0.69 ± 0.11 | 0.77 ± 0.12 | 0.71 ± 0.15 | 0.82 ± 0.12 | 0.89 ± 0.16 | = 0.063 |  |
| Anterior amygdalar area | 58 | 23 | 0.48 ± 0.08 | 0.53 ± 0.10 | 0.42 ± 0.10 | 0.53 ± 0.22 | 0.50 ± 0.10 | 0.86 ± 0.08 | 0.90 ± 0.06 | < 0.001 | t<sub>1</sub> vs. t<sub>6</sub>: p = 0.008, t<sub>1</sub> vs. t<sub>7</sub>: p = 0.020, t<sub>3</sub> vs. t<sub>6</sub>: p < 0.001, t<sub>3</sub> vs. t<sub>7</sub>: p < 0.001, t<sub>4</sub> vs. t<sub>6</sub>: p = 0.021, t<sub>4</sub> vs. t<sub>7</sub>: p = 0.045, t<sub>5</sub> vs. t<sub>6</sub>: p = 0.009, t<sub>5</sub> vs. t<sub>7</sub>: p = 0.023 |
| Central amygdalar nucleus | 59 | 536 | 0.56 ± 0.10 | 0.50 ± 0.08 | 0.52 ± 0.10 | 0.50 ± 0.13 | 0.46 ± 0.11 | 0.74 ± 0.08 | 0.78 ± 0.14 | < 0.001 | t<sub>2</sub> vs. t<sub>6</sub>: p = 0.009, t<sub>2</sub> vs. t<sub>7</sub>: p = 0.031, t<sub>3</sub> vs. t<sub>6</sub>: p = 0.022, t<sub>4</sub> vs. t<sub>6</sub>: p = 0.005, t<sub>4</sub> vs. t<sub>7</sub>: p = 0.021, t<sub>5</sub> vs. t<sub>6</sub>: p < 0.001, t<sub>5</sub> vs. t<sub>7</sub>: p = 0.003 |
| Intercalated amygdalar nucleus | 60 | 1105 | 0.46 ± 0.07 | 0.52 ± 0.06 | 0.42 ± 0.07 | 0.47 ± 0.10 | 0.45 ± 0.10 | 0.68 ± 0.22 | 0.65 ± 0.16 | = 0.004 | t<sub>3</sub> vs. t<sub>6</sub>: p = 0.026, t<sub>3</sub> vs. t<sub>7</sub>: p = 0.017 |
| Pallidum dorsal region | 61 | 818 | 0.87 ± 0.13 | 0.90 ± 0.27 | 0.87 ± 0.12 | 0.82 ± 0.13 | 0.94 ± 0.19 | 1.01 ± 0.14 | 1.05 ± 0.23 | = 0.144 |  |
| Pallidum ventral region | 62 | 835 | 0.65 ± 0.21 | 0.76 ± 0.17 | 0.62 ± 0.15 | 0.56 ± 0.11 | 0.75 ± 0.23 | 0.93 ± 0.20 | 0.88 ± 0.14 | = 0.001 | t<sub>3</sub> vs. t<sub>6</sub>: p = 0.022, t<sub>4</sub> vs. t<sub>6</sub>: p = 0.001, t<sub>4</sub> vs. t<sub>7</sub>: p = 0.018 |
| Triangular nucleus of septum | 63 | 581 | 0.75 ± 0.06 | 0.70 ± 0.07 | 0.69 ± 0.09 | 0.77 ± 0.11 | 0.68 ± 0.14 | 0.79 ± 0.10 | 0.85 ± 0.15 | = 0.044 |  |
| Ventral anterior-lateral complex of the thalamus | 64 | 629 | 1.16 ± 0.11 | 1.04 ± 0.18 | 0.83 ± 0.20 | 0.98 ± 0.28 | 0.97 ± 0.21 | 0.92 ± 0.10 | 0.96 ± 0.10 | = 0.012 | t<sub>1</sub> vs. t<sub>3</sub>: p = 0.001 |
| Ventral medial nucleus of the thalamus | 65 | 685 | 1.17 ± 0.15 | 0.95 ± 0.19 | 1.00 ± 0.17 | 1.12 ± 0.14 | 1.18 ± 0.28 | 1.16 ± 0.10 | 1.08 ± 0.18 | = 0.043 |  |
| Ventral posterior complex of the thalamus | 66 | 709 | 1.17 ± 0.25 | 1.20 ± 0.18 | 1.00 ± 0.26 | 1.03 ± 0.18 | 1.26 ± 0.20 | 1.09 ± 0.16 | 1.11 ± 0.15 | = 0.126 |  |
| Subparafascicular nucleus | 67 | 406 | 0.98 ± 0.16 | 0.97 ± 0.27 | 0.80 ± 0.22 | 0.84 ± 0.29 | 0.78 ± 0.25 | 0.99 ± 0.12 | 0.92 ± 0.16 | = 0.263 |  |
| Subparafascicular area | 68 | 609 | 0.81 ± 0.11 | 0.96 ± 0.33 | 0.72 ± 0.24 | 0.77 ± 0.29 | 0.74 ± 0.24 | 0.89 ± 0.17 | 0.84 ± 0.07 | = 0.400 |  |
| Dorsal part of the lateral geniculate complex | 69 | 170 | 0.69 ± 0.06 | 0.68 ± 0.14 | 0.73 ± 0.11 | 0.71 ± 0.08 | 0.73 ± 0.07 | 0.68 ± 0.10 | 0.71 ± 0.07 | = 0.835 |  |
| Lateral group of the dorsal thalamus | 70 | 138 | 1.40 ± 0.18 | 1.44 ± 0.13 | 1.06 ± 0.17 | 1.11 ± 0.20 | 1.37 ± 0.24 | 1.07 ± 0.10 | 1.13 ± 0.18 | < 0.001 | t<sub>1</sub> vs. t<sub>3</sub>: p = 0.006, t<sub>1</sub> vs. t<sub>6</sub>: p = 0.015, t<sub>2</sub> vs. t<sub>3</sub>: p = 0.001, t<sub>2</sub> vs. t<sub>4</sub>: p = 0.020, t<sub>2</sub> vs. t<sub>6</sub>: p = 0.004, t<sub>3</sub> vs. t<sub>5</sub>: p = 0.021, t<sub>5</sub> vs. t<sub>6</sub>: p = 0.046 |
| Anteroventral nucleus of thalamus | 71 | 255 | 1.37 ± 0.17 | 1.21 ± 0.15 | 1.23 ± 0.24 | 1.51 ± 0.35 | 1.59 ± 0.34 | 1.05 ± 0.13 | 1.19 ± 0.15 | < 0.001 | t<sub>1</sub> vs. t<sub>6</sub>: p = 0.024, t<sub>4</sub> vs. t<sub>6</sub>: p = 0.003, t<sub>5</sub> vs. t<sub>6</sub>: p < 0.001 |
| Anteromedial nucleus | 72 | 127 | 1.06 ± 0.08 | 1.09 ± 0.19 | 0.86 ± 0.09 | 0.88 ± 0.23 | 0.97 ± 0.15 | 1.27 ± 0.13 | 1.19 ± 0.15 | < 0.001 | t<sub>3</sub> vs. t<sub>6</sub>: p < 0.001, t<sub>3</sub> vs. t<sub>7</sub>: p = 0.011, t<sub>4</sub> vs. t<sub>6</sub>: p = 0.001, t<sub>5</sub> vs. t<sub>6</sub>: p = 0.012 |
| Anterodorsal nucleus | 73 | 64 | 1.26 ± 0.15 | 1.12 ± 0.12 | 1.09 ± 0.11 | 1.24 ± 0.22 | 1.13 ± 0.13 | 1.10 ± 0.06 | 1.10 ± 0.11 | = 0.026 |  |
| Interanteromedial nucleus of the thalamus | 74 | 1120 | 0.82 ± 0.17 | 0.90 ± 0.24 | 0.49 ± 0.13 | 0.57 ± 0.23 | 0.67 ± 0.21 | 0.93 ± 0.13 | 0.93 ± 0.12 | < 0.001 | t<sub>2</sub> vs. t<sub>3</sub>: p = 0.016, t<sub>3</sub> vs. t<sub>6</sub>: p = 0.003, t<sub>3</sub> vs. t<sub>7</sub>: p = 0.001, t<sub>4</sub> vs. t<sub>6</sub>: p = 0.009, t<sub>4</sub> vs. t<sub>7</sub>: p = 0.024 |
| Interanterodorsal nucleus of the thalamus | 75 | 1113 | 1.18 ± 0.13 | 1.14 ± 0.11 | 0.94 ± 0.17 | 1.05 ± 0.19 | 1.04 ± 0.16 | 1.23 ± 0.18 | 1.06 ± 0.12 | = 0.009 | t<sub>1</sub> vs. t<sub>3</sub>: p = 0.019, t<sub>3</sub> vs. t<sub>6</sub>: p = 0.007 |
| Lateral dorsal nucleus of thalamus | 76 | 155 | 1.05 ± 0.14 | 1.00 ± 0.25 | 1.07 ± 0.19 | 1.07 ± 0.28 | 1.13 ± 0.28 | 0.90 ± 0.10 | 0.83 ± 0.12 | = 0.121 |  |
| Intermediodorsal nucleus of the thalamus | 77 | 59 | 1.30 ± 0.17 | 1.47 ± 0.14 | 1.12 ± 0.11 | 1.30 ± 0.23 | 1.33 ± 0.10 | 1.03 ± 0.19 | 0.97 ± 0.08 | < 0.001 | t<sub>2</sub> vs. t<sub>3</sub>: p = 0.001, t<sub>2</sub> vs. t<sub>6</sub>: p < 0.001, t<sub>2</sub> vs. t<sub>7</sub>: p < 0.001, t<sub>4</sub> vs. t<sub>7</sub>: p = 0.026, t<sub>5</sub> vs. t<sub>6</sub>: p = 0.033, t<sub>5</sub> vs. t<sub>7</sub>: p = 0.014 |
| Mediodorsal nucleus of thalamus | 78 | 362 | 1.27 ± 0.19 | 1.29 ± 0.10 | 1.09 ± 0.19 | 1.31 ± 0.20 | 1.37 ± 0.23 | 1.60 ± 0.19 | 1.50 ± 0.16 | < 0.001 | t<sub>1</sub> vs. t<sub>6</sub>: p = 0.023, t<sub>3</sub> vs. t<sub>6</sub>: p < 0.001, t<sub>3</sub> vs. t<sub>7</sub>: p = 0.020 |
| Submedial nucleus of the thalamus | 79 | 366 | 0.74 ± 0.22 | 0.70 ± 0.37 | 0.69 ± 0.36 | 0.60 ± 0.25 | 0.62 ± 0.24 | 1.15 ± 0.20 | 0.95 ± 0.21 | = 0.006 | t<sub>2</sub> vs. t<sub>6</sub>: p = 0.046, t<sub>3</sub> vs. t<sub>6</sub>: p = 0.040, t<sub>4</sub> vs. t<sub>6</sub>: p = 0.005, t<sub>5</sub> vs. t<sub>6</sub>: p = 0.008 |
| Perireunensis nucleus | 80 | 1077 | 0.99 ± 0.20 | 0.89 ± 0.17 | 1.00 ± 0.23 | 0.92 ± 0.18 | 1.07 ± 0.24 | 1.07 ± 0.12 | 1.07 ± 0.16 | = 0.288 |  |
| Midline group of the dorsal thalamus | 81 | 571 | 1.04 ± 0.17 | 1.02 ± 0.25 | 0.68 ± 0.13 | 0.77 ± 0.27 | 0.85 ± 0.12 | 0.83 ± 0.07 | 0.82 ± 0.14 | = 0.002 | t<sub>1</sub> vs. t<sub>3</sub>: p < 0.001, t<sub>2</sub> vs. t<sub>3</sub>: p = 0.006 |
| Intralaminar nuclei of the dorsal thalamus | 82 | 51 | 0.65 ± 0.14 | 0.86 ± 0.35 | 0.55 ± 0.20 | 0.58 ± 0.19 | 0.62 ± 0.20 | 0.99 ± 0.25 | 0.86 ± 0.21 | = 0.004 | t<sub>3</sub> vs. t<sub>6</sub>: p = 0.006, t<sub>4</sub> vs. t<sub>6</sub>: p = 0.022 |
| Reticular nucleus of the thalamus | 83 | 262 | 1.15 ± 0.05 | 1.09 ± 0.10 | 1.03 ± 0.16 | 1.08 ± 0.09 | 1.19 ± 0.14 | 1.16 ± 0.07 | 1.21 ± 0.09 | = 0.032 |  |
| Ventral part of the lateral geniculate complex | 84 | 178 | 0.87 ± 0.07 | 0.88 ± 0.18 | 0.86 ± 0.12 | 0.83 ± 0.07 | 0.93 ± 0.11 | 0.85 ± 0.11 | 0.89 ± 0.19 | = 0.595 |  |
| Epithalamus | 85 | 958 | 1.06 ± 0.13 | 0.99 ± 0.20 | 1.12 ± 0.39 | 0.96 ± 0.40 | 1.03 ± 0.40 | 1.04 ± 0.33 | 0.97 ± 0.20 | = 0.884 |  |
| Accessory supraoptic group | 86 | 332 | 0.56 ± 0.11 | 0.59 ± 0.13 | 0.58 ± 0.09 | 0.57 ± 0.09 | 0.52 ± 0.10 | 0.65 ± 0.08 | 0.66 ± 0.09 | = 0.114 |  |
| Periventricular hypothalamic nucleus anterior part | 87 | 30 | 0.74 ± 0.25 | 0.91 ± 0.25 | 0.65 ± 0.17 | 0.70 ± 0.13 | 0.85 ± 0.21 | 0.70 ± 0.21 | 0.61 ± 0.05 | = 0.046 |  |
| Periventricular hypothalamic nucleus intermediate part | 88 | 118 | 0.95 ± 0.29 | 1.06 ± 0.23 | 1.09 ± 0.28 | 1.06 ± 0.16 | 1.18 ± 0.27 | 0.86 ± 0.18 | 0.78 ± 0.06 | = 0.019 | t<sub>5</sub> vs. t<sub>7</sub>: p = 0.027 |
| Dorsomedial nucleus of the hypothalamus | 89 | 830 | 0.61 ± 0.10 | 0.69 ± 0.13 | 0.64 ± 0.11 | 0.64 ± 0.09 | 0.65 ± 0.05 | 0.77 ± 0.09 | 0.76 ± 0.10 | = 0.021 | t<sub>1</sub> vs. t<sub>6</sub>: p = 0.012 |
| Subparaventricular zone | 90 | 347 | 0.70 ± 0.19 | 0.85 ± 0.23 | 0.67 ± 0.09 | 0.72 ± 0.12 | 0.82 ± 0.15 | 0.68 ± 0.13 | 0.60 ± 0.06 | = 0.022 | t<sub>2</sub> vs. t<sub>7</sub>: p = 0.039, t<sub>5</sub> vs. t<sub>7</sub>: p = 0.029 |
| Subfornical organ | 91 | 338 | 0.73 ± 0.12 | 0.55 ± 0.12 | 0.67 ± 0.16 | 0.77 ± 0.15 | 0.70 ± 0.12 | 0.69 ± 0.07 | 0.83 ± 0.18 | = 0.012 | t<sub>1</sub> vs. t<sub>2</sub>: p = 0.026, t<sub>2</sub> vs. t<sub>4</sub>: p = 0.017, t<sub>2</sub> vs. t<sub>7</sub>: p = 0.017 |
| Anterior hypothalamic nucleus | 92 | 88 | 0.64 ± 0.22 | 0.68 ± 0.16 | 0.41 ± 0.11 | 0.50 ± 0.17 | 0.52 ± 0.20 | 0.75 ± 0.20 | 0.70 ± 0.13 | < 0.001 | t<sub>2</sub> vs. t<sub>3</sub>: p = 0.007, t<sub>3</sub> vs. t<sub>6</sub>: p = 0.004, t<sub>3</sub> vs. t<sub>7</sub>: p = 0.029 |
| Supramammillary nucleus | 93 | 525 | 0.63 ± 0.13 | 0.63 ± 0.11 | 0.68 ± 0.17 | 0.60 ± 0.12 | 0.64 ± 0.20 | 0.85 ± 0.24 | 0.84 ± 0.19 | = 0.112 |  |
| Medial preoptic nucleus | 94 | 515 | 0.66 ± 0.10 | 0.78 ± 0.19 | 0.64 ± 0.07 | 0.65 ± 0.13 | 0.69 ± 0.14 | 0.74 ± 0.09 | 0.74 ± 0.15 | = 0.193 |  |
| Dorsal premammillary nucleus | 95 | 980 | 0.47 ± 0.12 | 0.53 ± 0.10 | 0.52 ± 0.12 | 0.49 ± 0.10 | 0.49 ± 0.11 | 0.78 ± 0.26 | 0.83 ± 0.24 | = 0.010 | t<sub>1</sub> vs. t<sub>6</sub>: p = 0.038 |
| Paraventricular hypothalamic nucleus descending division | 96 | 63 | 0.66 ± 0.18 | 0.78 ± 0.23 | 0.69 ± 0.15 | 0.69 ± 0.15 | 0.78 ± 0.15 | 0.77 ± 0.10 | 0.81 ± 0.13 | = 0.377 |  |
| Posterior hypothalamic nucleus | 97 | 946 | 0.73 ± 0.19 | 0.73 ± 0.17 | 0.69 ± 0.14 | 0.68 ± 0.19 | 0.65 ± 0.17 | 0.83 ± 0.19 | 0.79 ± 0.21 | = 0.385 |  |
| Lateral hypothalamic area | 98 | 194 | 0.64 ± 0.17 | 0.77 ± 0.14 | 0.62 ± 0.18 | 0.55 ± 0.09 | 0.70 ± 0.15 | 0.77 ± 0.13 | 0.76 ± 0.16 | = 0.008 | t<sub>2</sub> vs. t<sub>4</sub>: p = 0.011, t<sub>4</sub> vs. t<sub>6</sub>: p = 0.017 |
| Lateral preoptic area | 99 | 226 | 0.60 ± 0.16 | 0.67 ± 0.16 | 0.65 ± 0.13 | 0.53 ± 0.09 | 0.56 ± 0.14 | 0.76 ± 0.11 | 0.84 ± 0.07 | < 0.001 | t<sub>1</sub> vs. t<sub>7</sub>: p = 0.008, t<sub>4</sub> vs. t<sub>7</sub>: p = 0.002, t<sub>5</sub> vs. t<sub>7</sub>: p = 0.017 |
| Preparasubthalamic nucleus | 100 | 356 | 0.69 ± 0.15 | 0.78 ± 0.12 | 0.87 ± 0.35 | 0.65 ± 0.13 | 0.78 ± 0.26 | 0.96 ± 0.16 | 0.99 ± 0.31 | = 0.029 | t<sub>4</sub> vs. t<sub>6</sub>: p = 0.018 |
| Parasubthalamic nucleus | 101 | 364 | 0.63 ± 0.12 | 0.68 ± 0.09 | 0.70 ± 0.18 | 0.57 ± 0.09 | 0.65 ± 0.19 | 1.00 ± 0.18 | 1.08 ± 0.27 | < 0.001 | t<sub>1</sub> vs. t<sub>6</sub>: p = 0.011, t<sub>1</sub> vs. t<sub>7</sub>: p = 0.048, t<sub>4</sub> vs. t<sub>6</sub>: p < 0.001, t<sub>4</sub> vs. t<sub>7</sub>: p = 0.002, t<sub>5</sub> vs. t<sub>6</sub>: p = 0.020 |
| Subthalamic nucleus | 102 | 470 | 0.97 ± 0.16 | 1.04 ± 0.13 | 1.23 ± 0.30 | 1.02 ± 0.14 | 1.16 ± 0.29 | 1.33 ± 0.22 | 1.43 ± 0.17 | = 0.003 | t<sub>1</sub> vs. t<sub>6</sub>: p = 0.025, t<sub>1</sub> vs. t<sub>7</sub>: p = 0.011, t<sub>4</sub> vs. t<sub>7</sub>: p = 0.041 |
| Zona incerta | 103 | 797 | 0.90 ± 0.10 | 0.86 ± 0.13 | 0.89 ± 0.16 | 0.85 ± 0.15 | 0.89 ± 0.16 | 0.97 ± 0.09 | 1.03 ± 0.18 | = 0.213 |  |
| optic nerve | 104 | 848 | 0.77 ± 0.17 | 0.83 ± 0.21 | 0.65 ± 0.12 | 0.63 ± 0.16 | 0.72 ± 0.13 | 0.99 ± 0.31 | 0.82 ± 0.18 | = 0.020 | t<sub>3</sub> vs. t<sub>6</sub>: p = 0.040, t<sub>4</sub> vs. t<sub>6</sub>: p = 0.019 |
| corpus callosum anterior forceps | 105 | 956 | 0.61 ± 0.11 | 0.60 ± 0.09 | 0.59 ± 0.10 | 0.53 ± 0.12 | 0.54 ± 0.10 | 0.59 ± 0.11 | 0.57 ± 0.10 | = 0.490 |  |
| corpus callosum extreme capsule | 106 | 964 | 0.45 ± 0.08 | 0.51 ± 0.10 | 0.47 ± 0.08 | 0.48 ± 0.12 | 0.47 ± 0.09 | 0.59 ± 0.10 | 0.55 ± 0.11 | = 0.074 |  |
| corpus callosum splenium | 107 | 986 | 0.48 ± 0.06 | 0.55 ± 0.06 | 0.54 ± 0.07 | 0.55 ± 0.06 | 0.59 ± 0.10 | 0.70 ± 0.07 | 0.76 ± 0.10 | < 0.001 | t<sub>1</sub> vs. t<sub>6</sub>: p < 0.001, t<sub>1</sub> vs. t<sub>7</sub>: p < 0.001, t<sub>2</sub> vs. t<sub>6</sub>: p = 0.017, t<sub>2</sub> vs. t<sub>7</sub>: p = 0.019, t<sub>3</sub> vs. t<sub>6</sub>: p = 0.017, t<sub>3</sub> vs. t<sub>7</sub>: p = 0.019, t<sub>4</sub> vs. t<sub>6</sub>: p = 0.032, t<sub>4</sub> vs. t<sub>7</sub>: p = 0.032 |
| corticospinal tract | 108 | 784 | 1.12 ± 0.09 | 1.13 ± 0.11 | 1.06 ± 0.16 | 1.04 ± 0.09 | 1.21 ± 0.13 | 1.17 ± 0.08 | 1.27 ± 0.17 | = 0.010 | t<sub>4</sub> vs. t<sub>5</sub>: p = 0.033, t<sub>4</sub> vs. t<sub>7</sub>: p = 0.043 |
| thalamus related | 109 | 896 | 1.21 ± 0.09 | 1.15 ± 0.12 | 1.07 ± 0.22 | 1.12 ± 0.10 | 1.31 ± 0.16 | 1.19 ± 0.08 | 1.21 ± 0.14 | = 0.039 | t<sub>3</sub> vs. t<sub>5</sub>: p = 0.020, t<sub>4</sub> vs. t<sub>5</sub>: p = 0.040 |
| cerebral nuclei related | 110 | 760 | 0.81 ± 0.13 | 0.87 ± 0.14 | 0.97 ± 0.25 | 0.83 ± 0.10 | 0.86 ± 0.18 | 1.01 ± 0.15 | 1.09 ± 0.28 | = 0.097 |  |
| amygdalar capsule | 111 | 884 | 0.46 ± 0.05 | 0.55 ± 0.09 | 0.47 ± 0.09 | 0.48 ± 0.12 | 0.48 ± 0.13 | 0.58 ± 0.09 | 0.52 ± 0.14 | = 0.109 |  |
| anterior commissure temporal limb | 112 | 908 | 0.51 ± 0.07 | 0.59 ± 0.12 | 0.44 ± 0.11 | 0.56 ± 0.28 | 0.50 ± 0.13 | 0.70 ± 0.15 | 0.64 ± 0.13 | = 0.012 | t<sub>3</sub> vs. t<sub>6</sub>: p = 0.007, t<sub>5</sub> vs. t<sub>6</sub>: p = 0.050 |
| cingulum bundle | 113 | 940 | 0.42 ± 0.04 | 0.40 ± 0.06 | 0.44 ± 0.05 | 0.44 ± 0.05 | 0.40 ± 0.08 | 0.64 ± 0.08 | 0.70 ± 0.09 | < 0.001 | t<sub>1</sub> vs. t<sub>6</sub>: p = 0.002, t<sub>1</sub> vs. t<sub>7</sub>: p = 0.004, t<sub>2</sub> vs. t<sub>6</sub>: p < 0.001, t<sub>2</sub> vs. t<sub>7</sub>: p = 0.002, t<sub>3</sub> vs. t<sub>6</sub>: p = 0.039, t<sub>4</sub> vs. t<sub>6</sub>: p = 0.035, t<sub>4</sub> vs. t<sub>7</sub>: p = 0.046, t<sub>5</sub> vs. t<sub>6</sub>: p = 0.002, t<sub>5</sub> vs. t<sub>7</sub>: p = 0.003 |
| alveus | 114 | 466 | 0.52 ± 0.06 | 0.54 ± 0.05 | 0.54 ± 0.07 | 0.58 ± 0.04 | 0.55 ± 0.07 | 0.75 ± 0.07 | 0.77 ± 0.06 | < 0.001 | t<sub>1</sub> vs. t<sub>6</sub>: p < 0.001, t<sub>1</sub> vs. t<sub>7</sub>: p = 0.002, t<sub>2</sub> vs. t<sub>6</sub>: p = 0.002, t<sub>2</sub> vs. t<sub>7</sub>: p = 0.008, t<sub>3</sub> vs. t<sub>6</sub>: p = 0.002, t<sub>3</sub> vs. t<sub>7</sub>: p = 0.007, t<sub>5</sub> vs. t<sub>6</sub>: p = 0.006, t<sub>5</sub> vs. t<sub>7</sub>: p = 0.020 |
| dorsal fornix | 115 | 530 | 0.43 ± 0.10 | 0.53 ± 0.20 | 0.39 ± 0.15 | 0.39 ± 0.06 | 0.45 ± 0.16 | 0.69 ± 0.23 | 0.80 ± 0.25 | = 0.001 | t<sub>3</sub> vs. t<sub>6</sub>: p = 0.008, t<sub>3</sub> vs. t<sub>7</sub>: p = 0.010, t<sub>4</sub> vs. t<sub>6</sub>: p = 0.025, t<sub>4</sub> vs. t<sub>7</sub>: p = 0.026 |
| fimbria | 116 | 603 | 0.88 ± 0.08 | 0.84 ± 0.05 | 0.82 ± 0.06 | 0.84 ± 0.07 | 0.85 ± 0.08 | 0.87 ± 0.04 | 0.87 ± 0.07 | = 0.421 |  |
| postcommissural fornix | 117 | 737 | 0.62 ± 0.15 | 0.68 ± 0.11 | 0.66 ± 0.14 | 0.57 ± 0.08 | 0.65 ± 0.12 | 0.82 ± 0.09 | 0.88 ± 0.12 | < 0.001 | t<sub>1</sub> vs. t<sub>7</sub>: p = 0.037, t<sub>4</sub> vs. t<sub>6</sub>: p = 0.002, t<sub>4</sub> vs. t<sub>7</sub>: p = 0.001 |
| hippocampal commissures | 118 | 618 | 0.55 ± 0.06 | 0.51 ± 0.06 | 0.51 ± 0.08 | 0.57 ± 0.10 | 0.51 ± 0.12 | 0.73 ± 0.10 | 0.79 ± 0.16 | < 0.001 | t<sub>1</sub> vs. t<sub>7</sub>: p = 0.003, t<sub>2</sub> vs. t<sub>7</sub>: p = 0.011, t<sub>3</sub> vs. t<sub>7</sub>: p = 0.004, t<sub>4</sub> vs. t<sub>7</sub>: p = 0.016, t<sub>5</sub> vs. t<sub>6</sub>: p = 0.006, t<sub>5</sub> vs. t<sub>7</sub>: p = 0.022 |
| stria terminalis | 119 | 301 | 0.59 ± 0.04 | 0.61 ± 0.06 | 0.61 ± 0.05 | 0.61 ± 0.03 | 0.64 ± 0.07 | 0.79 ± 0.06 | 0.82 ± 0.03 | < 0.001 | t<sub>1</sub> vs. t<sub>6</sub>: p < 0.001, t<sub>1</sub> vs. t<sub>7</sub>: p < 0.001, t<sub>2</sub> vs. t<sub>6</sub>: p = 0.003, t<sub>2</sub> vs. t<sub>7</sub>: p = 0.010, t<sub>3</sub> vs. t<sub>6</sub>: p = 0.004, t<sub>3</sub> vs. t<sub>7</sub>: p = 0.013, t<sub>4</sub> vs. t<sub>7</sub>: p = 0.011, t<sub>5</sub> vs. t<sub>7</sub>: p = 0.028 |
| supraoptic commissures | 120 | 349 | 0.85 ± 0.29 | 1.05 ± 0.22 | 0.69 ± 0.17 | 0.68 ± 0.17 | 0.93 ± 0.27 | 1.30 ± 0.34 | 1.30 ± 0.25 | < 0.001 | t<sub>1</sub> vs. t<sub>7</sub>: p = 0.033, t<sub>3</sub> vs. t<sub>6</sub>: p = 0.021, t<sub>3</sub> vs. t<sub>7</sub>: p = 0.002, t<sub>4</sub> vs. t<sub>6</sub>: p = 0.006, t<sub>4</sub> vs. t<sub>7</sub>: p = 0.001, t<sub>5</sub> vs. t<sub>7</sub>: p = 0.004 |
| mammillary related | 121 | 46 | 0.99 ± 0.15 | 0.85 ± 0.14 | 0.85 ± 0.20 | 0.83 ± 0.13 | 0.90 ± 0.18 | 1.10 ± 0.14 | 1.09 ± 0.19 | = 0.010 | t<sub>4</sub> vs. t<sub>6</sub>: p = 0.028 |
| epithalamus related | 122 | 1083 | 1.11 ± 0.04 | 1.02 ± 0.08 | 1.00 ± 0.06 | 1.00 ± 0.13 | 1.01 ± 0.11 | 1.07 ± 0.08 | 1.01 ± 0.02 | = 0.007 | t<sub>1</sub> vs. t<sub>3</sub>: p = 0.015, t<sub>1</sub> vs. t<sub>4</sub>: p = 0.008, t<sub>1</sub> vs. t<sub>5</sub>: p = 0.021 |

**Supplemental Table ST2.**
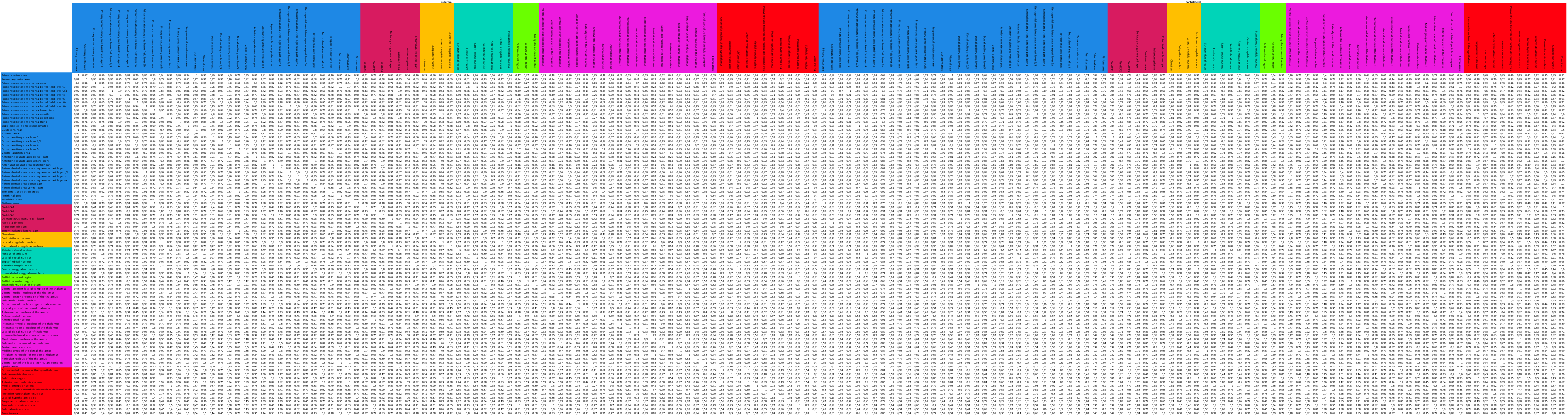
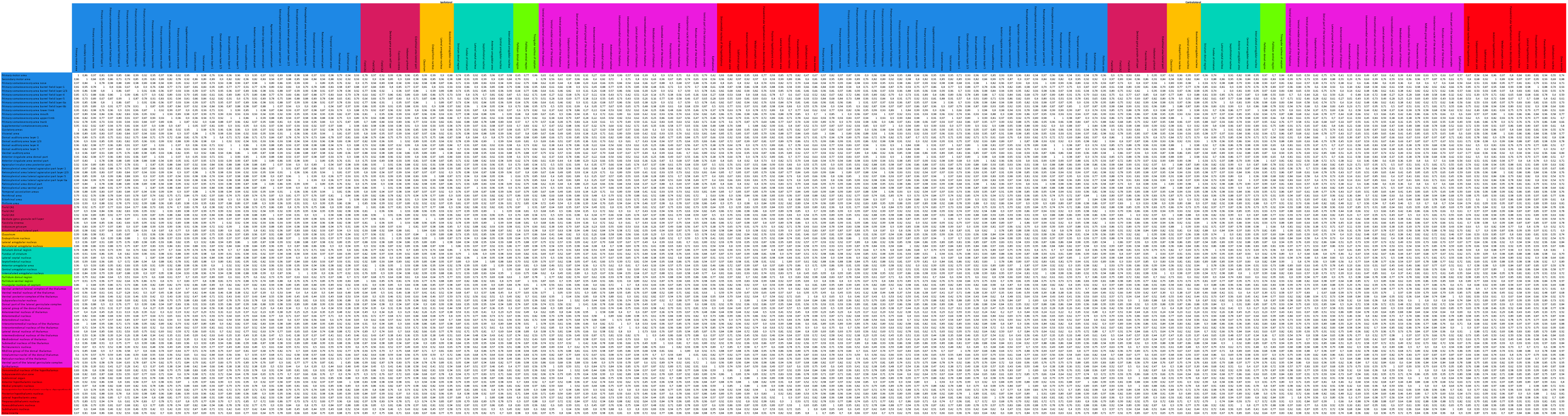
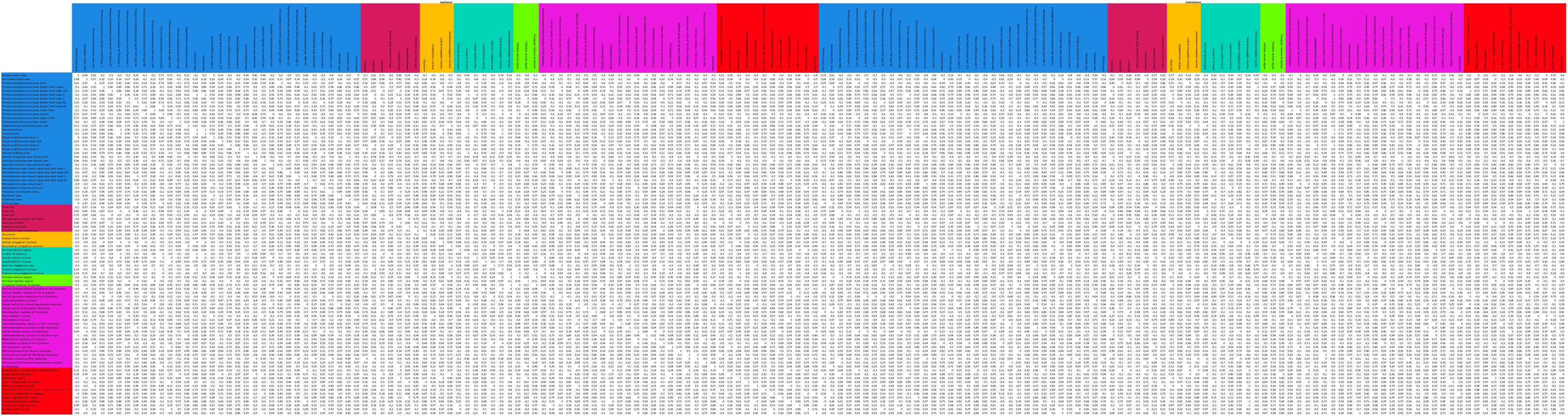
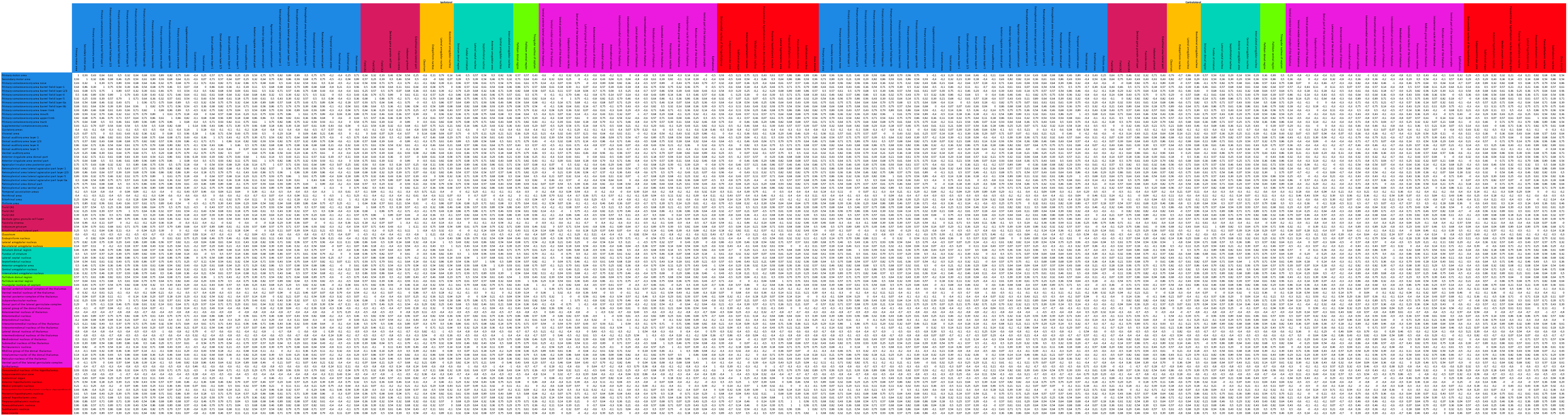
Biomechanical connectivity (BC) for parcellation and age matched to functional connectivity. Static BC i s before exponential transform and rescaling to 0-1. Dynamic BC i s before Fisher z transformation and rescaling. **Sheet**: Dynamic BC stiffness

**Supplemental Table ST3.**
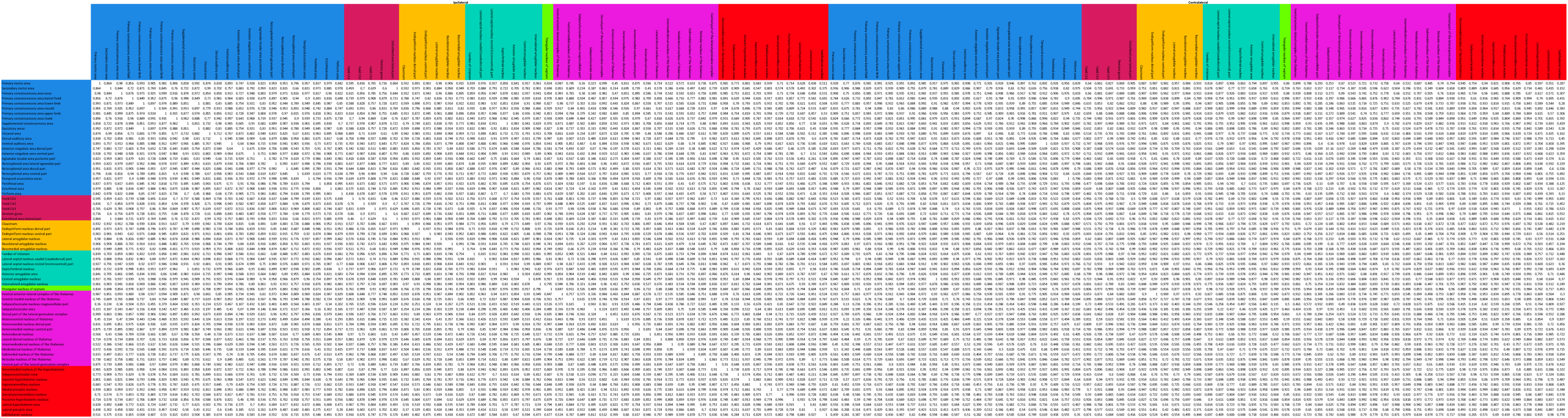
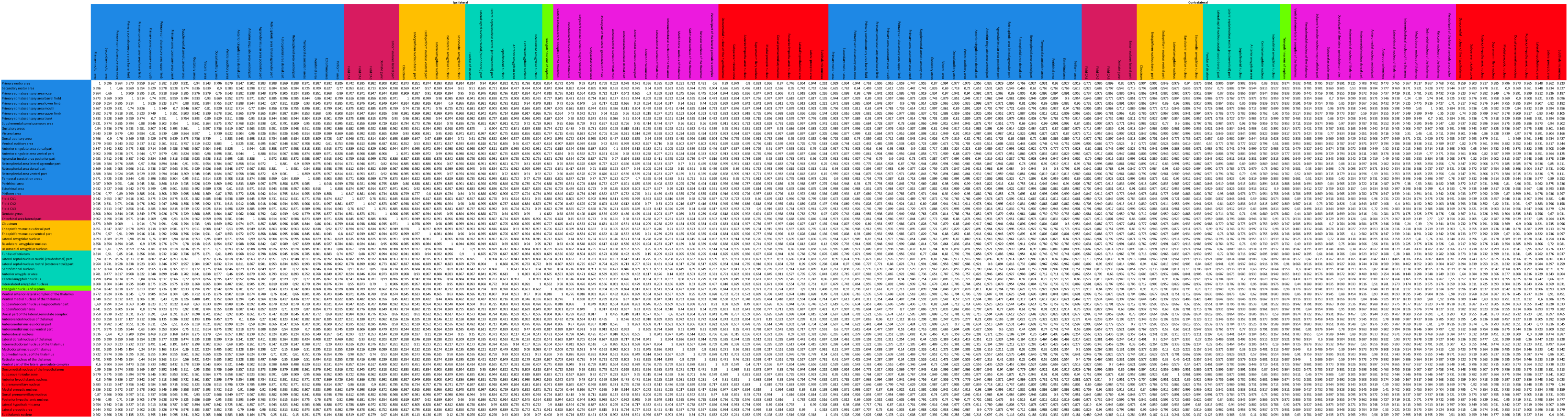
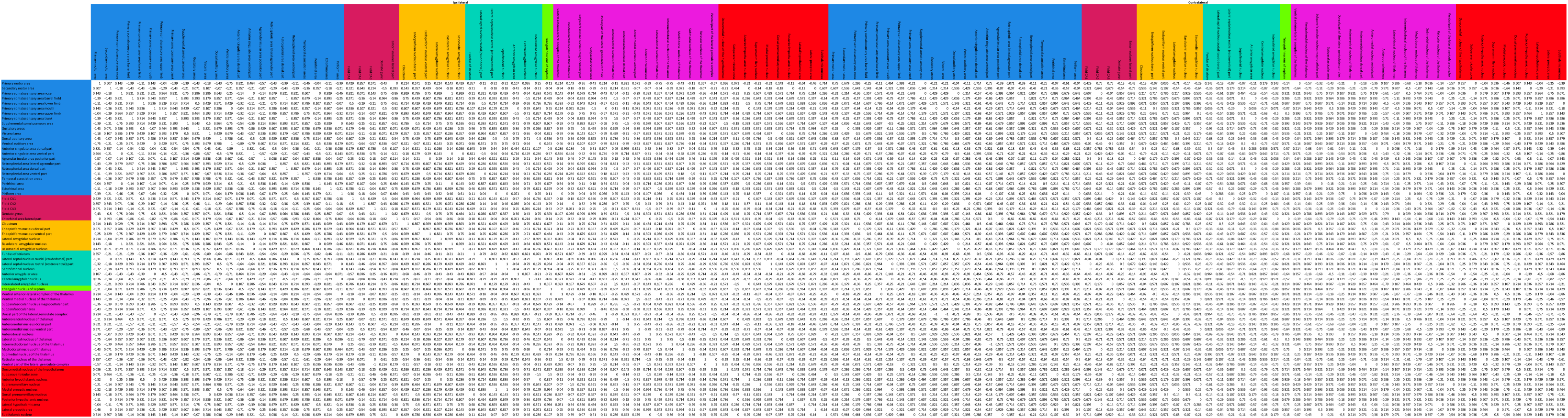
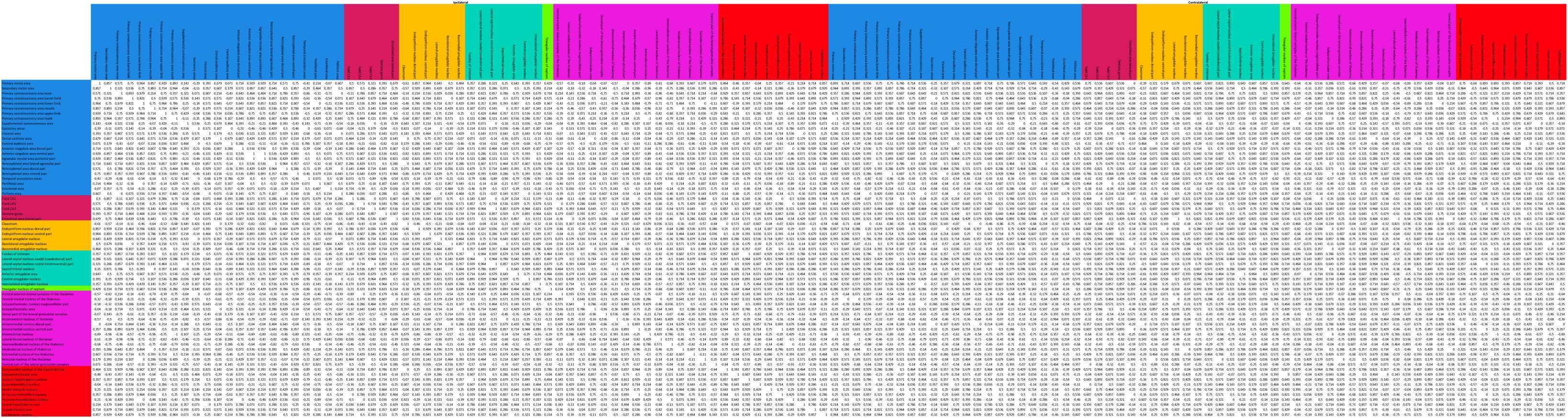
Biomechanical connectivity (BC) for parcellation and age matched to structural connectivity. Static BC is before exponential transform and rescaling to 0-1. Dynamic BC is before Fisher z transformation and rescaling. **Sheet**: Static BC stiffness

**Supplemental Table ST4.** Age-dependent community assignments for all 122 brain regions identified by community detection analysis using absolute and relative changes in cerebral biomechanical properties. Brain regions are color-coded according to their anatomical classifications in the Allen Brain Atlas.

| ROI label | ROI ID | ROI ID Allen | Community EAdol | Community MAdol | Community LAdol | Community EAdult | Community MAdult | Community LAdult | Community Old |
|---|---|---|---|---|---|---|---|---|---|
| Primary motor area | 1 | 985 | 1 | 1 | 1 | 1 | 1 | 3 | 1 |
| Primary somatosensory area nose | 3 | 353 | 1 | 1 | 1 | 4 | 1 | 1 | 1 |
| Primary somatosensory area barrel field layer 4 | 6 | 1047 | 1 | 1 | 1 | 4 | 2 | 1 | 2 |
| Primary somatosensory area barrel field layer 5 | 7 | 1070 | 1 | 1 | 2 | 2 | 2 | 2 | 1 |
| Primary somatosensory area barrel field layer 6a | 8 | 1038 | 1 | 2 | 2 | 2 | 4 | 2 | 3 |
| Primary somatosensory area barrel field layer 6b | 9 | 1062 | 1 | 1 | 1 | 2 | 2 | 1 | 1 |
| Primary somatosensory area lower limb | 10 | 337 | 1 | 1 | 1 | 1 | 2 | 2 | 1 |
| Primary somatosensory area mouth | 11 | 345 | 1 | 1 | 1 | 1 | 2 | 3 | 3 |
| Primary somatosensory area upper limb | 12 | 369 | 1 | 1 | 1 | 1 | 1 | 2 | 1 |
| Primary somatosensory area trunk | 13 | 361 | 1 | 1 | 2 | 1 | 2 | 5 | 2 |
| Supplemental somatosensory area | 14 | 378 | 1 | 1 | 1 | 1 | 2 | 1 | 1 |
| Gustatory areas | 15 | 1057 | 1 | 1 | 1 | 4 | 1 | 1 | 2 |
| Visceral area | 16 | 677 | 1 | 1 | 3 | 5 | 1 | 1 | 2 |
| Dorsal auditory area layer 1 | 17 | 527 | 1 | 1 | 1 | 5 | 1 | 5 | 2 |
| Dorsal auditory area layer 2/3 | 18 | 600 | 1 | 1 | 1 | 5 | 1 | 5 | 1 |
| Dorsal auditory area layer 4 | 19 | 678 | 1 | 1 | 2 | 4 | 2 | 1 | 2 |
| Dorsal auditory area layer 5 | 20 | 252 | 1 | 1 | 4 | 1 | 4 | 2 | 1 |
| Dorsal auditory area layer 6a | 21 | 156 | 1 | 2 | 4 | 2 | 3 | 3 | 3 |
| Dorsal auditory area layer 6b | 22 | 243 | 1 | 1 | 2 | 2 | 4 | 1 | 1 |
| Ventral auditory area | 23 | 1018 | 1 | 1 | 1 | 4 | 1 | 1 | 2 |
| Anterior cingulate area ventral part | 25 | 48 | 1 | 1 | 1 | 2 | 2 | 4 | 4 |
| Agranular insular area posterior part | 26 | 111 | 1 | 1 | 3 | 5 | 5 | 5 | 2 |
| Retrosplenial area lateral agranular part layer 5 | 29 | 774 | 1 | 1 | 2 | 2 | 2 | 2 | 1 |
| Retrosplenial area lateral agranular part layer 6a | 30 | 906 | 1 | 1 | 1 | 1 | 1 | 2 | 1 |
| Retrosplenial area lateral agranular part layer 6b | 31 | 279 | 1 | 1 | 3 | 4 | 5 | 1 | 1 |
| Retrosplenial area dorsal part | 32 | 879 | 1 | 1 | 1 | 1 | 2 | 1 | 2 |
| Retrosplenial area ventral part | 33 | 886 | 1 | 2 | 4 | 3 | 3 | 3 | 3 |
| Temporal association areas | 34 | 541 | 1 | 1 | 1 | 1 | 2 | 1 | 2 |
| Perirhinal area | 35 | 922 | 1 | 1 | 3 | 4 | 1 | 1 | 2 |
| Ectorhinal area | 36 | 895 | 1 | 1 | 1 | 1 | 2 | 1 | 2 |
| Field CA1 | 38 | 382 | 1 | 3 | 5 | 2 | 3 | 4 | 4 |
| Field CA2 | 39 | 423 | 1 | 2 | 2 | 1 | 2 | 4 | 4 |
| Field CA3 | 40 | 463 | 1 | 2 | 2 | 2 | 4 | 6 | 4 |
| Dentate gyrus molecular layer | 41 | 10703 | 1 | 1 | 2 | 2 | 4 | 3 | 3 |
| Dentate gyrus polymorph layer | 42 | 10704 | 1 | 1 | 2 | 2 | 3 | 3 | 3 |
| Dentate gyrus granule cell layer | 43 | 632 | 1 | 1 | 1 | 1 | 4 | 2 | 1 |
| Induseum griseum | 45 | 19 | 1 | 3 | 2 | 1 | 4 | 3 | 4 |
| Entorhinal area lateral part | 46 | 918 | 1 | 1 | 3 | 4 | 5 | 1 | 1 |
| Prosubiculum ventral part | 47 | 484682492 | 1 | 1 | 1 | 1 | 1 | 2 | 1 |
| Claustrum | 48 | 583 | 1 | 1 | 1 | 4 | 1 | 1 | 2 |
| Basomedial amygdalar nucleus | 52 | 319 | 1 | 1 | 3 | 1 | 1 | 2 | 2 |
| Striatum dorsal region | 54 | 485 | 1 | 2 | 4 | 3 | 3 | 6 | 4 |
| Fundus of striatum | 55 | 998 | 1 | 1 | 2 | 3 | 4 | 4 | 4 |
| Lateral septal nucleus | 56 | 242 | 1 | 1 | 3 | 4 | 5 | 1 | 2 |
| Septofimbrial nucleus | 57 | 310 | 1 | 2 | 2 | 2 | 2 | 1 | 1 |
| Anterior amygdalar area | 58 | 23 | 1 | 1 | 2 | 3 | 4 | 4 | 5 |
| Central amygdalar nucleus | 59 | 536 | 1 | 1 | 1 | 2 | 2 | 3 | 4 |
| Triangular nucleus of septum | 63 | 581 | 1 | 2 | 2 | 2 | 2 | 1 | 1 |
| Ventral anterior-lateral complex of the thalamus | 64 | 629 | 1 | 2 | 1 | 3 | 3 | 2 | 4 |
| Ventral medial nucleus of the thalamus | 65 | 685 | 1 | 1 | 2 | 6 | 3 | 4 | 3 |
| Ventral posterior complex of the thalamus | 66 | 709 | 1 | 2 | 2 | 3 | 3 | 6 | 4 |
| Dorsal part of the lateral geniculate complex | 69 | 170 | 1 | 1 | 1 | 1 | 1 | 3 | 1 |
| Anteromedial nucleus | 72 | 127 | 1 | 2 | 2 | 3 | 3 | 6 | 5 |
| Interanteromedial nucleus of the thalamus | 74 | 1120 | 1 | 2 | 4 | 2 | 3 | 3 | 3 |
| Interanterodorsal nucleus of the thalamus | 75 | 1113 | 1 | 2 | 4 | 3 | 3 | 3 | 3 |
| Submedial nucleus of the thalamus | 79 | 366 | 1 | 2 | 6 | 3 | 3 | 6 | 3 |
| Midline group of the dorsal thalamus | 81 | 571 | 1 | 2 | 1 | 1 | 2 | 2 | 1 |
| Intralaminar nuclei of the dorsal thalamus | 82 | 51 | 1 | 2 | 6 | 3 | 3 | 1 | 1 |
| Ventral part of the lateral geniculate complex | 84 | 178 | 1 | 1 | 2 | 2 | 2 | 3 | 1 |
| Accessory supraoptic group | 86 | 332 | 1 | 1 | 1 | 1 | 1 | 1 | 1 |
| Periventricular hypothalamic nucleus intermediate part | 88 | 118 | 1 | 2 | 1 | 2 | 1 | 1 | 2 |
| Dorsomedial nucleus of the hypothalamus | 89 | 830 | 1 | 2 | 1 | 1 | 2 | 2 | 3 |
| Subfornical organ | 91 | 338 | 1 | 2 | 1 | 1 | 1 | 5 | 2 |
| Supramammillary nucleus | 93 | 525 | 1 | 2 | 2 | 2 | 3 | 3 | 4 |
| Medial preoptic nucleus | 94 | 515 | 1 | 1 | 1 | 1 | 2 | 1 | 2 |
| Dorsal premammillary nucleus | 95 | 980 | 1 | 2 | 4 | 2 | 3 | 4 | 4 |
| Paraventricular hypothalamic nucleus descending division | 96 | 63 | 1 | 2 | 2 | 2 | 3 | 1 | 1 |
| Posterior hypothalamic nucleus | 97 | 946 | 1 | 2 | 2 | 2 | 3 | 3 | 3 |
| optic nerve | 104 | 848 | 1 | 1 | 3 | 1 | 1 | 3 | 1 |
| corpus callosum anterior forceps | 105 | 956 | 1 | 1 | 1 | 1 | 2 | 3 | 1 |
| corpus callosum splenium | 107 | 986 | 1 | 1 | 1 | 1 | 2 | 2 | 1 |
| anterior commissure temporal limb | 112 | 908 | 1 | 1 | 2 | 2 | 2 | 4 | 3 |
| cingulum bundle | 113 | 940 | 1 | 1 | 3 | 4 | 5 | 1 | 1 |
| alveus | 114 | 466 | 1 | 1 | 1 | 4 | 1 | 3 | 1 |
| dorsal fornix | 115 | 530 | 1 | 2 | 2 | 4 | 2 | 2 | 1 |
| fimbria | 116 | 603 | 1 | 1 | 1 | 1 | 1 | 3 | 1 |
| hippocampal commissures | 118 | 618 | 1 | 1 | 1 | 4 | 1 | 2 | 1 |
| stria terminalis | 119 | 301 | 1 | 1 | 1 | 1 | 1 | 2 | 1 |
| Pallidum dorsal region | 61 | 818 | 2 | 2 | 6 | 6 | 3 | 4 | 4 |
| Anteroventral nucleus of thalamus | 71 | 255 | 2 | 3 | 6 | 6 | 6 | 4 | 5 |
| Anterodorsal nucleus | 73 | 64 | 2 | 2 | 4 | 3 | 3 | 6 | 3 |
| Lateral dorsal nucleus of thalamus | 76 | 155 | 2 | 2 | 4 | 3 | 4 | 4 | 3 |
| Intermediodorsal nucleus of the thalamus | 77 | 59 | 2 | 2 | 5 | 3 | 3 | 1 | 2 |
| Mediodorsal nucleus of thalamus | 78 | 362 | 2 | 3 | 5 | 3 | 3 | 4 | 4 |
| Perireunensis nucleus | 80 | 1077 | 2 | 2 | 5 | 3 | 6 | 4 | 4 |
| Reticular nucleus of the thalamus | 83 | 262 | 2 | 2 | 4 | 3 | 3 | 6 | 4 |
| Epithalamus | 85 | 958 | 2 | 2 | 4 | 1 | 4 | 2 | 1 |
| Anterior hypothalamic nucleus | 92 | 88 | 2 | 2 | 2 | 2 | 2 | 2 | 1 |
| Lateral hypothalamic area | 98 | 194 | 2 | 3 | 5 | 6 | 6 | 3 | 1 |
| Lateral preoptic area | 99 | 226 | 2 | 3 | 6 | 6 | 6 | 3 | 3 |
| Zona incerta | 103 | 797 | 2 | 2 | 5 | 3 | 3 | 4 | 3 |
| corticospinal tract | 108 | 784 | 2 | 2 | 4 | 3 | 3 | 4 | 4 |
| thalamus related | 109 | 896 | 2 | 2 | 2 | 3 | 3 | 6 | 4 |
| postcommissural fornix | 117 | 737 | 2 | 2 | 2 | 3 | 3 | 2 | 3 |
| supraoptic commissures | 120 | 349 | 2 | 2 | 2 | 3 | 3 | 6 | 3 |
| epithalamus related | 122 | 1083 | 2 | 2 | 2 | 2 | 4 | 3 | 3 |
| Secondary motor area | 2 | 993 | 3 | 1 | 3 | 5 | 5 | 1 | 2 |
| Primary somatosensory area barrel field layer 1 | 4 | 981 | 3 | 1 | 3 | 5 | 5 | 5 | 2 |
| Primary somatosensory area barrel field layer 2/3 | 5 | 201 | 3 | 1 | 3 | 5 | 1 | 5 | 2 |
| Anterior cingulate area dorsal part | 24 | 39 | 3 | 1 | 3 | 5 | 5 | 1 | 1 |
| Retrosplenial area lateral agranular part layer 1 | 27 | 671 | 3 | 1 | 3 | 4 | 5 | 5 | 2 |
| Retrosplenial area lateral agranular part layer 2/3 | 28 | 965 | 3 | 1 | 1 | 4 | 2 | 5 | 2 |
| Piriform area | 37 | 961 | 3 | 1 | 3 | 5 | 1 | 5 | 2 |
| Endopiriform nucleus | 49 | 942 | 3 | 1 | 1 | 5 | 1 | 1 | 2 |
| Lateral amygdalar nucleus | 50 | 131 | 3 | 1 | 1 | 1 | 1 | 1 | 2 |
| Basolateral amygdalar nucleus | 51 | 295 | 3 | 1 | 3 | 4 | 2 | 1 | 2 |
| Intercalated amygdalar nucleus | 60 | 1105 | 3 | 1 | 1 | 4 | 1 | 2 | 1 |
| corpus callosum extreme capsule | 106 | 964 | 3 | 1 | 3 | 5 | 1 | 1 | 2 |
| amygdalar capsule | 111 | 884 | 3 | 1 | 1 | 4 | 1 | 1 | 2 |
| Lateral group of the dorsal thalamus | 70 | 138 | 4 | 4 | 6 | 6 | 6 | 6 | 5 |
| ROI label | ROI ID | ROI ID Allen | Community EAdol | Community MAdol | Community LAdol | Community EAdult | Community MAdult | Community LAdult | Community Old |
| Primary motor area | 1 | 985 | 1 | 1 | 1 | 1 | 1 | 1 | 1 |
| Secondary motor area | 2 | 993 | 1 | 1 | 6 | 6 | 1 | 1 | 1 |
| Primary somatosensory area nose | 3 | 353 | 1 | 1 | 1 | 1 | 1 | 1 | 1 |
| Primary somatosensory area barrel field layer 1 | 4 | 981 | 1 | 1 | 2 | 6 | 1 | 1 | 1 |
| Primary somatosensory area barrel field layer 2/3 | 5 | 201 | 1 | 1 | 1 | 1 | 1 | 1 | 1 |
| Primary somatosensory area barrel field layer 4 | 6 | 1047 | 1 | 1 | 3 | 3 | 4 | 1 | 1 |
| Primary somatosensory area barrel field layer 5 | 7 | 1070 | 1 | 1 | 3 | 3 | 1 | 1 | 1 |
| Primary somatosensory area barrel field layer 6a | 8 | 1038 | 1 | 1 | 1 | 1 | 1 | 1 | 1 |
| Primary somatosensory area barrel field layer 6b | 9 | 1062 | 1 | 1 | 1 | 1 | 1 | 1 | 1 |
| Primary somatosensory area lower limb | 10 | 337 | 1 | 1 | 1 | 3 | 1 | 1 | 1 |
| Primary somatosensory area mouth | 11 | 345 | 1 | 1 | 2 | 1 | 1 | 1 | 1 |
| Primary somatosensory area upper limb | 12 | 369 | 1 | 1 | 1 | 3 | 1 | 1 | 1 |
| Primary somatosensory area trunk | 13 | 361 | 1 | 1 | 3 | 3 | 1 | 1 | 1 |
| Supplemental somatosensory area | 14 | 378 | 1 | 1 | 2 | 1 | 1 | 1 | 1 |
| Gustatory areas | 15 | 1057 | 1 | 1 | 2 | 1 | 1 | 3 | 3 |
| Visceral area | 16 | 677 | 1 | 1 | 1 | 1 | 1 | 1 | 1 |
| Dorsal auditory area layer 1 | 17 | 527 | 1 | 2 | 4 | 2 | 2 | 1 | 1 |
| Dorsal auditory area layer 2/3 | 18 | 600 | 1 | 1 | 2 | 1 | 1 | 1 | 1 |
| Dorsal auditory area layer 4 | 19 | 678 | 1 | 1 | 2 | 1 | 1 | 1 | 1 |
| Dorsal auditory area layer 5 | 20 | 252 | 1 | 1 | 1 | 1 | 1 | 3 | 3 |
| Dorsal auditory area layer 6a | 21 | 156 | 1 | 1 | 2 | 1 | 1 | 3 | 1 |
| Dorsal auditory area layer 6b | 22 | 243 | 1 | 1 | 4 | 2 | 1 | 1 | 1 |
| Ventral auditory area | 23 | 1018 | 1 | 1 | 6 | 1 | 1 | 1 | 1 |
| Anterior cingulate area dorsal part | 24 | 39 | 1 | 1 | 1 | 1 | 1 | 1 | 1 |
| Anterior cingulate area ventral part | 25 | 48 | 1 | 1 | 1 | 3 | 1 | 1 | 1 |
| Agranular insular area posterior part | 26 | 111 | 1 | 1 | 1 | 1 | 1 | 1 | 1 |
| Retrosplenial area lateral agranular part layer 1 | 27 | 671 | 1 | 1 | 2 | 1 | 1 | 1 | 1 |
| Retrosplenial area lateral agranular part layer 2/3 | 28 | 965 | 1 | 1 | 1 | 1 | 1 | 1 | 1 |
| Retrosplenial area lateral agranular part layer 5 | 29 | 774 | 1 | 1 | 1 | 1 | 1 | 1 | 1 |
| Retrosplenial area lateral agranular part layer 6a | 30 | 906 | 1 | 1 | 1 | 1 | 1 | 1 | 1 |
| Retrosplenial area lateral agranular part layer 6b | 31 | 279 | 1 | 1 | 3 | 1 | 1 | 1 | 1 |
| Retrosplenial area dorsal part | 32 | 879 | 1 | 1 | 3 | 3 | 1 | 1 | 1 |
| Retrosplenial area ventral part | 33 | 886 | 1 | 1 | 1 | 1 | 1 | 1 | 1 |
| Temporal association areas | 34 | 541 | 1 | 1 | 2 | 1 | 1 | 3 | 3 |
| Perirhinal area | 35 | 922 | 1 | 1 | 2 | 2 | 1 | 1 | 1 |
| Ectorhinal area | 36 | 895 | 1 | 1 | 2 | 1 | 1 | 3 | 3 |
| Piriform area | 37 | 961 | 1 | 1 | 2 | 1 | 1 | 1 | 1 |
| Field CA1 | 38 | 382 | 1 | 1 | 4 | 2 | 1 | 1 | 1 |
| Field CA2 | 39 | 423 | 1 | 1 | 1 | 1 | 1 | 1 | 1 |
| Field CA3 | 40 | 463 | 1 | 1 | 2 | 2 | 1 | 1 | 1 |
| Dentate gyrus molecular layer | 41 | 10703 | 1 | 1 | 1 | 1 | 1 | 1 | 1 |
| Dentate gyrus polymorph layer | 42 | 10704 | 1 | 1 | 1 | 1 | 1 | 1 | 1 |
| Dentate gyrus granule cell layer | 43 | 632 | 1 | 1 | 1 | 1 | 1 | 1 | 1 |
| Fasciola cinerea | 44 | 982 | 1 | 1 | 2 | 1 | 1 | 1 | 3 |
| Induseum griseum | 45 | 19 | 1 | 1 | 3 | 3 | 1 | 1 | 2 |
| Entorhinal area lateral part | 46 | 918 | 1 | 1 | 1 | 1 | 1 | 1 | 1 |
| Prosubiculum ventral part | 47 | 484682492 | 1 | 1 | 1 | 1 | 1 | 1 | 1 |
| Endopiriform nucleus | 49 | 942 | 1 | 1 | 1 | 3 | 1 | 1 | 3 |
| Lateral amygdalar nucleus | 50 | 131 | 1 | 1 | 1 | 1 | 1 | 1 | 1 |
| Basolateral amygdalar nucleus | 51 | 295 | 1 | 1 | 1 | 1 | 1 | 1 | 3 |
| Basomedial amygdalar nucleus | 52 | 319 | 1 | 1 | 1 | 1 | 1 | 1 | 1 |
| Posterior amygdalar nucleus | 53 | 780 | 1 | 1 | 1 | 1 | 1 | 1 | 1 |
| Fundus of striatum | 55 | 998 | 1 | 1 | 3 | 1 | 1 | 1 | 1 |
| Lateral septal nucleus | 56 | 242 | 1 | 1 | 2 | 1 | 1 | 1 | 1 |
| Septofimbrial nucleus | 57 | 310 | 1 | 1 | 6 | 2 | 1 | 1 | 1 |
| Anterior amygdalar area | 58 | 23 | 1 | 1 | 3 | 1 | 1 | 1 | 1 |
| Central amygdalar nucleus | 59 | 536 | 1 | 1 | 1 | 1 | 1 | 1 | 1 |
| Intercalated amygdalar nucleus | 60 | 1105 | 1 | 1 | 3 | 1 | 1 | 1 | 1 |
| Pallidum ventral region | 62 | 835 | 1 | 1 | 2 | 1 | 1 | 1 | 1 |
| Triangular nucleus of septum | 63 | 581 | 1 | 1 | 6 | 2 | 1 | 1 | 1 |
| Dorsal part of the lateral geniculate complex | 69 | 170 | 1 | 1 | 6 | 2 | 1 | 1 | 1 |
| Submedial nucleus of the thalamus | 79 | 366 | 1 | 1 | 6 | 1 | 1 | 2 | 1 |
| Intralaminar nuclei of the dorsal thalamus | 82 | 51 | 1 | 2 | 1 | 1 | 1 | 2 | 1 |
| Accessory supraoptic group | 86 | 332 | 1 | 1 | 2 | 1 | 1 | 1 | 1 |
| Periventricular hypothalamic nucleus anterior part | 87 | 30 | 1 | 2 | 2 | 2 | 2 | 1 | 1 |
| Dorsomedial nucleus of the hypothalamus | 89 | 830 | 1 | 1 | 2 | 1 | 1 | 1 | 1 |
| Subparaventricular zone | 90 | 347 | 1 | 2 | 2 | 2 | 1 | 1 | 1 |
| Subfornical organ | 91 | 338 | 1 | 1 | 2 | 2 | 1 | 1 | 1 |
| Anterior hypothalamic nucleus | 92 | 88 | 1 | 1 | 3 | 1 | 1 | 1 | 1 |
| Supramammillary nucleus | 93 | 525 | 1 | 1 | 2 | 1 | 1 | 1 | 1 |
| Medial preoptic nucleus | 94 | 515 | 1 | 1 | 2 | 1 | 1 | 1 | 1 |
| Dorsal premammillary nucleus | 95 | 980 | 1 | 1 | 1 | 1 | 1 | 1 | 1 |
| Paraventricular hypothalamic nucleus descending division | 96 | 63 | 1 | 1 | 6 | 2 | 1 | 1 | 1 |
| Posterior hypothalamic nucleus | 97 | 946 | 1 | 1 | 6 | 2 | 1 | 1 | 1 |
| Lateral hypothalamic area | 98 | 194 | 1 | 1 | 2 | 1 | 1 | 1 | 1 |
| Lateral preoptic area | 99 | 226 | 1 | 1 | 2 | 1 | 1 | 1 | 1 |
| Preparasubthalamic nucleus | 100 | 356 | 1 | 1 | 4 | 1 | 1 | 2 | 2 |
| Parasubthalamic nucleus | 101 | 364 | 1 | 1 | 6 | 1 | 1 | 2 | 2 |
| optic nerve | 104 | 848 | 1 | 2 | 2 | 1 | 1 | 2 | 1 |
| corpus callosum anterior forceps | 105 | 956 | 1 | 1 | 2 | 1 | 1 | 1 | 3 |
| corpus callosum extreme capsule | 106 | 964 | 1 | 1 | 1 | 1 | 1 | 1 | 3 |
| corpus callosum splenium | 107 | 986 | 1 | 1 | 1 | 1 | 1 | 1 | 1 |
| amygdalar capsule | 111 | 884 | 1 | 1 | 1 | 1 | 1 | 1 | 3 |
| anterior commissure temporal limb | 112 | 908 | 1 | 1 | 3 | 1 | 1 | 1 | 1 |
| cingulum bundle | 113 | 940 | 1 | 1 | 3 | 3 | 1 | 1 | 1 |
| alveus | 114 | 466 | 1 | 1 | 1 | 1 | 1 | 1 | 1 |
| dorsal fornix | 115 | 530 | 1 | 1 | 3 | 3 | 1 | 1 | 1 |
| postcommissural fornix | 117 | 737 | 1 | 1 | 2 | 1 | 1 | 1 | 1 |
| hippocampal commissures | 118 | 618 | 1 | 1 | 1 | 1 | 1 | 1 | 1 |
| stria terminalis | 119 | 301 | 1 | 1 | 2 | 1 | 1 | 1 | 1 |
| Striatum dorsal region | 54 | 485 | 2 | 2 | 4 | 2 | 2 | 1 | 1 |
| Pallidum dorsal region | 61 | 818 | 2 | 2 | 4 | 2 | 2 | 2 | 2 |
| Subparafascicular nucleus | 67 | 406 | 2 | 2 | 4 | 2 | 1 | 2 | 1 |
| Subparafascicular area | 68 | 609 | 2 | 2 | 6 | 2 | 1 | 1 | 1 |
| Interanteromedial nucleus of the thalamus | 74 | 1120 | 2 | 2 | 1 | 1 | 1 | 1 | 1 |
| Lateral dorsal nucleus of thalamus | 76 | 155 | 2 | 2 | 5 | 4 | 3 | 1 | 1 |
| Midline group of the dorsal thalamus | 81 | 571 | 2 | 2 | 2 | 2 | 2 | 1 | 1 |
| Ventral part of the lateral geniculate complex | 84 | 178 | 2 | 2 | 4 | 2 | 2 | 1 | 1 |
| Epithalamus | 85 | 958 | 2 | 2 | 5 | 4 | 2 | 2 | 1 |
| Periventricular hypothalamic nucleus intermediate part | 88 | 118 | 2 | 2 | 5 | 4 | 3 | 1 | 1 |
| Zona incerta | 103 | 797 | 2 | 2 | 4 | 2 | 2 | 2 | 2 |
| fimbria | 116 | 603 | 2 | 2 | 4 | 2 | 2 | 1 | 1 |
| mammillary related | 121 | 46 | 2 | 2 | 4 | 2 | 2 | 2 | 2 |
| Ventral anterior-lateral complex of the thalamus | 64 | 629 | 3 | 2 | 4 | 4 | 2 | 1 | 1 |
| Ventral medial nucleus of the thalamus | 65 | 685 | 3 | 2 | 5 | 4 | 3 | 2 | 2 |
| Ventral posterior complex of the thalamus | 66 | 709 | 3 | 3 | 5 | 4 | 3 | 2 | 2 |
| Anterodorsal nucleus | 73 | 64 | 3 | 2 | 5 | 5 | 3 | 2 | 2 |
| Intermediodorsal nucleus of the thalamus | 77 | 59 | 3 | 4 | 5 | 5 | 3 | 2 | 1 |
| Mediodorsal nucleus of thalamus | 78 | 362 | 3 | 3 | 5 | 5 | 3 | 4 | 4 |
| Reticular nucleus of the thalamus | 83 | 262 | 3 | 2 | 5 | 4 | 3 | 2 | 2 |
| corticospinal tract | 108 | 784 | 3 | 3 | 5 | 4 | 3 | 2 | 2 |
| epithalamus related | 122 | 1083 | 3 | 2 | 5 | 4 | 2 | 2 | 2 |
| Lateral group of the dorsal thalamus | 70 | 138 | 4 | 4 | 5 | 4 | 3 | 2 | 2 |
| ROI label | ROI ID | ROI ID Allen | Comm. EAdol → MAdol | Comm. MAdol → LAdol | Comm. LAdol → EAdult | Comm. EAdult → MAdult | Comm. MAdult → LAdult | Comm. LAdult → Old |  |
| Primary motor area | 1 | 985 | 1 | 1 | 1 | 2 | 3 | 1 |  |
| Secondary motor area | 2 | 993 | 1 | 1 | 1 | 2 | 3 | 1 |  |
| Primary somatosensory area nose | 3 | 353 | 1 | 1 | 1 | 1 | 1 | 1 |  |
| Primary somatosensory area barrel field layer 1 | 4 | 981 | 1 | 1 | 1 | 1 | 1 | 1 |  |
| Primary somatosensory area barrel field layer 2/3 | 5 | 201 | 1 | 1 | 1 | 4 | 2 | 1 |  |
| Primary somatosensory area barrel field layer 4 | 6 | 1047 | 1 | 1 | 1 | 5 | 2 | 1 |  |
| Primary somatosensory area barrel field layer 5 | 7 | 1070 | 1 | 1 | 1 | 2 | 2 | 1 |  |
| Primary somatosensory area barrel field layer 6a | 8 | 1038 | 1 | 1 | 1 | 3 | 2 | 1 |  |
| Primary somatosensory area barrel field layer 6b | 9 | 1062 | 1 | 1 | 1 | 3 | 2 | 1 |  |
| Primary somatosensory area lower limb | 10 | 337 | 1 | 1 | 1 | 2 | 1 | 1 |  |
| Primary somatosensory area mouth | 11 | 345 | 1 | 1 | 1 | 1 | 1 | 1 |  |
| Primary somatosensory area upper limb | 12 | 369 | 1 | 1 | 1 | 2 | 1 | 1 |  |
| Primary somatosensory area trunk | 13 | 361 | 1 | 2 | 1 | 1 | 5 | 1 |  |
| Supplemental somatosensory area | 14 | 378 | 1 | 1 | 1 | 1 | 2 | 1 |  |
| Gustatory areas | 15 | 1057 | 1 | 1 | 1 | 2 | 1 | 2 |  |
| Visceral area | 16 | 677 | 1 | 1 | 1 | 1 | 1 | 1 |  |
| Dorsal auditory area layer 1 | 17 | 527 | 1 | 1 | 1 | 5 | 2 | 1 |  |
| Dorsal auditory area layer 2/3 | 18 | 600 | 1 | 1 | 1 | 1 | 1 | 1 |  |
| Dorsal auditory area layer 4 | 19 | 678 | 1 | 1 | 1 | 1 | 2 | 1 |  |
| Dorsal auditory area layer 5 | 20 | 252 | 1 | 2 | 1 | 1 | 2 | 1 |  |
| Dorsal auditory area layer 6a | 21 | 156 | 1 | 2 | 1 | 5 | 2 | 1 |  |
| Dorsal auditory area layer 6b | 22 | 243 | 1 | 1 | 1 | 1 | 5 | 1 |  |
| Ventral auditory area | 23 | 1018 | 1 | 1 | 1 | 1 | 2 | 1 |  |
| Anterior cingulate area dorsal part | 24 | 39 | 1 | 1 | 1 | 2 | 3 | 1 |  |
| Anterior cingulate area ventral part | 25 | 48 | 1 | 1 | 1 | 3 | 3 | 1 |  |
| Retrosplenial area lateral agranular part layer 1 | 27 | 671 | 1 | 1 | 1 | 2 | 2 | 1 |  |
| Retrosplenial area lateral agranular part layer 2/3 | 28 | 965 | 1 | 1 | 1 | 1 | 2 | 1 |  |
| Retrosplenial area lateral agranular part layer 5 | 29 | 774 | 1 | 1 | 1 | 2 | 2 | 1 |  |
| Retrosplenial area dorsal part | 32 | 879 | 1 | 2 | 1 | 1 | 5 | 1 |  |
| Retrosplenial area ventral part | 33 | 886 | 1 | 1 | 1 | 3 | 2 | 1 |  |
| Temporal association areas | 34 | 541 | 1 | 1 | 1 | 4 | 5 | 1 |  |
| Perirhinal area | 35 | 922 | 1 | 1 | 1 | 2 | 1 | 1 |  |
| Ectorhinal area | 36 | 895 | 1 | 1 | 1 | 1 | 2 | 1 |  |
| Piriform area | 37 | 961 | 1 | 1 | 1 | 1 | 1 | 1 |  |
| Dentate gyrus molecular layer | 41 | 10703 | 1 | 1 | 1 | 1 | 2 | 1 |  |
| Dentate gyrus polymorph layer | 42 | 10704 | 1 | 1 | 1 | 1 | 2 | 1 |  |
| Dentate gyrus granule cell layer | 43 | 632 | 1 | 1 | 1 | 1 | 2 | 1 |  |
| Fasciola cinerea | 44 | 982 | 1 | 1 | 1 | 1 | 3 | 1 |  |
| Prosubiculum ventral part | 47 | 484682492 | 1 | 1 | 1 | 2 | 1 | 1 |  |
| Claustrum | 48 | 583 | 1 | 1 | 1 | 1 | 1 | 2 |  |
| Lateral amygdalar nucleus | 50 | 131 | 1 | 2 | 1 | 2 | 2 | 1 |  |
| Basomedial amygdalar nucleus | 52 | 319 | 1 | 1 | 1 | 2 | 1 | 2 |  |
| Posterior amygdalar nucleus | 53 | 780 | 1 | 1 | 1 | 2 | 3 | 2 |  |
| Striatum dorsal region | 54 | 485 | 1 | 1 | 1 | 2 | 1 | 1 |  |
| Septofimbrial nucleus | 57 | 310 | 1 | 1 | 1 | 2 | 2 | 1 |  |
| Intercalated amygdalar nucleus | 60 | 1105 | 1 | 1 | 1 | 2 | 1 | 1 |  |
| Pallidum dorsal region | 61 | 818 | 1 | 2 | 1 | 2 | 2 | 1 |  |
| Pallidum ventral region | 62 | 835 | 1 | 1 | 1 | 2 | 5 | 1 |  |
| Triangular nucleus of septum | 63 | 581 | 1 | 1 | 1 | 3 | 2 | 1 |  |
| Ventral anterior-lateral complex of the thalamus | 64 | 629 | 1 | 1 | 3 | 1 | 4 | 4 |  |
| Ventral posterior complex of the thalamus | 66 | 709 | 1 | 1 | 1 | 1 | 1 | 1 |  |
| Dorsal part of the lateral geniculate complex | 69 | 170 | 1 | 2 | 1 | 3 | 3 | 1 |  |
| Lateral group of the dorsal thalamus | 70 | 138 | 1 | 3 | 1 | 4 | 5 | 3 |  |
| Anteromedial nucleus | 72 | 127 | 1 | 1 | 1 | 1 | 1 | 3 |  |
| Interanteromedial nucleus of the thalamus | 74 | 1120 | 1 | 2 | 1 | 1 | 2 | 1 |  |
| Lateral dorsal nucleus of thalamus | 76 | 155 | 1 | 1 | 1 | 3 | 1 | 1 |  |
| Mediodorsal nucleus of thalamus | 78 | 362 | 1 | 1 | 1 | 4 | 2 | 1 |  |
| Perireunensis nucleus | 80 | 1077 | 1 | 1 | 1 | 4 | 4 | 1 |  |
| Midline group of the dorsal thalamus | 81 | 571 | 1 | 1 | 1 | 5 | 2 | 1 |  |
| Reticular nucleus of the thalamus | 83 | 262 | 1 | 1 | 1 | 3 | 1 | 1 |  |
| Epithalamus | 85 | 958 | 1 | 1 | 1 | 4 | 2 | 1 |  |
| Accessory supraoptic group | 86 | 332 | 1 | 1 | 1 | 3 | 1 | 1 |  |
| Periventricular hypothalamic nucleus intermediate part | 88 | 118 | 1 | 3 | 1 | 6 | 1 | 1 |  |
| Subfornical organ | 91 | 338 | 1 | 1 | 1 | 3 | 5 | 1 |  |
| Paraventricular hypothalamic nucleus descending division | 96 | 63 | 1 | 1 | 1 | 4 | 4 | 1 |  |
| Lateral preoptic area | 99 | 226 | 1 | 1 | 1 | 1 | 4 | 1 |  |
| Zona incerta | 103 | 797 | 1 | 1 | 1 | 1 | 2 | 1 |  |
| corpus callosum anterior forceps | 105 | 956 | 1 | 1 | 1 | 1 | 1 | 1 |  |
| corpus callosum splenium | 107 | 986 | 1 | 1 | 1 | 2 | 1 | 1 |  |
| corticospinal tract | 108 | 784 | 1 | 1 | 1 | 2 | 1 | 1 |  |
| thalamus related | 109 | 896 | 1 | 1 | 1 | 2 | 1 | 1 |  |
| cerebral nuclei related | 110 | 760 | 1 | 1 | 1 | 1 | 1 | 1 |  |
| anterior commissure temporal limb | 112 | 908 | 1 | 2 | 1 | 3 | 3 | 1 |  |
| cingulum bundle | 113 | 940 | 1 | 1 | 1 | 2 | 3 | 1 |  |
| alveus | 114 | 466 | 1 | 1 | 1 | 1 | 3 | 1 |  |
| fimbria | 116 | 603 | 1 | 1 | 1 | 2 | 3 | 1 |  |
| postcommissural fornix | 117 | 737 | 1 | 1 | 1 | 4 | 4 | 1 |  |
| hippocampal commissures | 118 | 618 | 1 | 1 | 1 | 2 | 1 | 1 |  |
| stria terminalis | 119 | 301 | 1 | 1 | 1 | 2 | 1 | 1 |  |
| mammillary related | 121 | 46 | 1 | 1 | 1 | 5 | 2 | 1 |  |
| epithalamus related | 122 | 1083 | 1 | 1 | 1 | 1 | 1 | 1 |  |
| Field CA2 | 39 | 423 | 2 | 1 | 1 | 1 | 3 | 1 |  |
| Field CA3 | 40 | 463 | 2 | 1 | 1 | 1 | 3 | 1 |  |
| Endopiriform nucleus | 49 | 942 | 2 | 1 | 1 | 1 | 1 | 2 |  |
| Basolateral amygdalar nucleus | 51 | 295 | 2 | 1 | 1 | 4 | 2 | 1 |  |
| Anteroventral nucleus of thalamus | 71 | 255 | 2 | 1 | 1 | 4 | 4 | 3 |  |
| Interanterodorsal nucleus of the thalamus | 75 | 1113 | 2 | 1 | 1 | 1 | 5 | 1 |  |
| Intralaminar nuclei of the dorsal thalamus | 82 | 51 | 2 | 4 | 4 | 2 | 4 | 1 |  |
| Dorsomedial nucleus of the hypothalamus | 89 | 830 | 2 | 3 | 1 | 1 | 2 | 1 |  |
| Supramammillary nucleus | 93 | 525 | 2 | 1 | 1 | 5 | 2 | 1 |  |
| Lateral hypothalamic area | 98 | 194 | 2 | 1 | 1 | 5 | 4 | 1 |  |
| Subthalamic nucleus | 102 | 470 | 2 | 1 | 1 | 5 | 3 | 2 |  |
| corpus callosum extreme capsule | 106 | 964 | 2 | 1 | 1 | 1 | 1 | 2 |  |
| amygdalar capsule | 111 | 884 | 2 | 1 | 1 | 1 | 1 | 2 |  |
| dorsal fornix | 115 | 530 | 2 | 1 | 1 | 5 | 1 | 1 |  |
| Agranular insular area posterior part | 26 | 111 | 3 | 1 | 1 | 2 | 1 | 1 |  |
| Retrosplenial area lateral agranular part layer 6a | 30 | 906 | 3 | 1 | 1 | 2 | 1 | 1 |  |
| Retrosplenial area lateral agranular part layer 6b | 31 | 279 | 3 | 1 | 1 | 3 | 3 | 1 |  |
| Entorhinal area lateral part | 46 | 918 | 3 | 1 | 1 | 2 | 1 | 1 |  |
| Fundus of striatum | 55 | 998 | 3 | 2 | 1 | 3 | 3 | 1 |  |
| Lateral septal nucleus | 56 | 242 | 3 | 1 | 1 | 3 | 3 | 2 |  |
| Anterior amygdalar area | 58 | 23 | 3 | 2 | 2 | 3 | 1 | 4 |  |
| Central amygdalar nucleus | 59 | 536 | 3 | 2 | 1 | 6 | 3 | 3 |  |
| Ventral medial nucleus of the thalamus | 65 | 685 | 3 | 2 | 2 | 3 | 2 | 1 |  |
| Anterodorsal nucleus | 73 | 64 | 3 | 1 | 1 | 1 | 1 | 2 |  |
| Ventral part of the lateral geniculate complex | 84 | 178 | 3 | 1 | 1 | 3 | 3 | 1 |  |
| Medial preoptic nucleus | 94 | 515 | 3 | 1 | 1 | 2 | 2 | 1 |  |
| optic nerve | 104 | 848 | 3 | 1 | 2 | 3 | 3 | 2 |  |
| Field CA1 | 38 | 382 | 4 | 1 | 1 | 4 | 2 | 1 |  |
| ROI label | ROI ID | ROI ID Allen | Comm. EAdol → MAdol | Comm. MAdol → LAdol | Comm. LAdol → EAdult | Comm. EAdult → MAdult | Comm. MAdult → LAdult | Comm. LAdult → Old |  |
| Primary motor area | 1 | 985 | 1 | 3 | 1 | 1 | 1 | 3 |  |
| Secondary motor area | 2 | 993 | 1 | 6 | 1 | 2 | 2 | 1 |  |
| Primary somatosensory area nose | 3 | 353 | 1 | 2 | 2 | 3 | 1 | 1 |  |
| Primary somatosensory area barrel field layer 1 | 4 | 981 | 1 | 2 | 2 | 2 | 2 | 1 |  |
| Primary somatosensory area barrel field layer 2/3 | 5 | 201 | 1 | 1 | 1 | 1 | 4 | 1 |  |
| Primary somatosensory area barrel field layer 4 | 6 | 1047 | 1 | 1 | 1 | 1 | 4 | 1 |  |
| Primary somatosensory area barrel field layer 5 | 7 | 1070 | 1 | 1 | 1 | 3 | 1 | 2 |  |
| Primary somatosensory area barrel field layer 6a | 8 | 1038 | 1 | 1 | 2 | 1 | 1 | 1 |  |
| Primary somatosensory area lower limb | 10 | 337 | 1 | 3 | 1 | 1 | 4 | 3 |  |
| Primary somatosensory area mouth | 11 | 345 | 1 | 3 | 1 | 1 | 2 | 1 |  |
| Primary somatosensory area upper limb | 12 | 369 | 1 | 1 | 1 | 1 | 1 | 1 |  |
| Primary somatosensory area trunk | 13 | 361 | 1 | 1 | 1 | 1 | 4 | 1 |  |
| Supplemental somatosensory area | 14 | 378 | 1 | 1 | 1 | 1 | 2 | 1 |  |
| Gustatory areas | 15 | 1057 | 1 | 1 | 4 | 1 | 2 | 2 |  |
| Visceral area | 16 | 677 | 1 | 1 | 1 | 1 | 2 | 1 |  |
| Dorsal auditory area layer 1 | 17 | 527 | 1 | 1 | 4 | 4 | 2 | 2 |  |
| Dorsal auditory area layer 2/3 | 18 | 600 | 1 | 1 | 4 | 1 | 1 | 1 |  |
| Dorsal auditory area layer 4 | 19 | 678 | 1 | 1 | 1 | 3 | 1 | 1 |  |
| Dorsal auditory area layer 6a | 21 | 156 | 1 | 3 | 1 | 3 | 3 | 3 |  |
| Anterior cingulate area ventral part | 25 | 48 | 1 | 1 | 1 | 1 | 4 | 1 |  |
| Agranular insular area posterior part | 26 | 111 | 1 | 3 | 2 | 3 | 2 | 1 |  |
| Retrosplenial area lateral agranular part layer 2/3 | 28 | 965 | 1 | 1 | 1 | 2 | 4 | 2 |  |
| Retrosplenial area lateral agranular part layer 5 | 29 | 774 | 1 | 3 | 2 | 3 | 4 | 1 |  |
| Retrosplenial area dorsal part | 32 | 879 | 1 | 1 | 1 | 2 | 4 | 1 |  |
| Retrosplenial area ventral part | 33 | 886 | 1 | 1 | 2 | 2 | 2 | 1 |  |
| Temporal association areas | 34 | 541 | 1 | 1 | 1 | 1 | 3 | 1 |  |
| Ectorhinal area | 36 | 895 | 1 | 3 | 2 | 3 | 3 | 1 |  |
| Field CA2 | 39 | 423 | 1 | 2 | 2 | 1 | 4 | 2 |  |
| Field CA3 | 40 | 463 | 1 | 2 | 3 | 3 | 1 | 1 |  |
| Induseum griseum | 45 | 19 | 1 | 2 | 1 | 2 | 4 | 3 |  |
| Entorhinal area lateral part | 46 | 918 | 1 | 1 | 2 | 2 | 3 | 1 |  |
| Prosubiculum ventral part | 47 | 484682492 | 1 | 1 | 2 | 1 | 1 | 1 |  |
| Claustrum | 48 | 583 | 1 | 1 | 4 | 1 | 3 | 2 |  |
| Endopiriform nucleus | 49 | 942 | 1 | 1 | 1 | 1 | 2 | 2 |  |
| Lateral amygdalar nucleus | 50 | 131 | 1 | 1 | 2 | 1 | 1 | 1 |  |
| Basomedial amygdalar nucleus | 52 | 319 | 1 | 2 | 3 | 3 | 2 | 2 |  |
| Striatum dorsal region | 54 | 485 | 1 | 1 | 1 | 2 | 2 | 1 |  |
| Fundus of striatum | 55 | 998 | 1 | 2 | 3 | 3 | 1 | 1 |  |
| Lateral septal nucleus | 56 | 242 | 1 | 4 | 1 | 2 | 1 | 1 |  |
| Anterior amygdalar area | 58 | 23 | 1 | 2 | 3 | 3 | 4 | 1 |  |
| Intercalated amygdalar nucleus | 60 | 1105 | 1 | 2 | 2 | 1 | 1 | 1 |  |
| Pallidum dorsal region | 61 | 818 | 1 | 1 | 1 | 4 | 2 | 1 |  |
| Ventral posterior complex of the thalamus | 66 | 709 | 1 | 4 | 2 | 5 | 6 | 1 |  |
| Subparafascicular nucleus | 67 | 406 | 1 | 4 | 2 | 3 | 1 | 2 |  |
| Dorsal part of the lateral geniculate complex | 69 | 170 | 1 | 3 | 1 | 1 | 3 | 1 |  |
| Lateral group of the dorsal thalamus | 70 | 138 | 1 | 5 | 2 | 5 | 5 | 1 |  |
| Interanteromedial nucleus of the thalamus | 74 | 1120 | 1 | 5 | 3 | 4 | 1 | 1 |  |
| Mediodorsal nucleus of thalamus | 78 | 362 | 1 | 4 | 5 | 2 | 1 | 2 |  |
| Midline group of the dorsal thalamus | 81 | 571 | 1 | 5 | 3 | 2 | 3 | 1 |  |
| Ventral part of the lateral geniculate complex | 84 | 178 | 1 | 1 | 1 | 4 | 3 | 1 |  |
| Accessory supraoptic group | 86 | 332 | 1 | 1 | 1 | 3 | 2 | 1 |  |
| Periventricular hypothalamic nucleus intermediate part | 88 | 118 | 1 | 3 | 1 | 4 | 5 | 2 |  |
| Supramammillary nucleus | 93 | 525 | 1 | 3 | 1 | 2 | 1 | 1 |  |
| Dorsal premammillary nucleus | 95 | 980 | 1 | 1 | 1 | 1 | 4 | 1 |  |
| Posterior hypothalamic nucleus | 97 | 946 | 1 | 1 | 1 | 1 | 1 | 2 |  |
| Lateral preoptic area | 99 | 226 | 1 | 1 | 4 | 2 | 1 | 1 |  |
| Parasubthalamic nucleus | 101 | 364 | 1 | 1 | 4 | 4 | 4 | 1 |  |
| corpus callosum anterior forceps | 105 | 956 | 1 | 1 | 1 | 1 | 2 | 1 |  |
| corpus callosum extreme capsule | 106 | 964 | 1 | 1 | 1 | 1 | 2 | 2 |  |
| corpus callosum splenium | 107 | 986 | 1 | 1 | 1 | 2 | 2 | 1 |  |
| corticospinal tract | 108 | 784 | 1 | 2 | 1 | 6 | 3 | 1 |  |
| cerebral nuclei related | 110 | 760 | 1 | 3 | 4 | 2 | 2 | 1 |  |
| amygdalar capsule | 111 | 884 | 1 | 2 | 1 | 1 | 2 | 2 |  |
| anterior commissure temporal limb | 112 | 908 | 1 | 4 | 3 | 3 | 1 | 2 |  |
| cingulum bundle | 113 | 940 | 1 | 3 | 1 | 3 | 1 | 1 |  |
| alveus | 114 | 466 | 1 | 1 | 2 | 3 | 1 | 1 |  |
| dorsal fornix | 115 | 530 | 1 | 2 | 1 | 2 | 1 | 1 |  |
| postcommissural fornix | 117 | 737 | 1 | 1 | 4 | 4 | 1 | 1 |  |
| stria terminalis | 119 | 301 | 1 | 1 | 1 | 1 | 1 | 1 |  |
| Retrosplenial area lateral agranular part layer 6a | 30 | 906 | 2 | 3 | 2 | 3 | 1 | 1 |  |
| Retrosplenial area lateral agranular part layer 6b | 31 | 279 | 2 | 3 | 2 | 3 | 1 | 1 |  |
| Perirhinal area | 35 | 922 | 2 | 1 | 3 | 2 | 6 | 1 |  |
| Piriform area | 37 | 961 | 2 | 3 | 2 | 1 | 3 | 3 |  |
| Dentate gyrus molecular layer | 41 | 10703 | 2 | 1 | 2 | 3 | 1 | 1 |  |
| Dentate gyrus polymorph layer | 42 | 10704 | 2 | 1 | 2 | 3 | 1 | 1 |  |
| Dentate gyrus granule cell layer | 43 | 632 | 2 | 1 | 2 | 3 | 1 | 1 |  |
| Septofimbrial nucleus | 57 | 310 | 2 | 1 | 3 | 3 | 2 | 1 |  |
| Central amygdalar nucleus | 59 | 536 | 2 | 1 | 1 | 3 | 1 | 1 |  |
| Triangular nucleus of septum | 63 | 581 | 2 | 1 | 2 | 3 | 2 | 1 |  |
| Ventral anterior-lateral complex of the thalamus | 64 | 629 | 2 | 4 | 3 | 1 | 3 | 1 |  |
| Lateral dorsal nucleus of thalamus | 76 | 155 | 2 | 3 | 1 | 2 | 5 | 2 |  |
| Perireunensis nucleus | 80 | 1077 | 2 | 6 | 1 | 4 | 3 | 1 |  |
| Reticular nucleus of the thalamus | 83 | 262 | 2 | 2 | 2 | 4 | 3 | 1 |  |
| Epithalamus | 85 | 958 | 2 | 6 | 4 | 2 | 3 | 2 |  |
| Zona incerta | 103 | 797 | 2 | 1 | 1 | 2 | 2 | 1 |  |
| thalamus related | 109 | 896 | 2 | 2 | 2 | 6 | 6 | 1 |  |
| fimbria | 116 | 603 | 2 | 1 | 2 | 1 | 3 | 1 |  |
| hippocampal commissures | 118 | 618 | 2 | 1 | 2 | 3 | 1 | 1 |  |
| epithalamus related | 122 | 1083 | 2 | 1 | 1 | 1 | 2 | 2 |  |
| Primary somatosensory area barrel field layer 6b | 9 | 1062 | 3 | 1 | 2 | 2 | 2 | 1 |  |
| Dorsal auditory area layer 6b | 22 | 243 | 3 | 6 | 4 | 1 | 3 | 3 |  |
| Ventral auditory area | 23 | 1018 | 3 | 1 | 4 | 2 | 3 | 1 |  |
| Anterior cingulate area dorsal part | 24 | 39 | 3 | 1 | 1 | 3 | 2 | 4 |  |
| Field CA1 | 38 | 382 | 3 | 1 | 1 | 1 | 2 | 2 |  |
| Basolateral amygdalar nucleus | 51 | 295 | 3 | 1 | 3 | 1 | 2 | 2 |  |
| Posterior amygdalar nucleus | 53 | 780 | 3 | 2 | 1 | 4 | 1 | 2 |  |
| Pallidum ventral region | 62 | 835 | 3 | 2 | 1 | 6 | 1 | 2 |  |
| Intermediodorsal nucleus of the thalamus | 77 | 59 | 3 | 5 | 5 | 2 | 5 | 2 |  |
| Intralaminar nuclei of the dorsal thalamus | 82 | 51 | 3 | 5 | 2 | 2 | 4 | 2 |  |
| Dorsomedial nucleus of the hypothalamus | 89 | 830 | 3 | 1 | 1 | 1 | 2 | 1 |  |
| Subparaventricular zone | 90 | 347 | 3 | 4 | 2 | 4 | 6 | 2 |  |
| Medial preoptic nucleus | 94 | 515 | 3 | 2 | 1 | 2 | 2 | 1 |  |
| Paraventricular hypothalamic nucleus descending division | 96 | 63 | 3 | 2 | 1 | 4 | 3 | 1 |  |
| Lateral hypothalamic area | 98 | 194 | 3 | 4 | 1 | 4 | 2 | 1 |  |
| supraoptic commissures | 120 | 349 | 3 | 5 | 1 | 5 | 4 | 1 |  |
| Ventral medial nucleus of the thalamus | 65 | 685 | 4 | 3 | 3 | 2 | 3 | 2 |  |
| Subfornical organ | 91 | 338 | 4 | 6 | 3 | 3 | 3 | 3 |  |
| mammillary related | 121 | 46 | 4 | 1 | 1 | 2 | 1 | 1 |  |

